# Leveraging the local genetic structure for trans-ancestry association mapping

**DOI:** 10.1101/2022.03.26.485910

**Authors:** Jiashun Xiao, Mingxuan Cai, Xinyi Yu, Xianghong Hu, Xiang Wan, Gang Chen, Can Yang

## Abstract

Over the past two decades, genome-wide association studies (GWASs) have successfully advanced our understanding of genetic basis of complex traits. Despite the fruitful discovery of GWASs, most GWAS samples are collected from European populations, and these GWASs are often criticized for their lack of ancestry diversity. Trans-ancestry association mapping (TRAM) offers an exciting opportunity to fill the gap of disparities in genetic studies between non-Europeans and Europeans. Here we propose a statistical method, LOG-TRAM, to leverage the <u>lo</u>cal genetic architecture for TRAM. By using biobank-scale datasets, we showed that LOG-TRAM can greatly improve the statistical power of identifying risk variants in under-represented populations while producing well-calibrated *p*-values. We applied LOG-TRAM to the GWAS summary statistics of 29 complex traits/diseases from Biobank Japan (BBJ) and UK Biobank (UKBB), and achieved substantial gains in power (the effective sample sizes increased by 49% in average compared to the BBJ GWASs) and effective correction of confounding biases compared to existing methods. Finally, we demonstrated that LOG-TRAM can be successfully applied to identify ancestry-specific loci and the LOG-TRAM output can be further used for construction of more accurate polygenic risk scores (PRSs) in under-represented populations.

## Introduction

Thousands of genome-wide association studies (GWASs) have been conducted to understand the genetic basis of complex human traits/diseases since the first GWAS publication in 2005 [1]. As of January 2022, more than 326,000 genome-wide significant associations (*p*-value ≤ 5 × 10^−8^) have been identified between single nucleotide polymorphisms (SNPs) and complex human traits [2]. The summary statistics from GWASs are accessible through public gateways, such as the GWAS Catalog [2]. These datasets contain rich information on genetic variations and phenotypes, providing an unprecedented research opportunity in biomedical and social science [3].

Despite the great achievements, GWAS findings are limited by the lack of ancestral diversity. According to the GWAS Diversity Monitor (https://gwasdiversitymonitor.com), about 88.9% of GWAS participants have been of European ancestry to date [4]. Non-European populations are severely under-represented in GWASs. For example, the proportions of participants of East Asian (EAS) and African ancestries are less than 6.9% and 0.4%, respectively [4]. Most of GWAS findings are based on individuals of European ancestry. Unfortunately, these findings may not be directly extrapolated to non-European ancestries [5]. As genetic studies with diversified ancestries are important in achieving global health equity, trans-ancestry association mapping (TRAM), which aims to identify risk genetic variants across ancestries (particularly in the under-represented populations), has become a critical step toward precision medicine [5, 6].

The challenges of TRAM arise from two major aspects. First, the genetic architectures of a phenotype are heterogeneous across ancestries [5]. Some trait-associated SNPs have vastly different allele frequencies between European and non-European ancestries [5, 7]. SNP effect sizes and linkage disequilibrium (LD) patterns can also vary across ancestries [8]. Second, the publicly released GWAS summary statistics still suffer from confounding biases [9]. Although principal component analysis (PCA) [10] and linear mixed models (LMMs) [11] are commonly used for association mapping in GWASs, the population stratification in biobank-scale data, such as socioeconomic status [12] or geographic structure [13], may not be fully accounted for in these standard approaches. Without correcting confounding biases hidden in GWAS summary statistics, TRAM will produce many false positive findings.

Much effort has been devoted to develop statistical methods for TRAM. To handle heterogeneity of genetic architectures, the random-effects (RE) model naturally becomes a popular tool for meta-analysis of GWASs [14, 15, 16]. Despite its popularity, the RE model assumes that effect sizes vary greatly under the null hypothesis. This assumption can be fairly conservative [17]. To improve the power of RE models for TRAM, a new RE method named RE2 [17] has been developed by assuming no heterogeneity under the null hypothesis. Later, these authors further improved the RE2 method by allowing meta-analysis of correlated statistics, namely, RE2C [18]. Unlike the general hypothesis testing methods, MANTRA was specifically designed for meta-analysis of trans-ancestry GWAS data [19]. MANTRA assumes that the effect sizes should be closely matched for similar genetic ancestries; however, it allows for heterogeneity in effect sizes for more distal ancestries. MTAG [20] is a recently developed method to analyze multiple GWASs from a single ancestry. MTAG can greatly improve the statistical power of association mapping because it uses the global genetic correlation to borrow information across related traits. It assumes that the variance and covariance of marginal effect sizes are homogeneous across SNPs. This assumption may limit its usage in the trans-ancestry setting. Very recently, MAMA [21] has been developed to improve MTAG for TRAM by accounting for heterogeneity in LD across ancestries. However, both MTAG and MAMA are still not fully satisfactory when the local genetic architecture differs from the global architecture.

In this work, we develop a statistical method to leverage the <u>lo</u>cal genetic architecture for <u>tr</u>ans-ancestry <u>a</u>ssociation <u>m</u>apping (LOG-TRAM). Not only can LOG-TRAM control type I errors at the nominal level, it can also improve the statistical power of identifying risk variants in under-represented populations. Below are the keys to the success of LOG-TRAM. First, it can greatly improve the statistical power of association mapping by making use of biobank-scale datasets of the auxiliary population (e.g., the UK Biobank dataset (UKBB)) while accounting for heterogeneity among multiple ancestries. Second, compared to existing meta-analysis methods that consider the global genetic architecture, LOG-TRAM focuses on the local genetic architectures to localize risk variants, including local heritability, local co-heritability, allele frequencies, SNP effect sizes, and LD patterns. Third, it is capable of correcting the confounding bias hidden in GWAS summary statistics to avoid inflated type I errors. Fourth, LOG-TRAM only takes summary statistics from multiple ancestries as inputs and outputs well-calibrated *p*-values. With the innovations of our model design, LOG-TRAM is computationally efficient as it has a closed-form solution at each step. Through comprehensive simulation studies, we demonstrated that LOG-TRAM largely outperformed existing meta-analysis approaches in terms of type I error rate and power. Then we applied LOG-TRAM to the GWAS summary statistics of 29 complex traits and diseases from BioBank Japan (BBJ) and UKBB. The analysis results show that LOG-TRAM can effectively account for confounding biases in the GWAS summary statistics and achieve a substantial gain in power for identification of risk variants. Furthermore, we successfully applied LOG-TRAM to identify ancestry-specific loci. Finally, we showed that the LOG-TRAM output can be used for the construction of more accurate polygenic risk scores (PRSs) in under-represented populations.

## Material and Methods

### Notation and problem setup

To introduce LOG-TRAM, we begin our formulation with the individual-level GWAS data. Without loss of generality, we consider two populations and the following model that relates genotypes with phenotypes in populations 1 and 2, respectively,

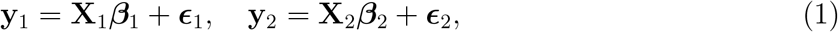

where 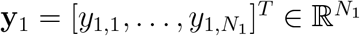 and 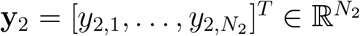 are two phenotype vectors, 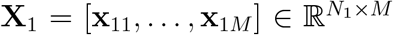 and 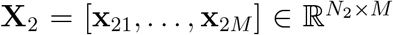 are standardized genotype matrices whose columns have zero mean and unit variance, ***β***_1_ = [*β*_11_, *β*_12_, …, *β*_1*M*_]^*T*^ ∈ ℝ^*M*^ and ***β***_2_ = [*β*_21_, *β*_22_, …, *β*_2*M*_]^*T*^ ∈ ℝ^*M*^ are the SNP effect sizes, the residual vectors 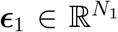 and 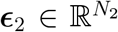 are independent error terms, *M* is the number of SNPs, *N*_1_ and *N*_2_ are the sample sizes of populations one and two, respectively. We consider population one as the under-represented target population and population two as the auxiliary population with a biobank-scale sample size, i.e., *N*_1_ ≪ *N*_2_. We expect that information from population two can be useful for association mapping in population one. Here we assume that the covariates (e.g., age, sex and principal components) have been properly adjusted. More detailed treatment on covariate adjustment can be found in our previous work [22] and other related work [23]. We also implicitly assume that SNP effect sizes increase as the allele frequencies decrease when working with standardized genotype matrices [22].

Suppose that individual-level data {**y**_1_, **X**_1_, **y**_2_, **X**_2_} are not accessible but summary statistics 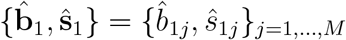 and 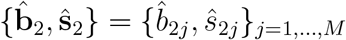 from the simple linear regressions are available:

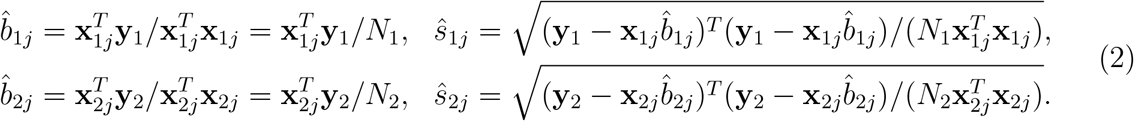

Traditional association methods [24] often report genome-wide significant association status based on the *z*-scores 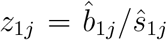 and 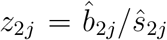. However, these association methods have several limitations. First, they do not account for the heterogeneity of LD patterns across ancestries. Second, they do not correct possible confounding factors hidden in summary statistics 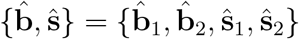, resulting in inflated type I errors. Third, they do not fully utilize the correlation between ***β***_1_ and ***β***_2_, leading to a suboptimal statistical power.

Recent meta-analysis studies have revealed that statistical power of association mapping in under-represented populations can be improved by leveraging global cross-population genetic correlation [19, 21]. However, current trans-ancestry GWAS meta-analysis relies on the assumption that variance and covariance of SNP effect sizes are homogeneous across the entire genome [20, 21]. As a matter of fact, accumulating evidence suggests different genomic regions contribute disproportionately to the heritability across the genome, inducing heterogeneous genetic similarity between populations in different local regions [25, 26, 27, 28, 29]. Therefore, it is more appropriate to localize risk variants by leveraging the local genetic architecture rather than the global architecture. To see this more intuitively, let us consider the global genetic correlation *r*_*g*_ = *Corr*(*β*_1*j*_, *β*_2*j*_) averaged across the entire genome *j* ∈ {1, …, *M*} and the local genetic correlation *r*_*g*,***ℛ***_ = *Corr*(*β*_1*j*_, *β*_2*j*_) defined on a local genomic region *j* ∈ ***ℛ***, where the local genetic correlation *r*_*g*,***ℛ***_ can be very different from the global *r*_*g*_ [26, 30, 31, 27]. On the one hand, *r*_*g*,***ℛ***_ can be nearly zero even if the global correlation is substantial, e.g., *r*_*g*_ = 0.7. Clearly, no information should be borrowed from the auxiliary population for association mapping of the target population in the local region ***ℛ***. In such a case, leveraging the global genetic correlation (*r*_*g*_ = 0.7) for TRAM leads to inflated type I errors. On the other hand, when using the global genetic correlation for TRAM, we lose statistical power substantially in the presence of strong local genetic correlation *r*_*g*,***ℛ***_ = 0.7 but weak global genetic correlation (e.g., *r*_*g*_ ≈ 0).

To leverage the local genetic architecture for TRAM, we propose a statistical method named LOG-TRAM which only requires GWAS summary statistics 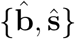 and a reference genome (e.g., The 1000 Genomes Project) as inputs (Fig. 1). The output of LOG-TRAM 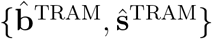 has the following desired properties: (i) by borrowing information from biobank-scale European datasets through the local genetic architecture, the statistical power of non-EUR association studies can be largely improved; (ii) the influence of confounding factors can be adjusted; (iii) the LOG-TRAM estimate is unbiased and its standard error can be much smaller than that of standard GWASs (see Appendix D); and (iv) the LOG-TRAM output can be used for downstream analysis, such as construction of more accurate polygenic risk scores in the under-represented populations.

**Figure 1:**
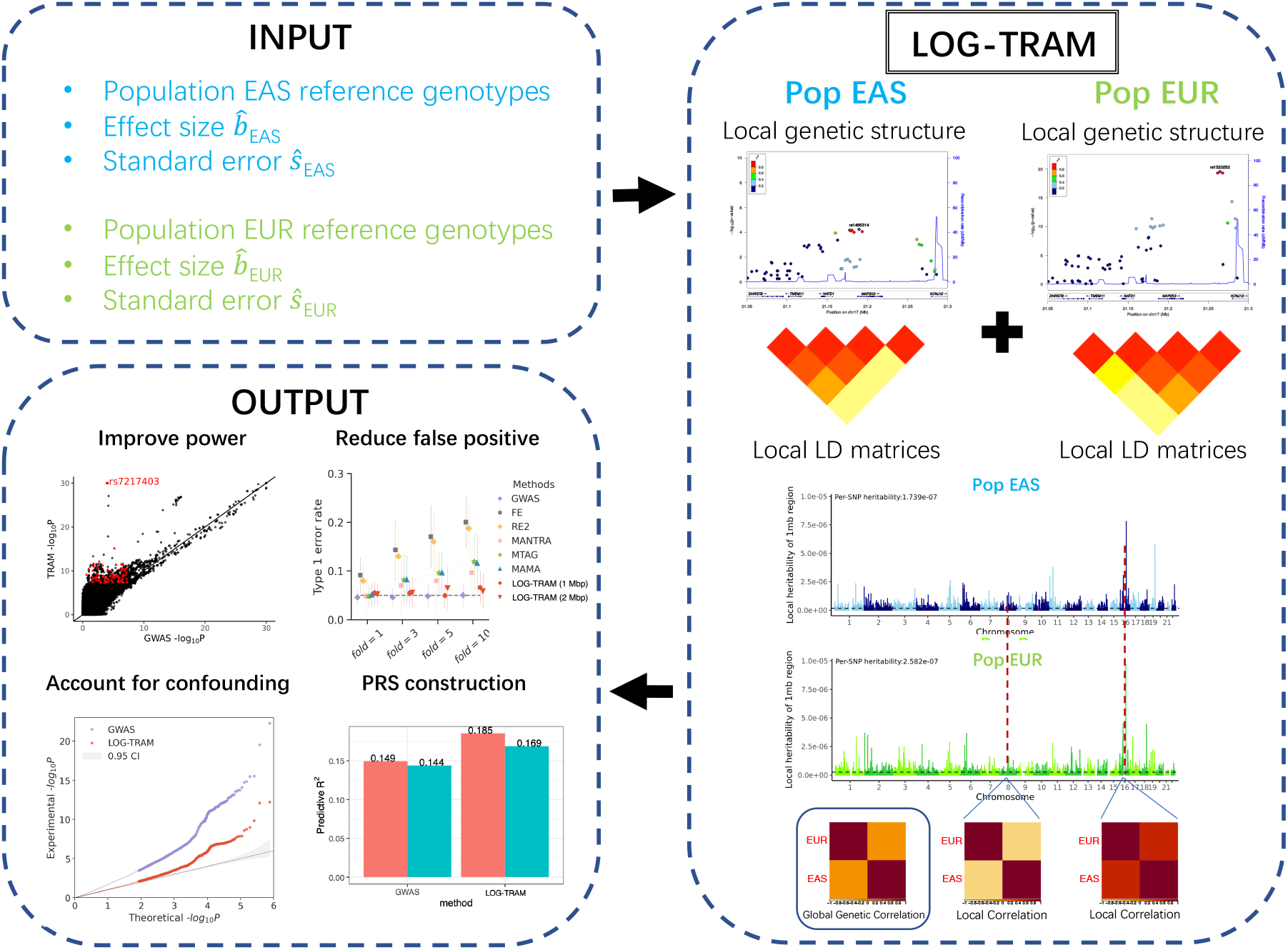
LOG-TRAM overview. LOG-TRAM only requires GWAS summary statistics 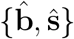 and reference genomes of the considered populations as inputs. During the trans-ancestry meta-analysis, LOG-TRAM can handle the heterogeneity of genetic architectures (e.g., allele frequencies, LD patterns, and SNP effect sizes) across populations, and leverage information from biobank-scale European datasets through the local genetic architecture to increase statistical power and simultaneously reduce false positives for non-EUR association studies.

### The LOG-TRAM model

To leverage the local genetic architecture, we first partition the genome into consecutive non-overlapping regions (e.g., 1 M base-pair segments). Then we focus on identification of risk variants within a given region ***ℛ*** at a time. Given the local region ***ℛ***, we extend model (1) as:

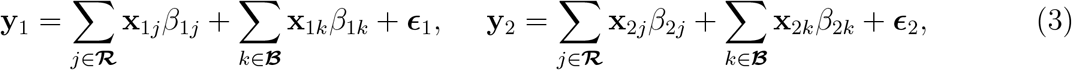

where ***ℬ*** = {1, …, *M*} \ ***ℛ*** is the set of all SNPs in the genome except the local region ***ℛ, ϵ***_1_ and ***ϵ***_2_ are vectors of independent noise with 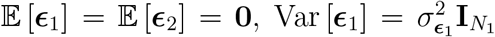 and 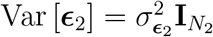. Partitioning the genome into the local region ***ℛ*** and the background region ***ℬ*** allows us to model the regional genetic architecture differently from the global pattern. To achieve this, we introduce the following probabilistic structure for ***β***_1_ and ***β***_2_:

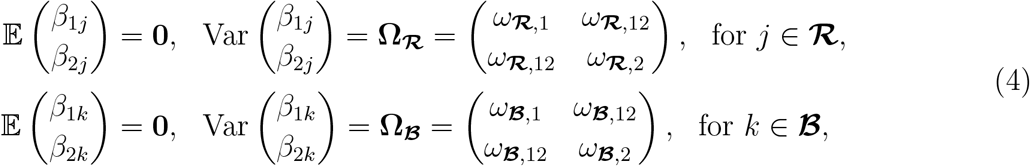

where **Ω**_***ℛ***_ captures the genetic covariance of two populations in the local region ***ℛ*** with its diagonal elements *ω*_***ℛ***,1_ and *ω*_***ℛ***,2_ representing the local per-SNP heritability for populations one and two, respectively; and the off-diagonal element *ω*_***ℛ***,12_ representing the local per-SNP co-heritability between population one and population two, **Ω**_***ℬ***_ is the covariance matrix of the genetic effects from the background region ***ℬ*** (the entire genome except ***ℛ***) with *ω*_***ℬ***,1_ and *ω*_***ℬ***,2_ representing the background per-SNP heritability from populations one and two, respectively; and *ω*_***ℛ***,12_ representing the background per-SNP co-heritability between population one and population two. Without loss of generality, we assume that the phenotypes have been standardized, i.e., Var(*y*_1,*i*_) = Var(*y*_2,*i*_) = 1. Based on models (3) and (4), we have 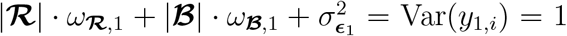 and 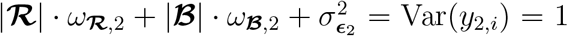, where |***ℛ***| and |***ℬ***| represent the number of SNPs in ***ℛ*** and ***ℬ***, respectively.

The flexible statistical structure given in Eq. (4) has two salient properties. First, it models the genetic covariance between the two populations (*ω*_***ℛ***,12_ and *ω*_***ℬ***,12_), allowing us to borrow information from biobank-scale datasets. Second, rather than assuming that the genetic covariance of SNP effect sizes are homogeneous across the entire genome, i.e., **Ω**_***ℛ***_ = **Ω**_***ℬ***_, we allow the per-SNP heritability of the local genomic region (*ω*_***ℛ***,1_ and *ω*_***ℛ***,2_) to be different from the background pattern (*ω*_***ℬ***,1_ and *ω*_***ℬ***,2_) and the local per-SNP co-heritability (*ω*_***ℛ***,12_) to be different from the global per-SNP co-heritability (*ω*_***ℬ***,12_), effectively capturing the heterogeneous local structures.

So far, we have described the method without covariates. In the presence of covariates such as gender, age, and principal component scores, we extend model (3) as:

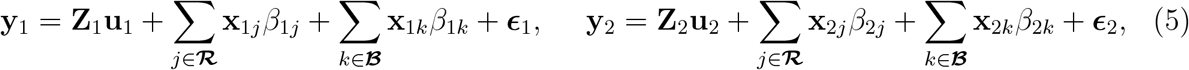

where 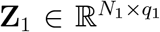 and 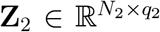 are the covariate matrices of the target and auxiliary population, respectively, **u**_1_ and **u**_2_ are the corresponding vectors of covariates effects. To get ride of the covariates, we define the projection matrices 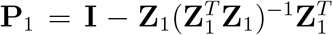 and 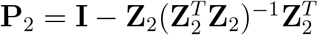 and obtain the new working model as:

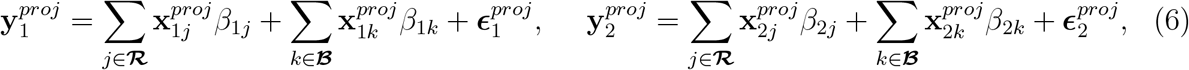

where 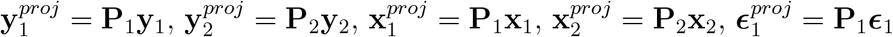, and 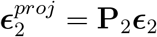. By the projection of covariates, model (5) reduces to model (3). In genomics, this projection approach is commonly used for association mapping [32] and heritability estimation [33]. In the main text, without loss of generality, we work on model (3) in the absence of covariates.

### The LOG-TRAM model with summary-level data

The individual-level GWAS data are often not easily accessible due to privacy concerns. To overcome this difficulty, we consider the rows of **X**_1_ and **X**_2_ as independent and identically distributed samples from the target population and auxiliary population, respectively. As such, we define the correlation between SNP *j* and SNP *k* as 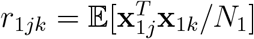 and 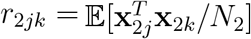 for target and auxiliary populations, respectively. We can then define the underlying true marginal effect sizes as:

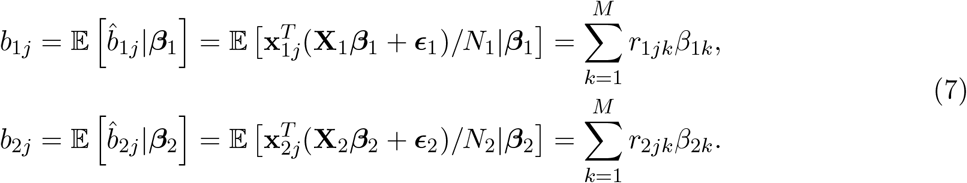

As we can observe, the true marginal effect size *b*_1*j*_ is the summation of underlying true effect sizes *β*_1*k*_ in LD with SNP *j* weighted by the pairwise SNP correlation between SNPs *j* and *k*. With the above model specification, we can show that the following relationship holds for the obtained summary statistics 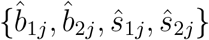 for SNP *j* in the local region ***ℛ***,

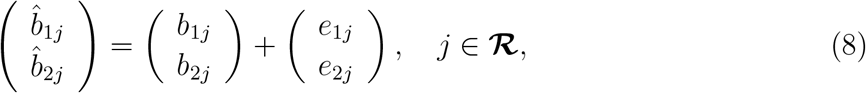

where *e*_1*j*_ and *e*_2*j*_ are the independent estimation errors with 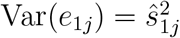 and 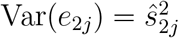. Because the two datasets are from different populations, we have Cov(*e*_1*j*_, *e*_2*j*_) = 0. More specifically, we can derive the relationship (see Appendix A):

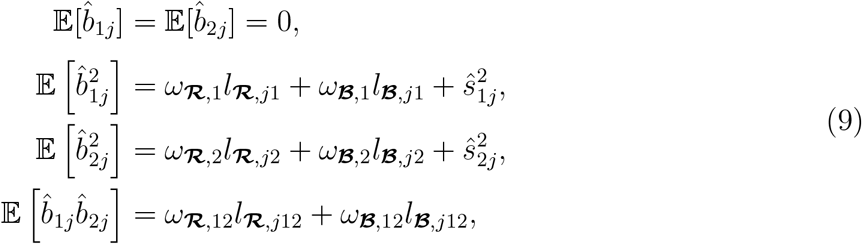

where 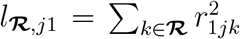 and 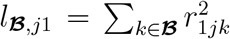 are the LD scores of SNP *j* in population one for regions ***ℛ*** and ***ℬ***, respectively, 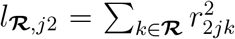 and 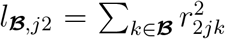 are the LD scores of SNP *j* in population two for regions ***ℛ*** and ***ℬ***, respectively. Please be noted that *l*_***ℛ***,*j*12_ = ∑_*k*∈***ℛ***_ *r*_1*jk*_*r*_2*jk*_ and *l*_***ℬ***,*j*12_ = ∑_*k*∈***Bℬ***_ *r*_1*jk*_*r*_2*jk*_ are the cross population LD scores of SNP *j* for regions ***ℛ*** and ***ℬ***, respectively. As LD decays with distance, Eq. (9) implies that the obtained summary statistics largely depend on the local genetic architecture, including the local LD scores (*l*_***ℛ***,*j*1_, *l*_***ℛ***,*j*2_ and *l*_***ℛ***,*j*12_), local per-SNP heritability (*ω*_***ℛ***,1_ and *ω*_***ℛ***,2_) and local per-SNP co-heritability (*ω*_***ℛ***,12_). Combining Eq. (8) and Eq. (9), we can further derive the relationship between the summary statistics and the model parameters **Ω**_***ℛ***_ and **Ω**_***ℬ***_,

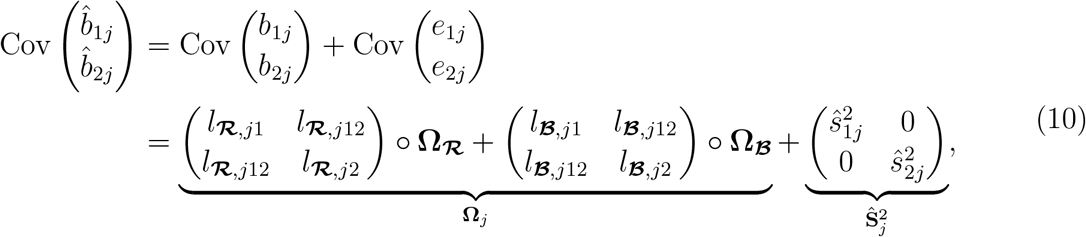

where ∘ is the element-wise product.

To account for the hidden confounding biases in GWAS summary statistics 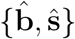, we generalize linear model (3) based on the genetic drift model used in LDSC [9] and modify Eq. (9) as (See Appendix B):

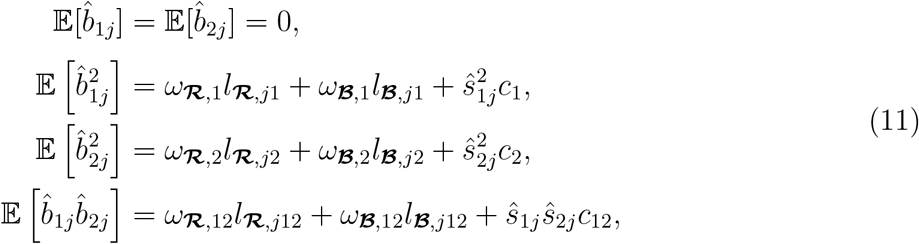

where *c*_1_, *c*_2_ and *c*_12_ are inflation constants which adjust for the confounding biases of GWAS standard errors. From Eq. (11), we can see that the influence of population stratification on the variance of marginal estimates remains nearly constant across SNPs (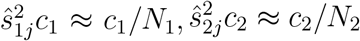 and 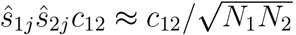), while the magnitudes of genetic effects are tagged by LD scores. Therefore, the Eq. (10) can be updated as:

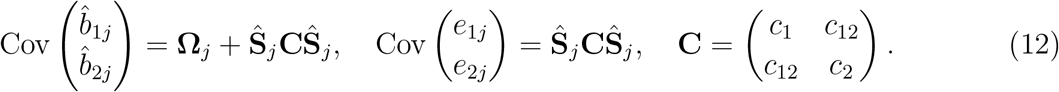

In real data analysis, we find that inflation constants *c*_1_ and *c*_2_ are often larger than one, and *c*_12_ is very close to zero because of no overlapped samples in trans-ancestry association studies.

### The LOG-TRAM estimator

We use the generalized method of moments (GMM) [21] to derive the LOG-TRAM estimator 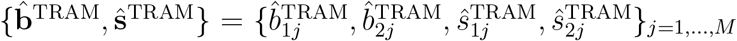. Using model (12), we first obtain the conditional mean and conditional variance (see Appendix C) as

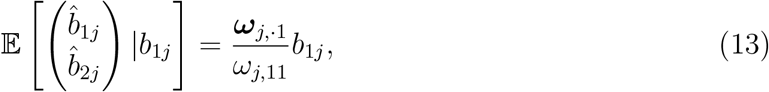

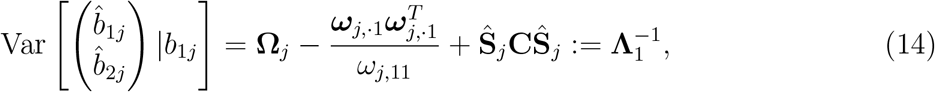

where ***ω***_*j*,·1_ = [***ω***_*j*,11_, ***ω***_*j*,12_]^*T*^ is the first column of **Ω**_*j*_. Based on Eq. (13), we can define 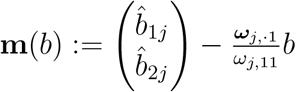 and give the first-order moment condition

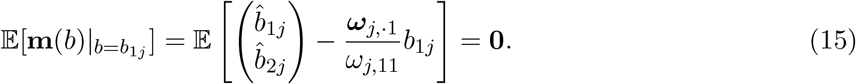

Based on GMM, we can obtain the LOG-TRAM estimator as

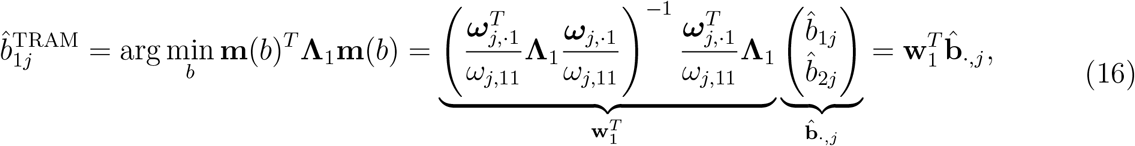

and its variance as

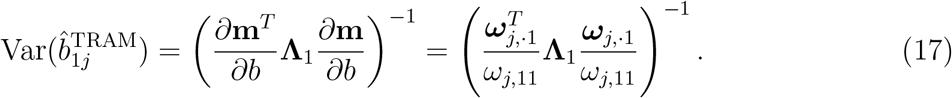

Similarly, we can obtain 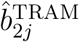 and its variance 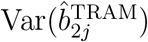. Eq. (16) implies that the LOG-TRAM estimator incorporates three parts of information. First, the LD heterogeneity between populations is properly adjusted by the cross-population LD scores *l*_***ℛ***,12_ and *l*_***ℬ***,12_ when constructing **Ω**_*j*_. When the LD structure is very different, the cross-population LD scores become smaller, which down-weighs the genetic covariance. Second, by introducing the local genetic covariance **Ω**_***ℛ***_, LOG-TRAM is able to utilize the local genetic architecture to improve association mapping. Third, LOG-TRAM corrects confounding biases from the GWAS data through the term **Ŝ**_*j*_ **CŜ**_*j*_ and thus produces well calibrated statistics.

### Parameter estimation and statistical inference

To obtain unknown parameters ***θ*** = {*ω*_***ℛ***,1_, *ω*_***ℛ***,2_, *ω*_***ℛ***,12_, *ω*_***ℬ***,1_, *ω*_***ℬ***,2_, *ω*_***ℬ***,12_, *c*_1_, *c*_2_, *c*_12_} in **Ω**_*j*_ and **C** of Eq. (16), we use Eq. (12) to regress the squared GWAS marginal effects 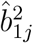 on the LD scores *l*_***ℛ***,*j*1_ and *l*_***ℬ***,*j*1_ [9]:

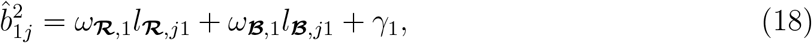

where *γ*_1_ is the intercept. Then the estimates 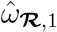 and 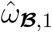 can be obtained from the regression coefficients and the estimate of *c*_1_ is given as 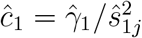. Following the LDSC [9], we exclude SNPs with *χ*^2^ *>* 30 to remove outliers that deviate from the linearity assumption. Similarly, 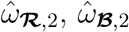 and *ĉ*_2_ can be obtained from the coefficients of regressing 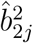 on the LD scores *l*_***ℛ***,*j*2_ and *l*_***ℬ***,*j*2_. The cross-population parameters 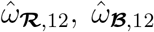 and *ĉ*_12_ can be obtained from the coefficients of regressing the product 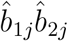 on the cross-population LD scores *l*_***ℛ***,*j*12_ and *l*_***ℬ***,*j*12_ [34]. As the LD scores are pre-calculated from the reference genome (e.g., The 1000 Genomes Project), the parameter estimation can be performed efficiently.

Plugging in the estimated parameters 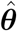, we can construct the estimates of **Ω**_*j*_ and **C** as:

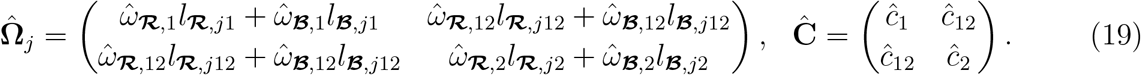

We then obtain the LOG-TRAM estimates 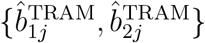 from Eq. (16) and the corresponding variance 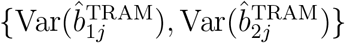 from Eq. (17). The standard errors are computed by 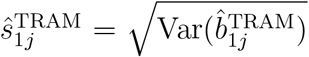 and 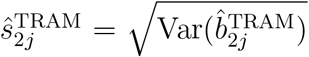. With the above LOG-TRAM outputs, we can detect nonzero SNP effect sizes (*b*_1*j*_ ≠ 0 and *b*_2*j*_ ≠ 0) based on the Wald test.

### Identification of variants with ancestry-specific effects by LOG-TRAM

Not only can LOG-TRAM provide inference on the SNP effect sizes from the two populations, but it also offers a statistical test to identify variants with ancestry-specific effects. Specifically, we denote the difference of the SNP effects size as *δ*_*j*_ = *b*_1*j*_ − *b*_2*j*_, where *b*_1*j*_ and *b*_2*j*_ are the underlying true marginal effect sizes defined in Eq. (7). The estimate of *δ*_*j*_ can be obtained from the LOG-TRAM output:

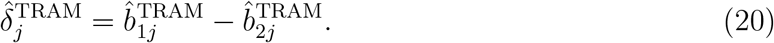

The variance of 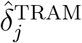 is given as

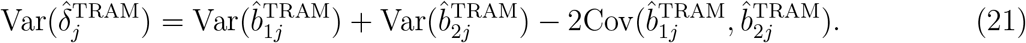

Here the covariance term is derived as

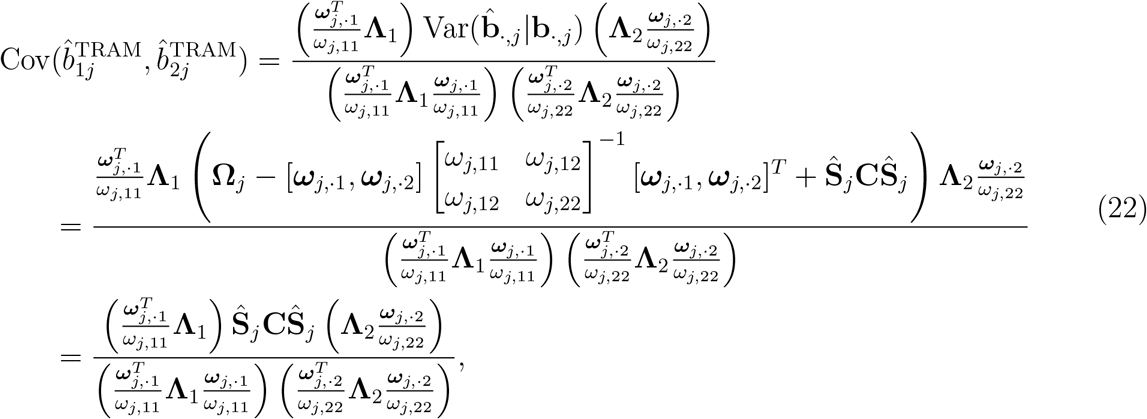

where **Λ**_2_ is defined as 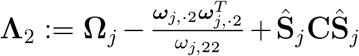. With Eq. (20) and Eq. (21), we can obtain the LOG-TRAM-based difference test statistic 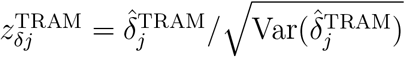 and apply the Wald test to test hypothesis H_0_ : *δ*_*j*_ = 0 v.s. H_1_ : *δ*_*j*_ ≠ 0.

Recall that the traditional methods simply obtain the estimate of *δ*_*j*_ and its variance based on the original GWAS estimate: 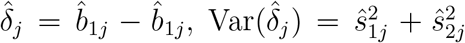, and then derive the GWAS-based test statistic 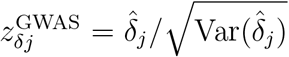. As LOG-TRAM can borrow information across populations and account for confounding bias, it can substantially improve the statistical power of identifying variants with ancestry-specific effects.

## Results

### Simulation study

We conducted comprehensive simulation studies to compare the performance of LOG-TRAM with several commonly used GWAS meta-analysis methods, including fixed-effect IVW meta-analysis (e.g., FE [35]), random-effects model (e.g., RE2 [17, 18]), MANTRA[19], MTAG [20], and MAMA [21]. FE assumes that a SNP has the same marginal effect across populations. Thus, it is computationally efficient but its type I error rate is not well controlled. RE2 and MANTRA account for the heterogeneity of marginal effects across populations while ignoring the LD difference. MTAG is developed for modeling the heterogeneity of marginal effect sizes across GWAS summary statistics of multi-traits in a single population. Recently, MAMA extends MTAG to handle multiple ancestries and takes the heterogeneity among populations (e.g., LD, minor allele frequency) into account. However, both MTAG and MAMA rely on the assumption that variance and covariance of SNP effect sizes are homogeneous across the whole genome.

We first conducted the simulations to evaluate the type I error rate. We considered 18K samples from the EAS cohort [22] and 100K samples from UKBB as the target and auxiliary populations, respectively. We used their genotype matrices to mimic the different LD patterns and allele frequencies between populations. To investigate the type I error rate, 17,248 HapMap3 matched SNPs from chromosome 20 were used in our simulation study. We partitioned chromosome 20 into two segments at base pair position 3,119,133 (GRCh37). One segment taking up about 95% of chromosome 20 was considered as the background region ***ℬ***. We generated a polygenic scenario by setting 10% SNPs with shared non-zero effects and varied the trans-ancestry genetic correlation (denoted as *r*_*g*_) among {0, 0.2, 0.4, 0.6}. The shared effects were simulated from the bivariate normal distribution 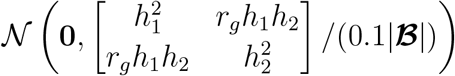, where 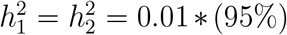 means that the total heritability contributed by chromosome 20 is around 0.01, and |***ℬ***| is number of SNPs in the background region ***ℬ***. Another segment taking up 5% of chromosome 20 was treated as the local region ***ℛ***. We then used the SNPs in ***ℛ*** to assess the type I error rate of compared methods. Specifically, we set *β*_1*j*_ = 0 for *j* ∈ ***ℛ*** for EAS while simulating the true effects of EUR (*β*_2*j*_, *j* ∈ ***ℛ***) by a normal distribution 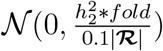, where 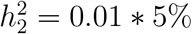, |***ℛ***| is number of SNPs in the local region ***ℛ***, and 0.1 represents that 10% of SNPs jointly contribute to heritability, 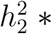 *fold*, with per-SNP heritability 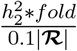. The parameter *fold* allows us to vary the enrichment fold of local heritability at 1,3,5, and 10, such that the local genetic architecture differs from the global one. Given the effect sizes and genotype matrices, quantitative phenotypes in both populations were simulated using Eq. (3). After that, we marginally regressed the simulated phenotypes on each SNP to obtain the *z*-scores of the two datasets. Finally, after performing meta-analysis, we reported the fraction of SNPs with a *p*-value less than 0.05 in the local region ***ℛ*** of EAS as the type I error rate.

As expected, Fig. 2A shows that only the single-ancestry-based GWAS and LOG-TRAM have well-controlled type I error rates regardless of the enrichment fold of local heritability for region ***ℛ*** in EUR. In contrast, the type I error rates of other meta-analysis methods with the homogeneous assumption (e.g., FE, RE2, MANTRA, MTAG, MAMA) increase as the enrichment fold increases from 1 to 10. More specifically, FE and RE2 approaches performed worst in most cases. When the background trans-ancestry genetic correlation (*r*_*g*_) is non-zero, MANTRA, MTAG and MAMA suffered from severely inflated type I error rates as they misused or overused information from large-scale EUR GWAS summary statistics. We also examined whether the performance of LOG-TRAM could be sensitive to the size of window (or local genomic region). Specifically, we considered two window sizes, 1M and 2M base pairs, to partition the whole chromosome into multiple non-overlapping local regions. We observed that the type I error rates of LOG-TRAM were well controlled despite the different window sizes. Therefore, we set a window size of 1M base pairs as the default setting. In summary, LOG-TRAM has a satisfactory control of the type I error rate and its performance is insensitive to the size of a local region. Because FE and RE2 showed severe inflation of the type I error rate, we did not include them in the power comparison.

**Figure 2:**
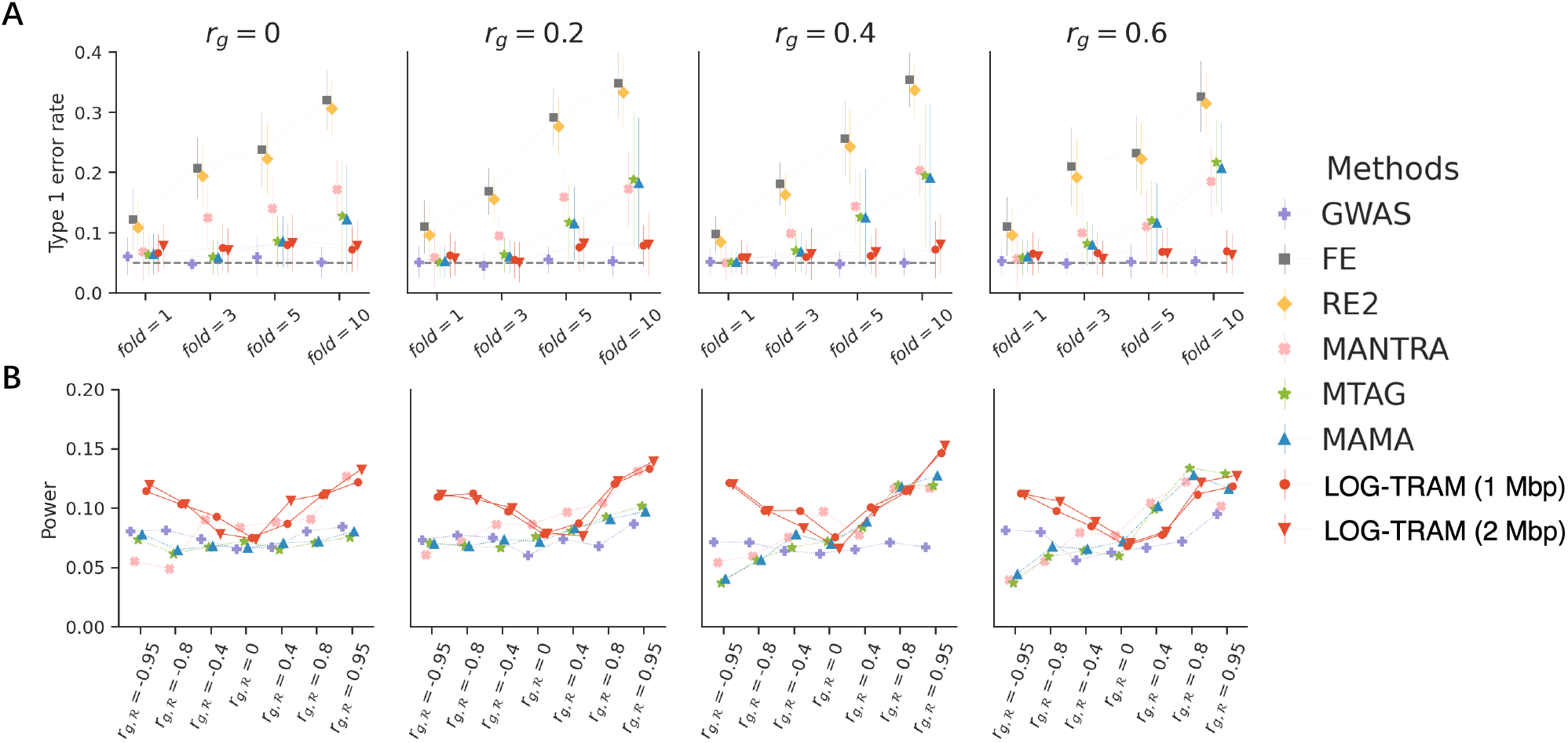
Comparisons among LOG-TRAM, MAMA, MTAG, MANTRA, RE2, FE, and single-ancestry GWAS in simulation studies. **A**: Average type I error rates assessed under different settings of background trans-ancestry genetic correlations (*r*_*g*_) and the fold enrichment (*fold*) of EUR local heritability. Error bars represent the standard errors of type I error rates evaluated on 20 replications. **B**: Average statistical power in multiple simulation scenarios with different combinations of background trans-ancestry genetic correlations (*r*_*g*_) and local trans-ancestry genetic correlations (*r*_*g*,***ℛ***_). Results are also summarized from 20 replications. More simulation results are provided in Fig. S1-S8.

Next we evaluated the power of compared methods in the cross-population setting. Different from the null SNP setting in the local region ***ℛ*** of EAS for the evaluation of type I error rates, we generated the SNP effects of the twolpopulation s in local region ***ℛ*** by a bivariate normal distribution 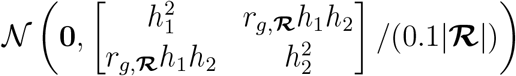, where *r*_*g*,***ℛ***_ is the local trans-ancestry genetic correlation of region ***ℛ***, and 0.1 represents that 10% of SNPs with non-zero effects jointly contribute to the local heritability. We considered a wide spectrum of local correlations to mimic the heterogeneous genetic similarity between populations in different gnomic regions: −0.95, −0.8, −0.4, 0, 0.4, 0.8, 0.95. We anticipated that LOG-TRAM would achieve substantial power gain in identifying SNPs with non-zero effects when the local genetic correlation was strong and the global genetic correlation was weak. As we described in the above, we simulated the quantitative phenotypes and obtained the *z*-scores of the two datasets. Finally, after performing meta-analysis, we reported the fraction of non-zero SNPs in the EAS local region ***ℛ*** with a *p*-value less than 0.05 as the power. As shown in Fig. 2B, LOG-TRAM achieved the best performance in all settings. When the global/background trans-ancestry genetic correlation *r*_*g*_ is 0, LOG-TRAM was still able to increase the power by utilizing the information from EUR datasets through non-zero local genetic correlation *r*_*g*,***ℛ***_. By contrast, MAMA and MTAG reduced to the standard single-ancestry GWAS since no information could be borrowed from the large-scale EUR datasets. When the global genetic correlation *r*_*g*_ became close to *r*_*g*,***ℛ***_, MAMA and MTAG achieved comparable performance with LOG-TRAM.

For identification of ancestry-specific loci, we compared the performance of GWAS-based and LOG-TRAM-based difference test in simulation studies. As shown in Fig. S14, the LOG-TRAM-based difference test has significant higher power than the GWAS-based test while achieving well-controlled type I error rates. In summary, simulation studies suggest that LOG-TRAM has a great advantage over other methods by leveraging the local genetic architecture.

### Real data analysis

#### Application of LOG-TRAM for trans-ancestry association mapping

To evaluate the performance of LOG-TRAM in real applications, we applied LOG-TRAM to publicly available GWAS summary statistics of 29 phenotypes from EAS and EUR. The details of datasets are given in Table S1. For each GWAS dataset, we used SNPs that overlapped the HapMap 3 list, and removed the SNPs with ambiguous alleles. For a pair of phenotypes from different populations, we aligned the sign of their effect sizes to the same allele. The LD scores were estimated with a sliding window of 1 M base-pair in the genome using 417 EUR and 377 EAS individuals from the 1000 Genomes Project. Fig. 3A shows a summary of results for all analyzed phenotypes in the EAS population. Among all 29 traits, we observed that LOG-TRAM consistently identified more independent loci than the standard single-ancestry GWASs. In summary, LOG-TRAM identified 1,954 lead SNPs in EAS, among which 842 were novel (i.e., not reported by the standard GWASs of BBJ). Besides, for each trait, we estimated the augmented effective sample size using the increase in mean *χ*^2^ statistics of LOG-TRAM relative to GWASs. Specifically, we assessed how large the sample size were needed to attain an equivalent increase in the mean *χ*^2^ statistics of LOG-TRAM. Overall, we observed that the original GWAS sample size had to be increased by 49% on average to achieve an equivalent power gained by LOG-TRAM (Fig. 3B and Table S2). As an example, compared to the 22 independent loci identified in the original GWAS of Systolic blood pressure (SBP) from BBJ, LOG-TRAM achieved a substantially higher power for EAS associations by identifying (69 − 22)*/*22 ≈ 214% more significant loci. LOG-TRAM increased the effective sample size of SBP from 136,597 in the original BBJ GWAS to 231,836, indicating that LOG-TRAM can borrow information from UKBB to perform association analysis in EAS. It is also worth noting that LOG-TRAM is computationally efficient. It only took 8 minutes on average to complete the analysis of the whole genome. The timing was evaluated by the Linux computing server with 20 CPU cores of Intel(R) Xeon(R) Gold 6230N CPU @ 2.30GHz processor, 1TB of memory, and a 22 TB solid-state disk.

**Figure 3:**
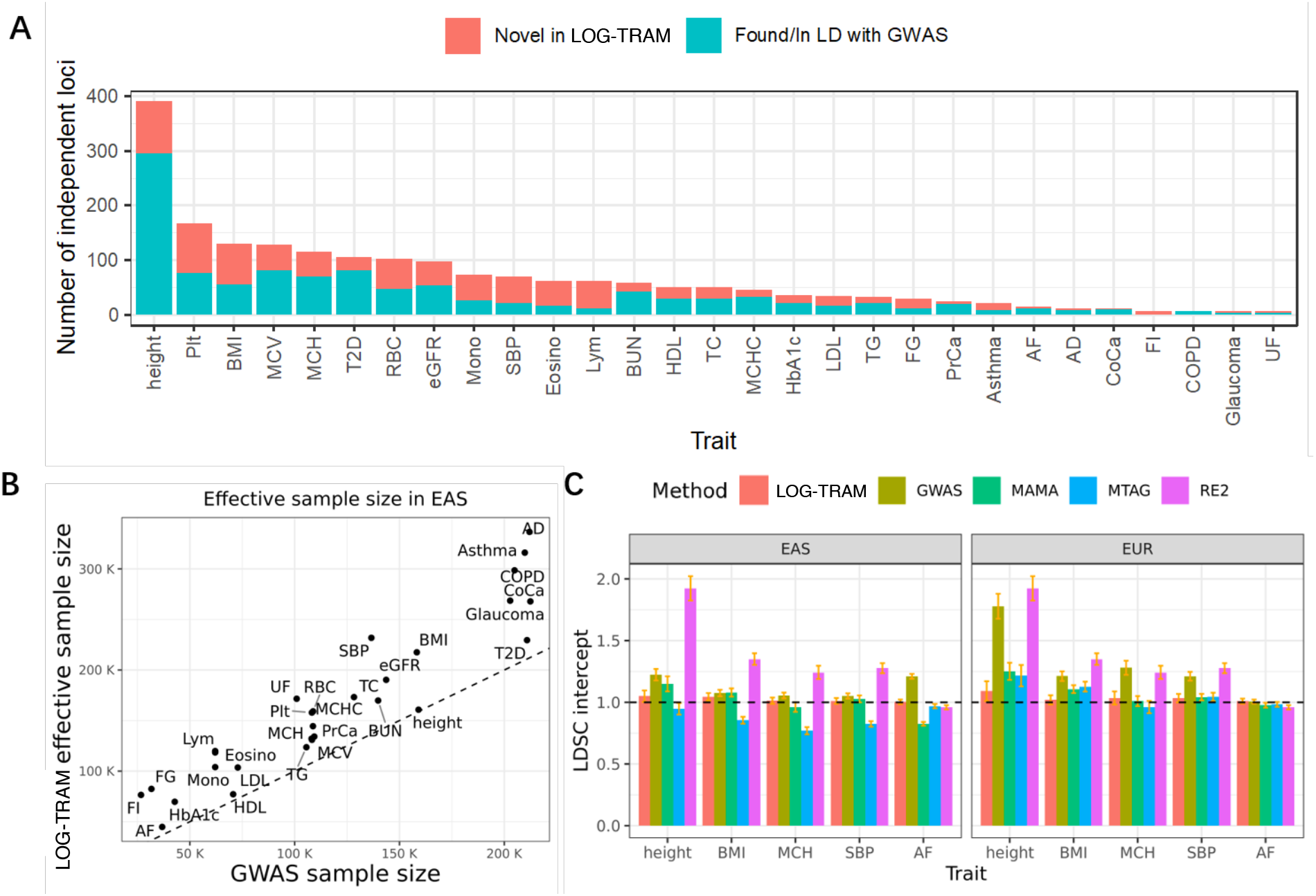
Summary of analysis results obtained by applying LOG-TRAM to combine the EAS and EUR GWAS summary statistics of 29 phenotypes (Table S1). **A**: LOG-TRAM identified independent novel lead SNPs for the 29 phenotypes in EAS compared to the original GWASs. **B**: The effective sample size of association statistics output by LOG-TRAM was computed as 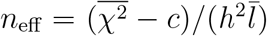, where 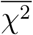 is the mean *χ*^2^ statistics of LOG-TRAM output, *c* is the LDSC intercept of LOG-TRAM summary statistics, *h*^2^ is the heritability of the target trait, and 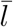 is the mean LD score. **C**: The estimated LDSC intercepts obtained by LOG-TRAM and alternative approaches, including the original GWAS, MAMA, MTAG, and RE2.

Next, we quantified the magnitude of confounding bias in the resulting meta-analysis association statistics using the LDSC intercept [9]. Under the widely accepted LDSC assumption, the LDSC intercept should be one in the absence of confounding bias. As shown in Fig. 3C, the LDSC intercept of the standard EAS height GWAS was estimated to be 1.22 (s.e. = 0.02), suggesting that the standard GWAS suffered from confounding bias. Through the special design of correction for confounding bias, the LDSC intercept of the LOG-TRAM result was reduced to 1.05 (s.e. = 0.02). By contrast, the LDSC intercept of the MTAG results was smaller than one, suggesting that MTAG over-corrected the confounding bias. The LDSC intercept of the MAMA results was 1.15 (s.e. = 0.03), indicating that MAMA was unable to fully correct the confounding biases. RE2 performed the worst with the LDSC intercept increased to 1.92 (s.e. = 0.05). Regarding the association result of EUR (right panel of Fig. 3C), it has been well-known that EUR height GWAS from UKBB heavily suffers from uncorrected population structure with the LDSC intercept as high as 1.78 (s.e. = 0.05). After correction by LOG-TRAM, the LDSC intercept was decreased to 1.09 (s.e. = 0.04). By comparison, the LDSC intercepts of MTAG, MAMA and RE2 were estimated to be 1.22 (s.e. = 0.04), 1.25 (s.e. = 0.03) and 1.92 (s.e. = 0.05), respectively, indicating that the results produced by these methods still suffered from confounding biases. Similarly for other traits, such as body mass index (BMI), Mean corpuscular hemoglobin (MCH), SBP and Atrial fibrillation (AF), the LDSC intercepts of the LOG-TRAM results were all close to one. The above evidence clearly indicates that LOG-TRAM can effectively account for the confounding bias (see Fig. S9-S11 and Table S3-S4 for more examples).

As a concrete example of TRAM, we applied LOG-TRAM to the GWASs of BMI, where the GWAS summary statistics of BBJ (male) and UKBB were used as the inputs. Then we obtained the LOG-TRAM results in EAS and EUR. We used the GWAS of BBJ females as a replication dataset to validate the LOG-TRAM output in EAS. Compared with the original GWAS of BBJ male (Fig. 4A), LOG-TRAM identified 53 novel lead SNPs (Fig. 4B). Among them, we observed the most significant novel lead SNP (Fig. 4C), rs7217403. The local region harboring this variant has 1.1/0.17 = 6.47 and 3.33/0.26 = 12.8 fold enrichment of local heritability in EAS and EUR, respectively (Fig. 4E). Furthermore, we also found that the local co-heritability of this region was higher than the global co-heritability (1.82 compared to 0.14, Fig. 4F), suggesting that LOG-TRAM can effectively leverage the local genetic architecture to boost the power of association mapping. Although rs7217403 is located in an intergenic region, it maps near *MAP2K3*, which has been reported to be significantly associated with BMI across diverse populations, including American Indians [36], Europeans [37, 38], and East Asians [39]. The expression level of *MAP2K3* was positively correlated with BMI in adipose tissue, and in vitro studies suggested that *MAP2K3* was activated during adipogenesis [36]. Given these statistical and biological evidences, *MAP2K3* appears to be a reproducible obesity locus, but knowledge for the molecular mechanisms underlying the association is still lacking. The intergenic variant, rs7217403, identified by LOG-TRAM in EAS may shed light on the genetic etiology of BMI and uncover biologically meaningful variation. In replication analysis, we showed that effect sizes of lead SNPs identified by LOG-TRAM were highly consistent with the replication dataset (Fig. 4F). In particular, the highlighted SNP, rs7217403, showed a notable consistent effect with LOG-TRAM results. Finally, we further verified the LOG-TRAM result by comparing the *p*-values of four sets of SNPs in the replication dataset. As shown in Fig. 4G, the lead SNPs identified by LOG-TRAM showed more significant *p*-values compared to those SNPs identified from the standard GWAS in the European ancestry.

**Figure 4:**
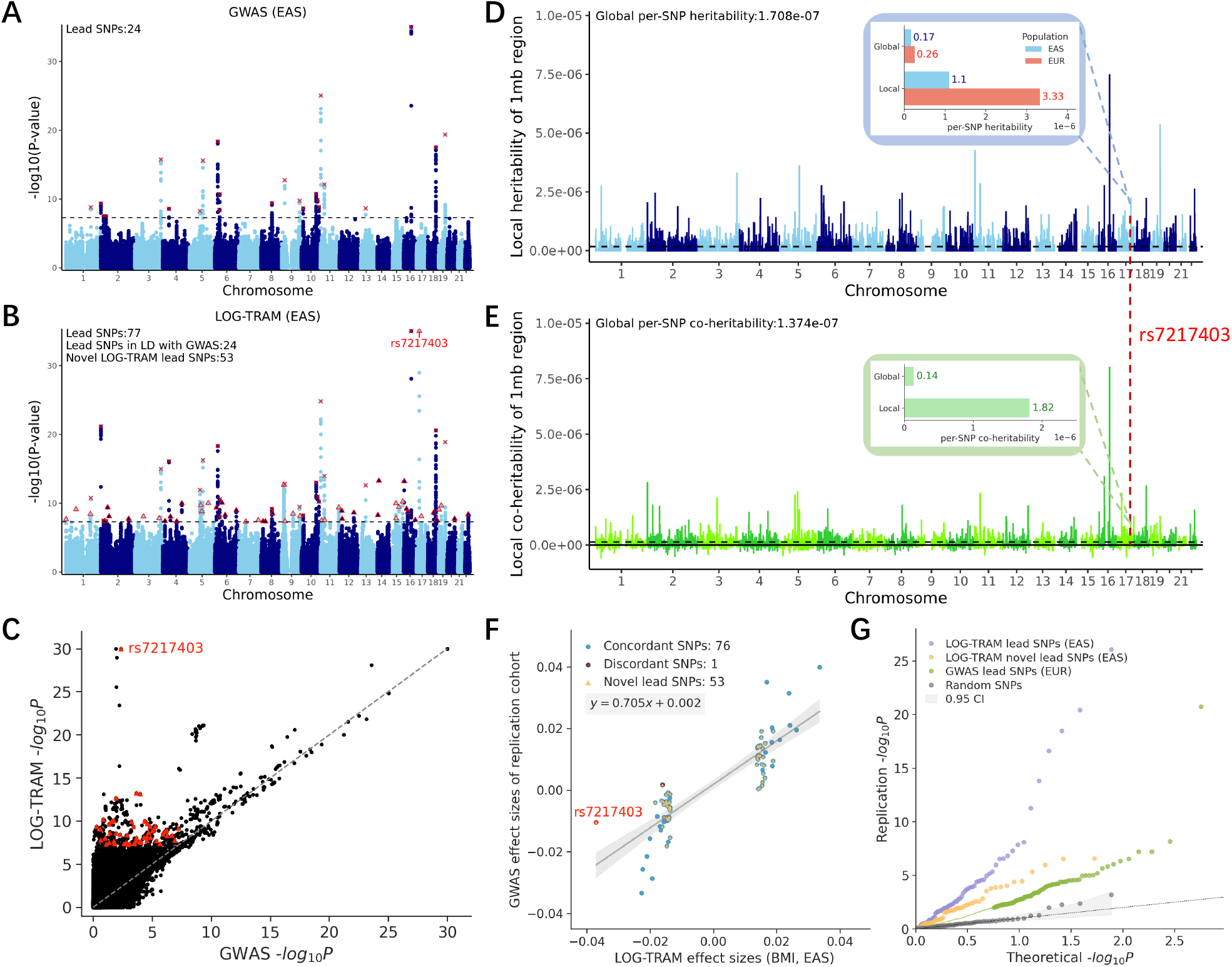
Trans-ancestry association mapping of BMI. **A**: Manhattan plot of the BMI GWAS for the BBJ males, where 24 independent lead SNPs were identified. **B**: Manhattan plot of the LOG-TRAM results for the BBJ male. LOG-TRAM identified 77 independent lead SNPs, of which 53 were not identified in the original BBJ male GWAS. Lead SNPs identified by both LOG-TRAM and GWAS are marked by a cross. Novel lead SNPs are marked by a triangle. **C**: Comparison of the *p*-value output by LOG-TRAM and original BMI GWAS *p*-values of the BBJ male. **D**: Local per-SNP heritability estimated by LOG-TRAM based on BBJ (male) and UKBB. **E**: Local per-SNP co-heritability between BBJ male and UKBB. **F**: Comparison of lead SNPs effect sizes output by LOG-TRAM and effect sizes of GWAS in the replication dataset (the BBJ female). **G**: The QQ plot compares the *p*-values in the replication GWAS data: (1) LOG-TRAM lead SNPs, (2) LOG-TRAM novel lead SNPs, (3) lead SNPs identified from UKBB, and (4) randomly selected SNPs from the replication GWAS. Clearly, SNPs reported by LOG-TRAM are strongly supported in the replication study.

To cross-validate our LOG-TRAM result, we used UKBB and the BBJ females as the discovery cohort and their summary statistics as the input of LOG-TRAM. Then we replicated the LOG-TRAM output using the BBJ males. We observed that the LOG-TRAM results were highly replicable (Fig. S12-S13). Compared with the original BMI GWAS of BBJ females, LOG-TRAM identified 63 novel lead SNPs (Fig. S12B). Similarly, we observed the same most significant novel lead SNP (Fig. S12C), rs7217403. The local region harboring this variant showed a consistent local genetic architecture with 1.81/0.19 = 9.53 fold enrichment for local heritability in EAS, and 2.33/0.16 = 14.5 fold enrichment for local co-heritability (Fig. S12D).

#### Identification of ancestry-specific loci

Taking the GWAS summary statistics from BBJ and UKBB as input, we applied the LOG-TRAM-based difference test to a number of complex traits, including mean corpuscular hemoglobin concentration (MCHC), mean corpuscular volume (MCV), high-density-lipoprotein (HDL), low-density-lipoprotein (LDL), type 2 diabetes (T2D), and BMI. The test statistics 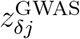 and 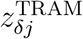 for overlapping SNPs across two populations were computed using Eq. (20) and Eq. (21). Here we take MCHC as an example. In addition to the 23 ancestry-specific lead SNPs (*p*-value*<* 5 *×* 10^−8^) identified by the standard GWAS-based difference test, the LOG-TRAM-based difference test identified six novel lead SNPs with significantly different effect sizes between EAS and EUR (Fig. 5A and B). Although those novel SNPs (red dot in Fig. 5C) show different effect sizes between EAS and EUR, they were not identified by the GWAS-based difference test due to the limited effective sample size. In contrast, the LOG-TRAM-based difference test successfully captured those signals as LOG-TRAM can increase the effective sample size. From a biological perspective, these SNPs identified by the LOG-TRAM-based difference test were enriched in functional categories related to population differences, suggesting the functional importance of ancestry-specific SNPs. Notably, as shown in Fig. 5D, SNPs in the top quantile of background selection statistic [40] have more significant *p*-values, while SNPs in the top quantile of recombination rate [41] have less significant *p*-values than the average. This phenomenon is consistent with the observation in [29], where the background selection statistic is positively correlated to the depletion of trans-ancestry genetic correlation while recombination rate has a reverse pattern. The ancestry-specific loci identification results of other five traits are provided in Fig. S15-S19.

**Figure 5:**
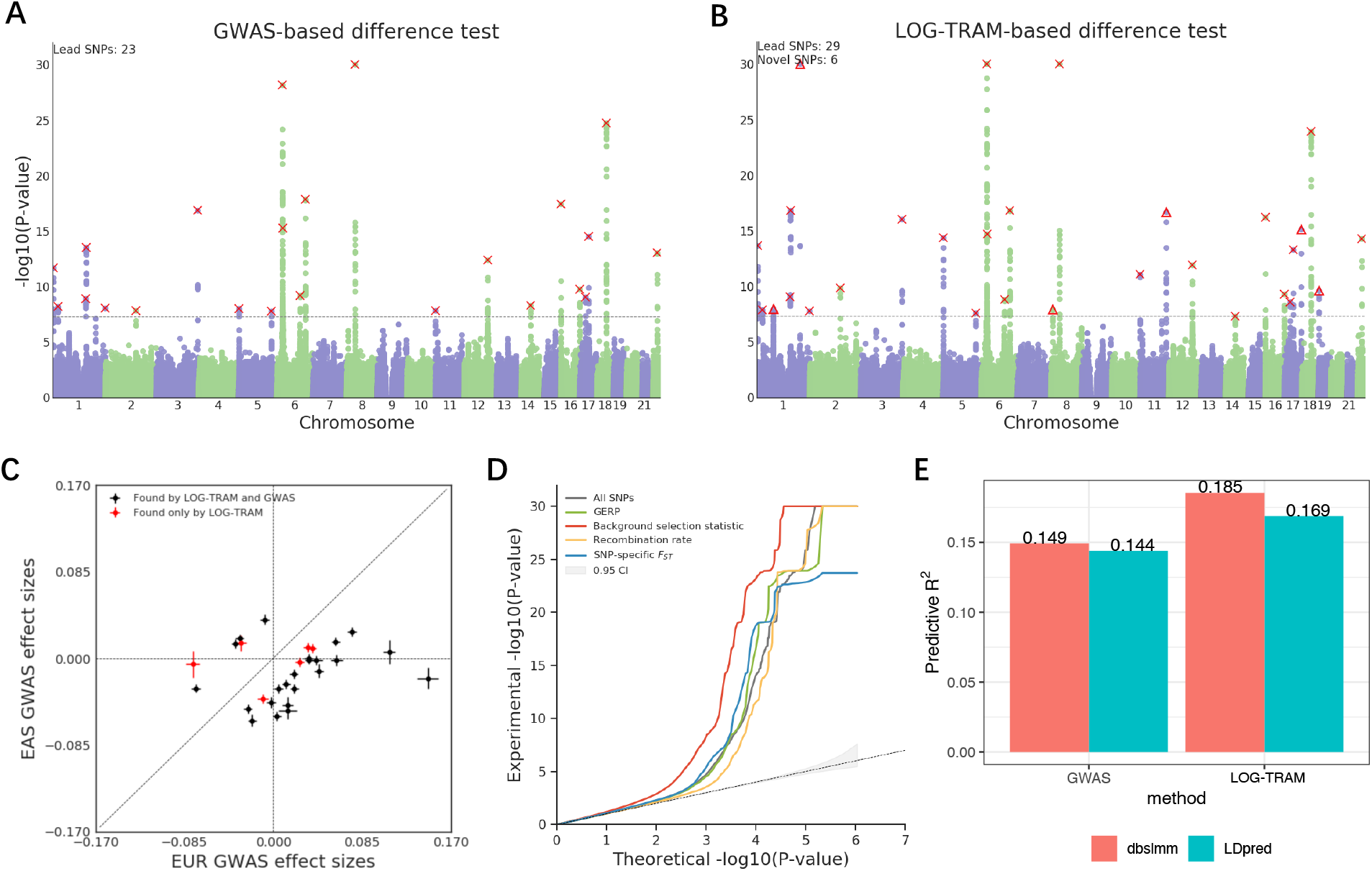
Applications of LOG-TRAM. **A**: Manhattan plot of GWAS-based difference test results for MCHC. Lead SNPs are marked by a cross. **B**: Manhattan plot of LOG-TRAM-based difference test results for MCHC. The dashed line marks the threshold for genome-wide significance (*P* = 5 *×* 10^−8^). Lead SNP identified by both the LOG-TRAM and GWAS-based difference tests are marked by crosses. Novel lead SNPs are marked by triangles. **C**: Comparison of MCHC GWAS effect sizes for lead SNPs between EAS and EUR. Red points denote novel loci identified by the LOG-TRAM-based difference test; black points denote lead loci identified in both the LOG-TRAM and GWAS-based difference tests. Vertical and horizontal error bars represent standard errors of GWAS effect sizes in EAS and EUR, respectively. **D** The QQ-plots of *p*-values for all SNPs and SNPs in the top quantile of four continuous-valued annotations. **E** Predictive *R*^2^ for EAS height PRS models based on GWAS and LOG-TRAM association statistics, respectively.

#### The LOG-TRAM result for construction of polygenic risk scores

To demonstrate the utility of LOG-TRAM in the construction of polygenic risk scores (PRSs) in under-represented populations, we compared the predictive power of PRS constructed by using the standard GWAS summary statistics and the LOG-TRAM summary statistics. We considered construction of PRS for human height as an example. We first applied the most commonly used PRS method, LDpred [42], to build an EAS height PRS model (denoted as GWAS-based PRS) using the GWAS summary statistics of height from BBJ (*n*=159,095). Using the GWAS summary statistics of height from BBJ and UKBB, we applied LOG-TRAM to generate association statistics for EAS population. Then we used the summary statistics (output of LOG-TRAM) to construct PRS, which is referred to as LOG-TRAM-based PRS. After that, we evaluated the prediction performance in an independent Chinese testing data (*n* = 35,908) recruited from the WeGene platform [22]. In detail, we measured the prediction accuracy of each PRS model using the predictive *R*^2^, defined as the square of correlation of predicted PRSs and true residual phenotypes after regressing out covariates (e.g., gender, age, genomics PCs) [22, 43].

As shown in Fig. 5E, the predictive accuracy obtained by GWAS-based PRS is *R*^2^ = 0.144. In contrast, PRS constructed from the LOG-TRAM association statistics achieved (0.169-0.144)/0.144) ≈ 17% accuracy gain, indicating that the LOG-TRAM association statistics of EAS height successfully borrowed information from the large-scale UKBB datasets. The improvement of prediction accuracy was quite stable when different PRS approaches were applied. When we applied a recently developed PRS method named ‘dbslmm’ [44], we obtained predictive *R*^2^ = 0.149 using the GWAS summary statistics. With the same PRS method, we achieved predictive *R*^2^ = 0.185 using the LOG-TRAM association statistics, which was (0.185-0.149)/0.149 ≈ 24% improvement. In summary, the above results suggest that the LOG-TRAM output can lead to a significant improvement in construction of accurate PRS for under-represented populations.

## Discussion

In this paper, we have introduced a novel trans-ancestry meta-analysis method, LOG-TRAM, aiming to improve the statistical power of GWASs in non-Europeans by leveraging locally shared genetic architectures with biobank-scale auxiliary datasets. Through comprehensive simulations, we showed that our method has greater statistical power while controlling the type I error rate compared to existing approaches, and our method is robust across various genetic architecture settings. We applied LOG-TRAM to GWASs of 29 complex traits and diseases from EAS and EUR, achieving substantial gains in power and effective correction of confounding biases. We found that LOG-TRAM results were reproducible in independent studies. Finally, we demonstrated that the LOG-TRAM results can be further used for identification of ancestry-specific loci and construction of PRS in under-represented populations.

As we mentioned, accumulating evidence reveal that the shared genetic basis between phenotypes/populations vary across genomic regions [25, 26, 28, 29]. For example, the effect sizes of SNPs can be phenotype/population-specific, and the genetic architecture of multiple ancestries can be locally different. Therefore, assuming a constant variance–covariance matrix of effect sizes across the whole genome violates biological intuition in many circumstances. Stratified LDSC [45] is a representative method to explore the local genetic architecture. It assumes that SNPs in different genomic regions (e.g., functional categories) contribute disproportionately to the heritability and estimates the per-SNP heritability in each region by regressing the association statistics to the LD score corresponding to each region. Recently, several other statistical methods, including *ρ*-HESS [26], SUPERGNOVA [27], and LOGODetect [30], have been developed to estimate local genetic correlation across traits in a single population. Their analysis results consistently suggest that the local genetic correlation can greatly differ from the global genetic correlation, offering new insights into complex human diseases and traits. However, the methods that explore the local genetic architecture have not accounted for heterogeneity across ancestries. A direct application of these methods for association mapping in the trans-ancestry setting is still problematic. To develop the LOG-TRAM method, we considered a sliding window (e.g., 1 M base-pair segment) to decompose the whole genome into local region ***ℛ*** and background region ***ℬ***, which allows us to model the regional genetic architectures differently from the global pattern. By successfully leveraging the local genetic architecture and accounting for confounding bias, not only can LOG-TRAM estimate local heritability but it also can be applied for association mapping in the trans-ancestry setting.

Although we have mainly focused on the local regions partitioned by the sliding window approach with a fixed window size in this study, it is worth mentioning that LOG-TRAM can also be applied to more generalized “local regions” defined by functional annotations or tissue/cell-type specifically expressed gene (SEG) annotations. Indeed, SNPs with biological functions (e.g., gene regulatory elements [46, 47, 48, 49], epigenomic regulations [45, 50, 51], and tissue-specific expressed genes [52, 29]) have been well known for their enrichment in the heritability of complex traits, which re-emphasizes the widespread of heterogeneous genetic architectures across the genome. Consequently, these functional annotations are widely used in genetic studies to increase statistical power, including GWAS [53, 54, 55], pleiotropy [56], fine-mapping [57, 58], and polygenic risk scores [59, 60]. Very recently, a study [61] suggests that functional annotations have great potential in improving the portability of trans-ancestry polygenic risk scores, indicating the substantial share of biological mechanisms across populations. Therefore, leveraging the functional/SEG annotations can also increase the power of trans-ancestry association mapping for under-represented populations. To see this, we considered SNPs annotated by 20 binary functional annotations from Baseline-LD-X model [29] as local regions and estimated their local genetic architectures for LOG-TRAM. Intuitively, each annotation can be analogized to one 1 M base-pair segment defined in the previous section. We applied the functional-informed LOG-TRAM to the GWASs of BMI and T2D from BBJ and UKBB. Compared to the original EAS GWASs, we identified 85% and 16% more independently significant loci for BMI and T2D (Fig. S22 and S25), respectively. In addition, we further used the 53 SEG annotations [29] as local regions and applied the SEG-informed LOG-TRAM to the same traits. As shown in Fig. S22 and S25, SEG-informed LOG-TRAM achieved comparable power gains in EAS and identified 114% and 25% more independently significant loci for BMI and T2D, respectively.

Our LOG-TRAM approach needs more investigation in the following directions. First, it has been shown that pleiotropic effects are widespread in the genome [62]. In a systematic analysis of 4,155 publicly available GWASs, 90% trait-associated loci affect multiple phenotypes simultaneously [2]. Hence, jointly modeling of multiple GWAS traits across populations may further boost the statistical power of trans-ancestry association mapping. Second, LOG-TRAM assumes that, for a given population, SNPs with lower allele frequencies tend to have larger effect sizes. More specifically, we considered the relationship of per-allele effect size *v* and allele frequency *f* as Var(*v*) ∝ [*f* (1 − *f*)]^*α*^, where *α* = −1. This assumption has been shown to be a stable choice in simulation studies [63], and it was also adopted in the previous trans-ancestry analysis [64]. Some recent studies on the estimation of *α* found that although estimated *α* are negative for most complex traits, they vary moderately across traits [65, 66]. Therefore, it would be more appropriate to obtain an estimate of *α* for each trait rather than fixing *α* = −1. We will explore these potential improvements in the near future.

## Appendix A: Derivation of the covariance of marginal effect size

We first apply the law of total expectation to obtain the first moments of 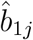 and 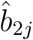:

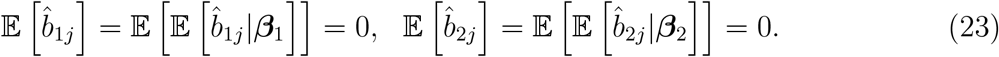

By the law of total variance, we have

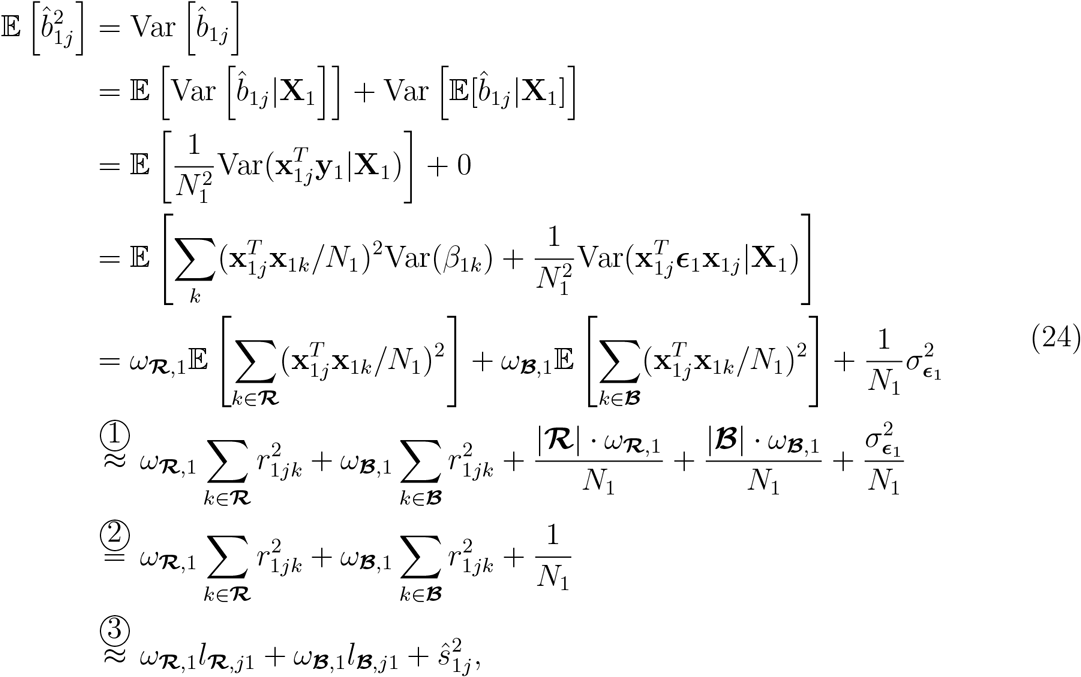

where we use 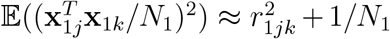 in approximation ➀, 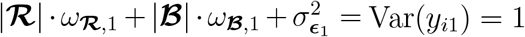 for equation ➁, and approximation ➂ is derived from

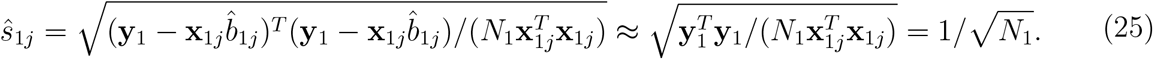

Similarly, we can derive 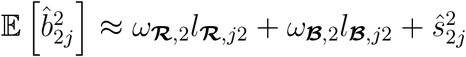 and the covariance between 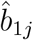 and 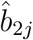 can be obtained as

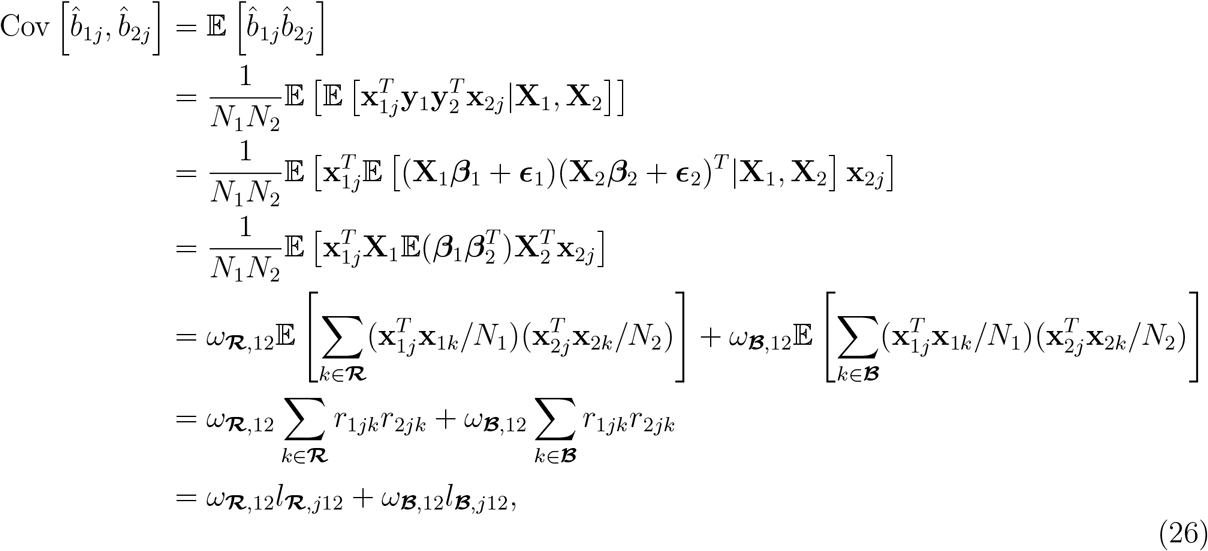

## Appendix B: The LOG-TRAM model accounting for population stratification

Now we consider the LOG-TRAM in the presence of population stratification. Following LDSC [9], we model population stratification by assuming that the samples in each population are a 50:50 mixture of two sub-groups with following genetic drift model: (i) for individual *i* in sub-group 1, we have 𝔼(**x**_1*j,i*_|*i* ∈ sub-group 1) = *f*_1*j*_ and 𝔼(**x**_2*j,i*_|*i* ∈ sub-group 1) = *f*_2*j*_. For individual *i* in sub-group 2, we have 𝔼(**x**_1*j,i*_|*i* ∈ sub-group 2) = −*f*_1*j*_ and 𝔼(**x**_2*j,i*_|*i* ∈ sub-group 2) = −*f*_2*j*_. We also have Var(**x**_1*j,i*_) = Var(**x**_2*j,i*_) = 1 for all SNP *j* because genotype matrices are standardized; (ii) the genetic drift term *f*_1*j*_ ∼ *𝒩*(0, *F*_1,*ST*_) and *f*_2*j*_ ∼ *𝒩*(0, *F*_2,*ST*_), where *F*_1,*ST*_ and *F*_2,*ST*_ are Wright’s *F*_*ST*_ to measure the allele frequency differences between the sub-groups in population 1 and 2, respectively (genetic stratification); and (iii) we assume the genetic drift term between the two populations has a covariance defined as Cov(*f*_1*j*_, *f*_2*j*_) = *F*_cov_, and Cov(*f*_1*j*_, *f*_1*k*_) = Cov(*f*_2*j*_, *f*_2*k*_) = 0 for all *j* ≠ *k*. Besides, we use *σ*_1,*p*_ and *σ*_2,*p*_ as the mean phenotype differences between the subgroups in population 1 and 2, respectively (environmental stratification). Without loss of generality, we assume that the level of genetic and environmental stratification are similar between the two populations. Therefore, we have *F*_1,*ST*_ ≈ *F*_2,*ST*_ := *F*_*ST*_ and *σ*_1,*p*_ ≈ *σ*_2,*p*_ := *σ*_*p*_. Then we can update the Eq. (3) as:

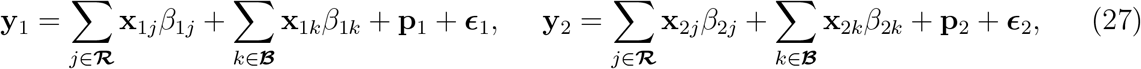

where 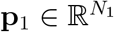 and 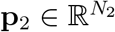 are the independent environmental stratification terms defined as

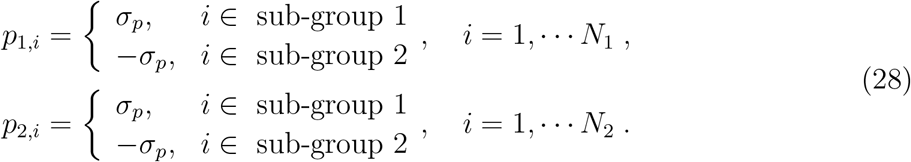

Here we assume that environmental stratification term (**p**) and noise term (***ϵ***) are independent of each other and we assume the the variances of error terms satisfy 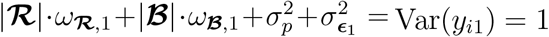 and 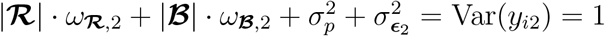.

Accordingly, we have

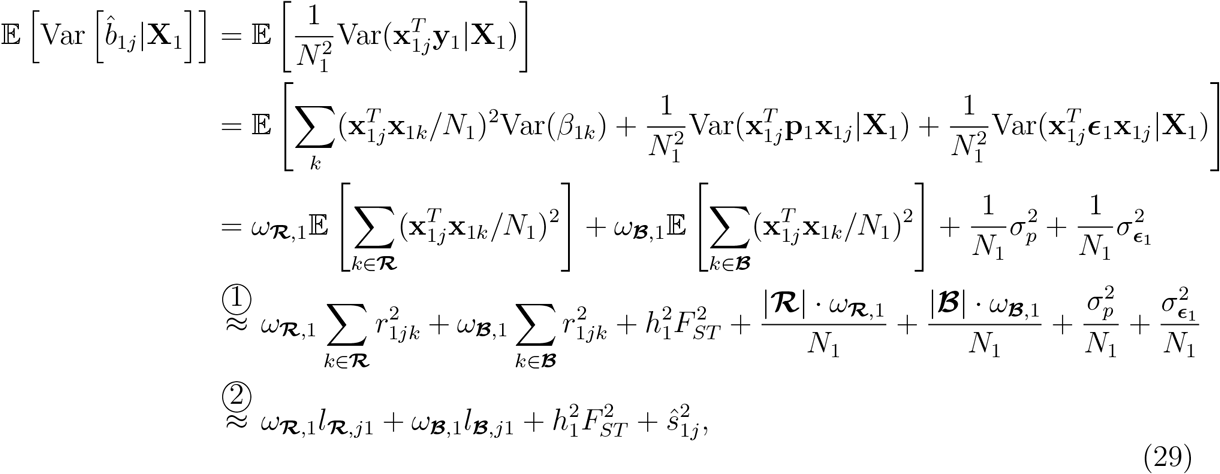

where we use 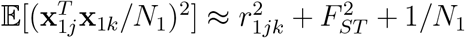 in approximation ➀, and 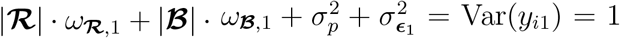 in approximation ➁, and 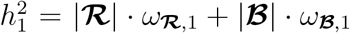 is the heritability of **y**_1_. Next, we can derive

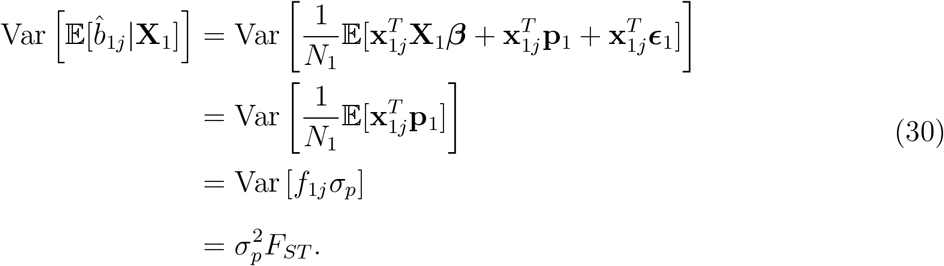

Thus, by the law of total variance, we have

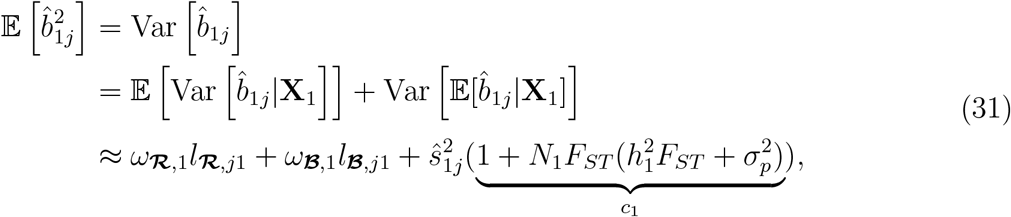

Similarly, we have

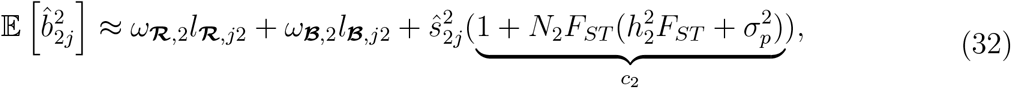

where 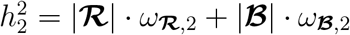 is the heritability of **y**_2_. Eqs. (31) and (32) show that the intercepts *c*_1_ and *c*_2_ will deviate from one in the presence of population stratification (*F*_*ST*_ ≠ 0), leading to inflated test statistics.

Now we derive the covariance term 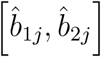, which is given by

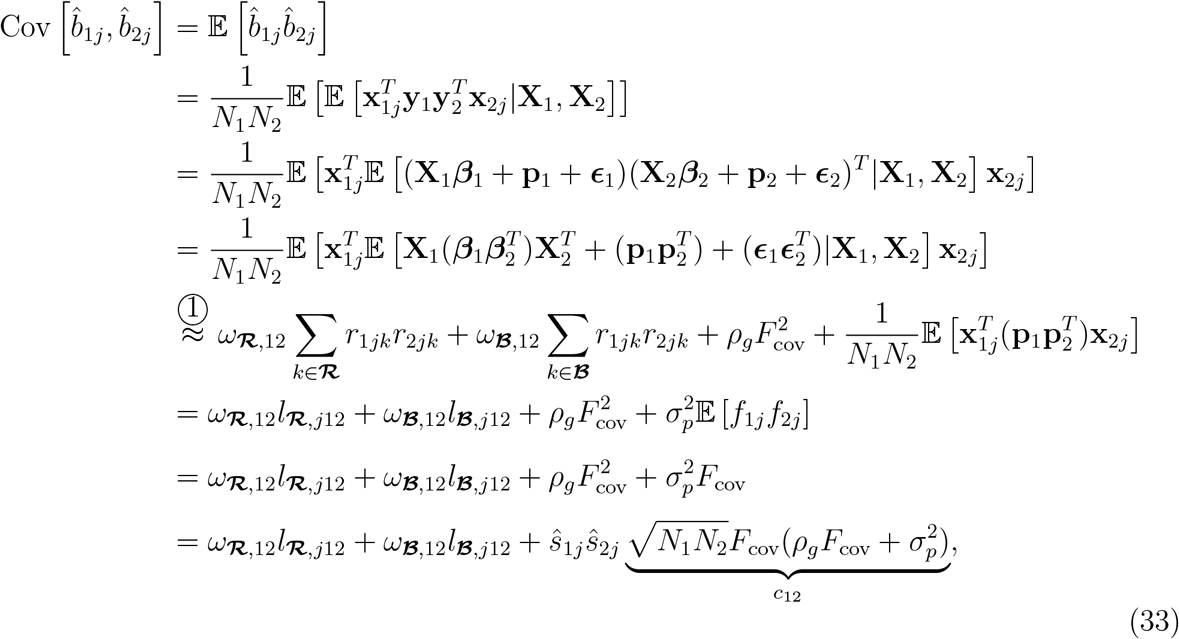

where we use 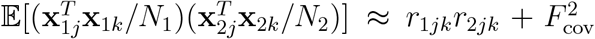 in approximation ➀ [67], and *ρ*_*g*_ = *ω*_***ℛ***,12_|***ℛ***| + *ω*_***ℬ***,12_|***ℬ***| is the genetic covariance between **y**_1_ and **y**_2_. Eq.(33) shows that the intercept *c*_12_ will deviate from zero in the presence of non-zero genetic drift correlation between the two populations (Cov[*f*_1*j*_, *f*_2*j*_] = *F*_cov_ ≠ 0).

In summary, LOG-TRAM model accounts for the population stratification using the following relationships

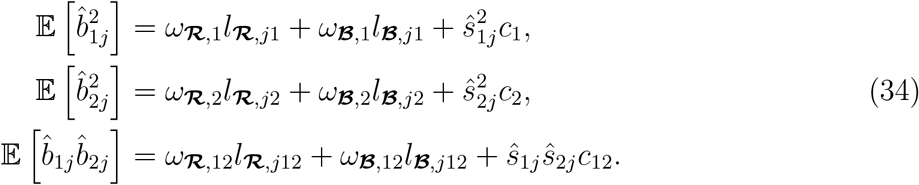

## Appendix C: Derivation of the moment conditions

Following the previous studies [20, 21], LOG-TRAM is also a GMM estimator [68] defined by a set of moment conditions and a weight matrix. The LOG-TRAM moment condition can be derived by the linear projection of GWAS estimates for SNP *j* (i.e., 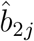) onto the true marginal effect of SNP *j* (i.e., *b*_1*j*_) as

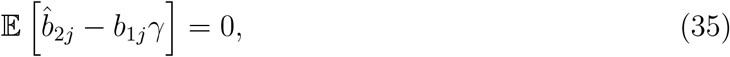

where *γ* is unknown coefficient. The GMM estimator for *γ* is derived from minimizing the following objective function using an identity matrix as the weight matrix

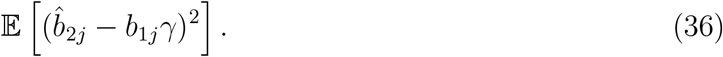

By taking the derivative with respect to *γ* and setting it equal to zero

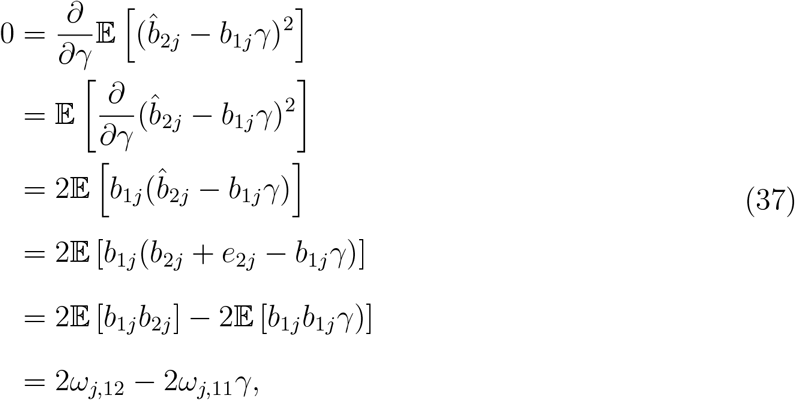

we obtain

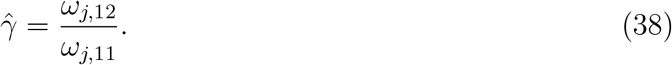

Finally, we show that the projection of 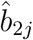 onto the true marginal effect *b*_1*j*_ is 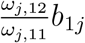. Similarly, we can extend moment condition (35) to

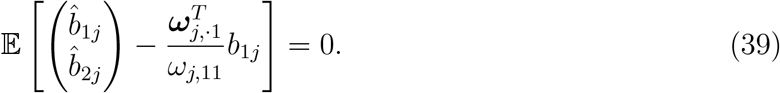

Therefore, we have conditional mean for 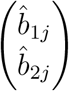 as

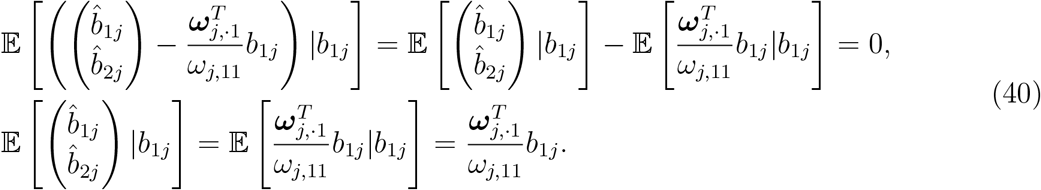

Next, we show that the conditional variance can be derive as

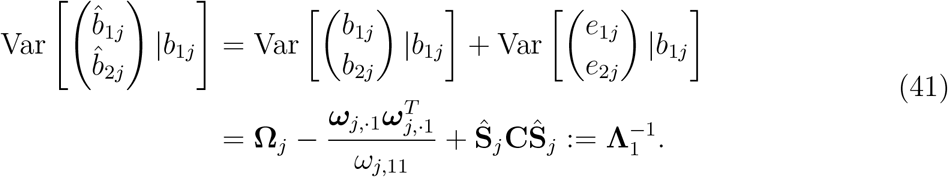

Similarly, we can obtain GMM estimator of *b*_1*j*_ by minimizing the following objective function with **Λ**_1_ as the weight matrix

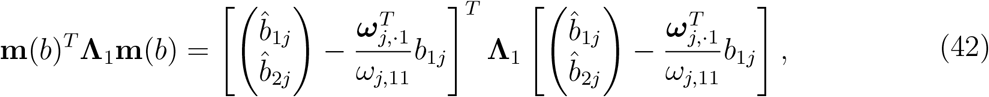

where 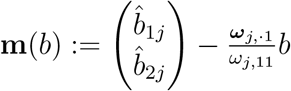. By taking the derivative with respect to *b*_1*j*_ and setting it equal to zero

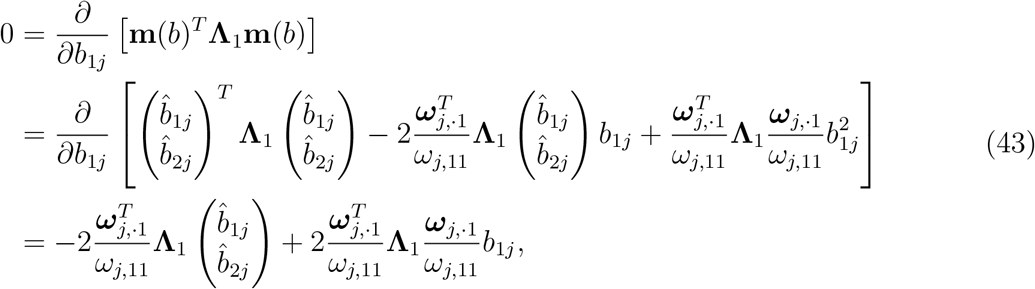

we solve the equation and obtain

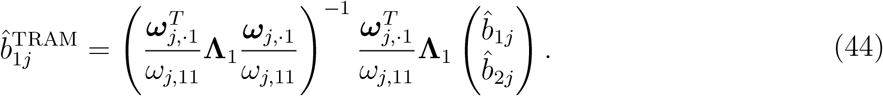

Using the established theoretical property of the GMM estimators, the LOG-TRAM estimator has asymptotic distribution

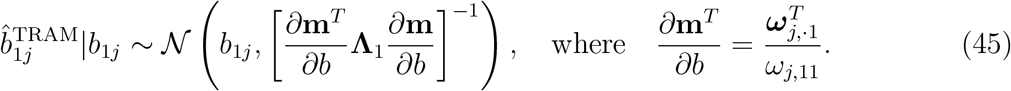

## Appendix D: Theoretical properties of LOG-TRAM

Following previous studies [20, 21], we can show that LOG-TRAM is also the best linear unbiased estimator (BLUE). To see this, we first show the LOG-TRAM estimate (e.g., 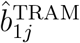) is an unbiased estimator of the true marginal effect *b*_1*j*_:

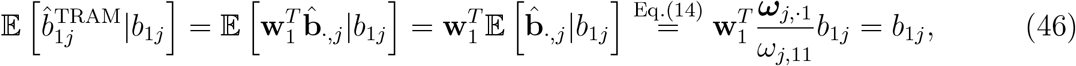

where **w**_1_ and 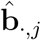 are defined in Eq. (16). Next, we consider a necessary condition for the set of all unbiased linear estimators denoted as 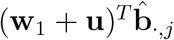, where **u** represents some vectors such that 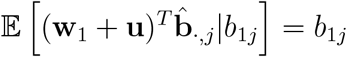. It follows that

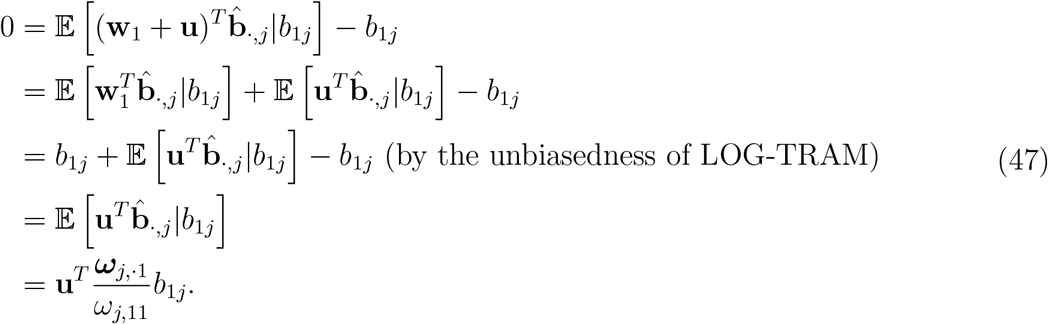

Therefore, we obtain **u**^*T*^ ***ω***_*j*, ·1_ = 0. We then show that the variance 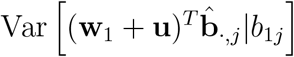 is larger than the variance 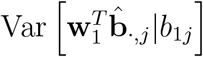. We expand

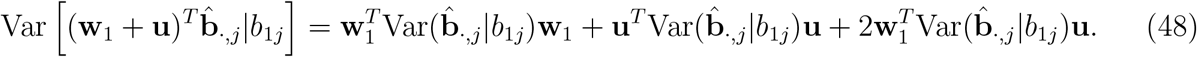

Based on Eq. (14) and Eq. (16), we have

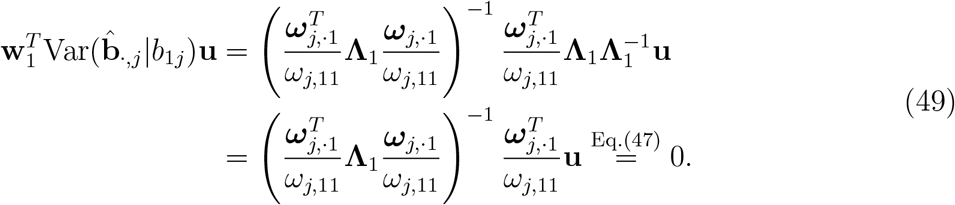

Therefore, we have 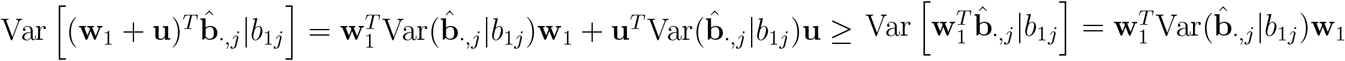. Therefore, LOG-TRAM has the lowest variance among all the unbiased estimators.

## Web resources

LDSC: https://github.com/bulik/ldsc;

LOG-TRAM: https://github.com/YangLabHKUST/LOG-TRAM;

MTAG: https://github.com/JonJala/mtag;

MAMA: https://github.com/JonJala/mama;

PLINK: https://www.cog-genomics.org/plink;

BOLT-LMM: https://alkesgroup.broadinstitute.org/BOLT-LMM.

LDpred: https://github.com/bvilhjal/ldpred;

dbslmm: https://biostat0903.github.io/DBSLMM;

S-LDXR: https://github.com/huwenboshi/s-ldxr;

UKBB: https://www.ukbiobank.ac.uk;

BBJ: http://jenger.riken.jp/en

## Data and Code Availability

The publicly available GWAS summary statistics for meta-analysis were obtained from the links summarized in Table S1. Access to the Chinese data has to be approved by the contact authors. Researchers who wish to gain access to the data are required to submit the data request by sending an email to the contact authors. The UKBB data are available through the UK Biobank Access Management System. LOG-TRAM software and source codes in this study were publicly available in Web resources.

## Acknowledgements

This work is supported in part by National Key R&D Program of China (2020YFA0713900), Hong Kong Research Grant Council [12301417, 16307818, 16301419, 16308120], Hong Kong Innovation and Technology Fund [PRP/029/19FX], Hong Kong University of Science and Technology [startup grant R9405, Z0428 from the Big Data Institute] and the Open Research Fund from Shenzhen Research Institute of Big Data [2019ORF01004]. The computational task for this work was partially performed using the X-GPU cluster supported by the RGC Collaborative Research Fund: C6021-19EF.

## References

[1] Robert J Klein, Caroline Zeiss, Emily Y Chew, Jen-Yue Tsai, Richard S Sackler, Chad Haynes, Alice K Henning, John Paul SanGiovanni, Shrikant M Mane, Susan T Mayne, et al. Complement factor H polymorphism in age-related macular degeneration. Science, 308(5720):385–389, 2005.

[2] Kyoko Watanabe, Sven Stringer, Oleksandr Frei, Maša Umićević Mirkov, Christiaan de Leeuw, Tinca JC Polderman, Sophie van der Sluis, Ole A Andreassen, Benjamin M Neale, and Danielle Posthuma. A global overview of pleiotropy and genetic architecture in complex traits. Nature genetics, 51(9):1339–1348, 2019.

[3] Vivian Tam, Nikunj Patel, Michelle Turcotte, Yohan Bossé, Guillaume Paré, and David Meyre. Benefits and limitations of genome-wide association studies. Nature Reviews Genetics, 20(8):467–484, 2019.

[4] Melinda C Mills and Charles Rahal. The gwas diversity monitor tracks diversity by disease in real time. Nature genetics, 52(3):242–243, 2020.

[5] Deepti Gurdasani, Inês Barroso, Eleftheria Zeggini, and Manjinder S Sandhu. Genomics of disease risk in globally diverse populations. Nature Reviews Genetics, 20(9):520–535, 2019.

[6] Lucia A Hindorff, Vence L Bonham, Lawrence C Brody, Margaret EC Ginoza, Carolyn M Hutter, Teri A Manolio, and Eric D Green. Prioritizing diversity in human genomics research. Nature Reviews Genetics, 19(3):175–185, 2018.

[7] 1000 Genomes Project Consortium et al. A global reference for human genetic variation. Nature, 526(7571):68–74, 2015.

[8] Genevieve L Wojcik, Mariaelisa Graff, Katherine K Nishimura, Ran Tao, Jeffrey Haessler, Christopher R Gignoux, Heather M Highland, Yesha M Patel, Elena P Sorokin, Christy L Avery, et al. Genetic analyses of diverse populations improves discovery for complex traits. Nature, 570(7762):514–518, 2019.

[9] Brendan K Bulik-Sullivan, Po-Ru Loh, Hilary K Finucane, Stephan Ripke, Jian Yang, Nick Patterson, Mark J Daly, Alkes L Price, and Benjamin M Neale. Ld score regression distinguishes confounding from polygenicity in genome-wide association studies. Nature genetics, 47(3):291–295, 2015.

[10] Alkes L Price, Nick J Patterson, Robert M Plenge, Michael E Weinblatt, Nancy A Shadick, and David Reich. Principal components analysis corrects for stratification in genome-wide association studies. Nature genetics, 38(8):904–909, 2006.

[11] Jian Yang, Noah A Zaitlen, Michael E Goddard, Peter M Visscher, and Alkes L Price. Advantages and pitfalls in the application of mixed-model association methods. Nature genetics, 46(2):100–106, 2014.

[12] Abdel Abdellaoui, David Hugh-Jones, Loic Yengo, Kathryn E Kemper, Michel G Nivard, Laura Veul, Yan Holtz, Brendan P Zietsch, Timothy M Frayling, Naomi R Wray, et al. Genetic correlates of social stratification in great britain. Nature human behaviour, 3(12):1332–1342, 2019.

[13] Simon Haworth, Ruth Mitchell, Laura Corbin, Kaitlin H Wade, Tom Dudding, Ashley Budu-Aggrey, David Carslake, Gibran Hemani, Lavinia Paternoster, George Davey Smith, et al. Apparent latent structure within the UK Biobank sample has implications for epidemiological analysis. Nature communications, 10(1):1–9, 2019.

[14] Rebecca DerSimonian and Nan Laird. Meta-analysis in clinical trials. Controlled clinical trials, 7(3):177–188, 1986.

[15] Evangelos Evangelou and John PA Ioannidis. Meta-analysis methods for genome-wide association studies and beyond. Nature Reviews Genetics, 14(6):379–389, 2013.

[16] Yun R Li and Brendan J Keating. Trans-ethnic genome-wide association studies: advantages and challenges of mapping in diverse populations. Genome medicine, 6(10):1–14, 2014.

[17] Buhm Han and Eleazar Eskin. Random-effects model aimed at discovering associations in meta-analysis of genome-wide association studies. The American Journal of Human Genetics, 88(5):586–598, 2011.

[18] CH Lee, Eleazar Eskin, and Buhm Han. Increasing the power of meta-analysis of genomewide association studies to detect heterogeneous effects. Bioinformatics, 33(14):i379–i388, 2017.

[19] Andrew P Morris. Transethnic meta-analysis of genomewide association studies. Genetic epidemiology, 35(8):809–822, 2011.

[20] Patrick Turley, Raymond K Walters, Omeed Maghzian, Aysu Okbay, James J Lee, Mark Alan Fontana, Tuan Anh Nguyen-Viet, Robbee Wedow, Meghan Zacher, Nicholas A Furlotte, et al. Multi-trait analysis of genome-wide association summary statistics using mtag. Nature genetics, 50(2):229–237, 2018.

[21] Patrick Turley, Alicia R Martin, Grant Goldman, Hui Li, Masahiro Kanai, Raymond K Walters, Jonathan B Jala, Kuang Lin, Iona Y Millwood, Caitlin E Carey, et al. Multiancestry meta-analysis yields novel genetic discoveries and ancestry-specific associations. bioRxiv, 2021.

[22] Mingxuan Cai, Jiashun Xiao, Shunkang Zhang, Xiang Wan, Hongyu Zhao, Gang Chen, and Can Yang. A unified framework for cross-population trait prediction by leveraging the genetic correlation of polygenic traits. The American Journal of Human Genetics, 108(4):632–655, 2021.

[23] Yang Luo, Xinyi Li, Xin Wang, Steven Gazal, Josep Maria Mercader, Me Research Team, SIGMA Type 2 Diabetes Consortium, Benjamin M Neale, Jose C Florez, Adam Auton, et al. Estimating heritability and its enrichment in tissue-specific gene sets in admixed populations. Human molecular genetics, 30(16):1521–1534, 2021.

[24] Matthew Stephens and David J Balding. Bayesian statistical methods for genetic association studies. Nature Reviews Genetics, 10(10):681–690, 2009.

[25] Po-Ru Loh, Gaurav Bhatia, Alexander Gusev, Hilary K Finucane, Brendan K Bulik-Sullivan, Samuela J Pollack, Teresa R de Candia, Sang Hong Lee, Naomi R Wray, Kenneth S Kendler, et al. Contrasting genetic architectures of schizophrenia and other complex diseases using fast variance-components analysis. Nature genetics, 47(12):1385–1392, 2015.

[26] Huwenbo Shi, Nicholas Mancuso, Sarah Spendlove, and Bogdan Pasaniuc. Local genetic correlation gives insights into the shared genetic architecture of complex traits. The American Journal of Human Genetics, 101(5):737–751, 2017.

[27] Yiliang Zhang, Qiongshi Lu, Yixuan Ye, Kunling Huang, Wei Liu, Yuchang Wu, Xiaoyuan Zhong, Boyang Li, Zhaolong Yu, Brittany G Travers, et al. SUPERGNOVA: local genetic correlation analysis reveals heterogeneous etiologic sharing of complex traits. Genome biology, 22(1):1–30, 2021.

[28] Josefin Werme, Sophie van der Sluis, Danielle Posthuma, and Christiaan de Leeuw. Lava: An integrated framework for local genetic correlation analysis. bioRxiv, pages 2020–12, 2021.

[29] Huwenbo Shi, Steven Gazal, Masahiro Kanai, Evan M Koch, Armin P Schoech, Katherine M Siewert, Samuel S Kim, Yang Luo, Tiffany Amariuta, Hailiang Huang, et al. Population-specific causal disease effect sizes in functionally important regions impacted by selection. Nature communications, 12(1):1–15, 2021.

[30] Hanmin Guo, James J Li, Qiongshi Lu, and Lin Hou. Detecting local genetic correlations with scan statistics. Nature communications, 12(1):1–13, 2021.

[31] Wouter van Rheenen, Wouter J Peyrot, Andrew J Schork, S Hong Lee, and Naomi R Wray. Genetic correlations of polygenic disease traits: from theory to practice. Nature Reviews Genetics, 20(10):567–581, 2019.

[32] Po-Ru Loh, George Tucker, Brendan K Bulik-Sullivan, Bjarni J Vilhjalmsson, Hilary K Finucane, Rany M Salem, Daniel I Chasman, Paul M Ridker, Benjamin M Neale, Bonnie Berger, et al. Efficient bayesian mixed-model analysis increases association power in large cohorts. Nature genetics, 47(3):284–290, 2015.

[33] Jian Yang, S Hong Lee, Michael E Goddard, and Peter M Visscher. Gcta: a tool for genome-wide complex trait analysis. The American Journal of Human Genetics, 88(1):76–82, 2011.

[34] Brendan Bulik-Sullivan, Hilary K Finucane, Verneri Anttila, Alexander Gusev, Felix R Day, Po-Ru Loh, Laramie Duncan, John RB Perry, Nick Patterson, Elise B Robinson, sset al. An atlas of genetic correlations across human diseases and traits. Nature genetics, 47(11):1236–1241, 2015.

[35] Cristen J Willer, Yun Li, and Gonçalo R Abecasis. Metal: fast and efficient meta-analysis of genomewide association scans. Bioinformatics, 26(17):2190–2191, 2010.

[36] Li Bian, Michael Traurig, Robert L Hanson, Alejandra Marinelarena, Sayuko Kobes, Yunhua L Muller, Alka Malhotra, Ke Huang, Jessica Perez, Alex Gale, et al. Map2k3 is associated with body mass index in american indians and caucasians and may mediate hypothalamic inflammation. Human molecular genetics, 22(21):4438–4449, 2013.

[37] Thomas W Winkler, Anne E Justice, Mariaelisa Graff, Llilda Barata, Mary F Feitosa, Su Chu, Jacek Czajkowski, Tõnu Esko, Tove Fall, Tuomas O Kilpeläainen, et al. The influence of age and sex on genetic associations with adult body size and shape: a large-scale genome-wide interaction study. PLoS genetics, 11(10):e1005378, 2015.

[38] Adam E Locke, Bratati Kahali, Sonja I Berndt, Anne E Justice, Tune H Pers, Felix R Day, Corey Powell, Sailaja Vedantam, Martin L Buchkovich, Jian Yang, et al. Genetic studies of body mass index yield new insights for obesity biology. Nature, 518(7538):197–206, 2015.

[39] Masato Akiyama, Yukinori Okada, Masahiro Kanai, Atsushi Takahashi, Yukihide Momozawa, Masashi Ikeda, Nakao Iwata, Shiro Ikegawa, Makoto Hirata, Koichi Matsuda, et al. Genome-wide association study identifies 112 new loci for body mass index in the japanese population. Nature genetics, 49(10):1458–1467, 2017.

[40] Steven Gazal, Hilary K Finucane, Nicholas A Furlotte, Po-Ru Loh, Pier Francesco Palamara, Xuanyao Liu, Armin Schoech, Brendan Bulik-Sullivan, Benjamin M Neale, Alexander Gusev, et al. Linkage disequilibrium–dependent architecture of human complex traits shows action of negative selection. Nature genetics, 49(10):1421–1427, 2017.

[41] Simon Myers, Leonardo Bottolo, Colin Freeman, Gil McVean, and Peter Donnelly. A fine-scale map of recombination rates and hotspots across the human genome. Science, 310(5746):321–324, 2005.

[42] Bjarni J Vilhjálmsson, Jian Yang, Hilary K Finucane, Alexander Gusev, Sara Lindstroäm, Stephan Ripke, Giulio Genovese, Po-Ru Loh, Gaurav Bhatia, Ron Do, et al. Modeling linkage disequilibrium increases accuracy of polygenic risk scores. The american journal of human genetics, 97(4):576–592, 2015.

[43] Jiashun Xiao, Mingxuan Cai, Xianghong Hu, Xiang Wan, Gang Chen, and Can Yang. Xpxp: Improving polygenic prediction by cross-population and cross-phenotype analysis. Bioinformatics, 2022.

[44] Sheng Yang and Xiang Zhou. Accurate and scalable construction of polygenic scores in large biobank data sets. The American Journal of Human Genetics, 106(5):679–693, 2020.

[45] Hilary K Finucane, Brendan Bulik-Sullivan, Alexander Gusev, Gosia Trynka, Yakir Reshef, Po-Ru Loh, Verneri Anttila, Han Xu, Chongzhi Zang, Kyle Farh, et al. Partitioning heritability by functional annotation using genome-wide association summary statistics. Nature genetics, 47(11):1228–1235, 2015.

[46] Joseph K Pickrell. Joint analysis of functional genomic data and genome-wide association studies of 18 human traits. The American Journal of Human Genetics, 94(4):559–573, 2014.

[47] Can Yang, Xiang Wan, Xinyi Lin, Mengjie Chen, Xiang Zhou, and Jin Liu. Comm: a collaborative mixed model to dissecting genetic contributions to complex traits by leveraging regulatory information. Bioinformatics, 35(10):1644–1652, 2019.

[48] Mingxuan Cai, Lin S Chen, Jin Liu, and Can Yang. Igrex for quantifying the impact of genetically regulated expression on phenotypes. NAR genomics and bioinformatics, 2(1):lqaa010, 2020.

[49] Xingjie Shi, Xiaoran Chai, Yi Yang, Qing Cheng, Yuling Jiao, Haoyue Chen, Jian Huang, Can Yang, and Jin Liu. A tissue-specific collaborative mixed model for jointly analyzing multiple tissues in transcriptome-wide association studies. Nucleic acids research, 48(19):e109–e109, 2020.

[50] Anshul Kundaje, Wouter Meuleman, Jason Ernst, Misha Bilenky, Angela Yen, Alireza Heravi-Moussavi, Pouya Kheradpour, Zhizhuo Zhang, Jianrong Wang, Michael J Ziller, et al. Integrative analysis of 111 reference human epigenomes. Nature, 518(7539):317–330, 2015.

[51] Qiongshi Lu, Boyang Li, Derek Ou, Margret Erlendsdottir, Ryan L Powles, Tony Jiang, Yiming Hu, David Chang, Chentian Jin, Wei Dai, et al. A powerful approach to estimating annotation-stratified genetic covariance via gwas summary statistics. The American Journal of Human Genetics, 101(6):939–964, 2017.

[52] Xiang Zhu and Matthew Stephens. Large-scale genome-wide enrichment analyses identify new trait-associated genes and pathways across 31 human phenotypes. Nature communications, 9(1):1–14, 2018.

[53] Dongjun Chung, Can Yang, Cong Li, Joel Gelernter, and Hongyu Zhao. Gpa: a statistical approach to prioritizing gwas results by integrating pleiotropy and annotation. PLoS genetics, 10(11):e1004787, 2014.

[54] Jingsi Ming, Mingwei Dai, Mingxuan Cai, Xiang Wan, Jin Liu, and Can Yang. Lsmm: a statistical approach to integrating functional annotations with genome-wide association studies. Bioinformatics, 34(16):2788–2796, 2018.

[55] Gleb Kichaev, Gaurav Bhatia, Po-Ru Loh, Steven Gazal, Kathryn Burch, Malika K Freund, Armin Schoech, Bogdan Pasaniuc, and Alkes L Price. Leveraging polygenic functional enrichment to improve gwas power. The American Journal of Human Genetics, 104(1):65–75, 2019.

[56] Jingsi Ming, Tao Wang, and Can Yang. Lpm: a latent probit model to characterize the relationship among complex traits using summary statistics from multiple gwass and functional annotations. Bioinformatics, 36(8):2506–2514, 2020.

[57] Gleb Kichaev, Wen-Yun Yang, Sara Lindstrom, Farhad Hormozdiari, Eleazar Eskin, Alkes L Price, Peter Kraft, and Bogdan Pasaniuc. Integrating functional data to prioritize causal variants in statistical fine-mapping studies. PLoS genetics, 10(10):e1004722, 2014.

[58] Yue Li and Manolis Kellis. Joint bayesian inference of risk variants and tissue-specific epigenomic enrichments across multiple complex human diseases. Nucleic acids research, 44(18):e144–e144, 2016.

[59] Yiming Hu, Qiongshi Lu, Ryan Powles, Xinwei Yao, Can Yang, Fang Fang, Xinran Xu, and Hongyu Zhao. Leveraging functional annotations in genetic risk prediction for human complex diseases. PLoS computational biology, 13(6):e1005589, 2017.

[60] Carla Márquez-Luna, Steven Gazal, Po-Ru Loh, Samuel S Kim, Nicholas Furlotte, Adam Auton, and Alkes L Price. Incorporating functional priors improves polygenic prediction accuracy in uk biobank and 23andme data sets. Nature Communications, 12(1):1–11, 2021.

[61] Tiffany Amariuta, Kazuyoshi Ishigaki, Hiroki Sugishita, Tazro Ohta, Masaru Koido, Kushal K Dey, Koichi Matsuda, Yoshinori Murakami, Alkes L Price, Eiryo Kawakami, et al. Improving the trans-ancestry portability of polygenic risk scores by prioritizing variants in predicted cell-type-specific regulatory elements. Nature genetics, 52(12):1346–1354, 2020.

[62] Can Yang, Cong Li, Qian Wang, Dongjun Chung, and Hongyu Zhao. Implications of pleiotropy: challenges and opportunities for mining big data in biomedicine. Frontiers in genetics, 6:229, 2015.

[63] Doug Speed, Gibran Hemani, Michael R Johnson, and David J Balding. Improved heritability estimation from genome-wide SNPs. The American Journal of Human Genetics, 91(6):1011–1021, 2012.

[64] Li Yang, Benjamin M Neale, Lu Liu, S Hong Lee, Naomi R Wray, Ning Ji, Haimei Li, Qiujin Qian, Dongliang Wang, Jun Li, et al. Polygenic transmission and complex neuro developmental network for attention deficit hyperactivity disorder: Genome-wide association study of both common and rare variants. American Journal of Medical Genetics Part B: Neuropsychiatric Genetics, 162(5):419–430, 2013.

[65] Jian Zeng, Ronald De Vlaming, Yang Wu, Matthew R Robinson, Luke R Lloyd-Jones, Loic Yengo, Chloe X Yap, Angli Xue, Julia Sidorenko, Allan F McRae, et al. Signatures of negative selection in the genetic architecture of human complex traits. Nature genetics, 50(5):746–753, 2018.

[66] Doug Speed, John Holmes, and David J Balding. Evaluating and improving heritability models using summary statistics. Nature Genetics, 52(4):458–462, 2020.

[67] Loic Yengo, Jian Yang, and Peter M Visscher. Expectation of the intercept from bivariate ld score regression in the presence of population stratification. bioRxiv, page 310565, 2018.

[68] Alastair R Hall et al. Generalized method of moments. Oxford university press, 2005.

