## Supplementary for "Leveraging the local genetic structure for trans-ancestry association mapping"

#### Contents

**1 Supplementary Figures** **2**

**2 Supplementary Tables** **24**

### 1 Supplementary Figures

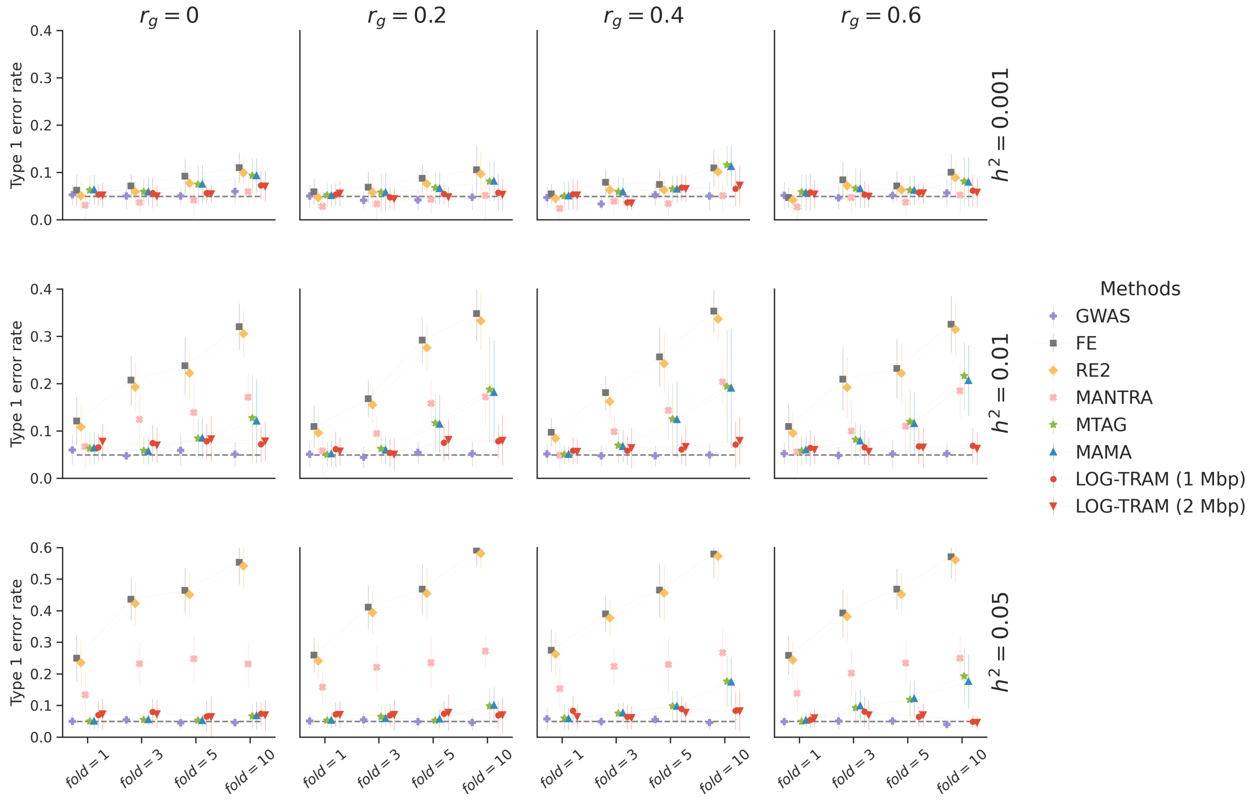


Figure S1: Comparisons among LOG-TRAM, MAMA, MTAG, MANTRA, RE2, FE, and GWAS in simulation studies of 10% SNPs with shared non-zero effects. Average type I error rates were evaluated under different settings of background trans-ancestry genetic correlations ($r_g$), fold enrichment ($fold$) of EUR local heritability and total heritability ($h^2$) of the whole chromosome. Error bars represent the standard errors of type I error rates evaluated on 20 replications.

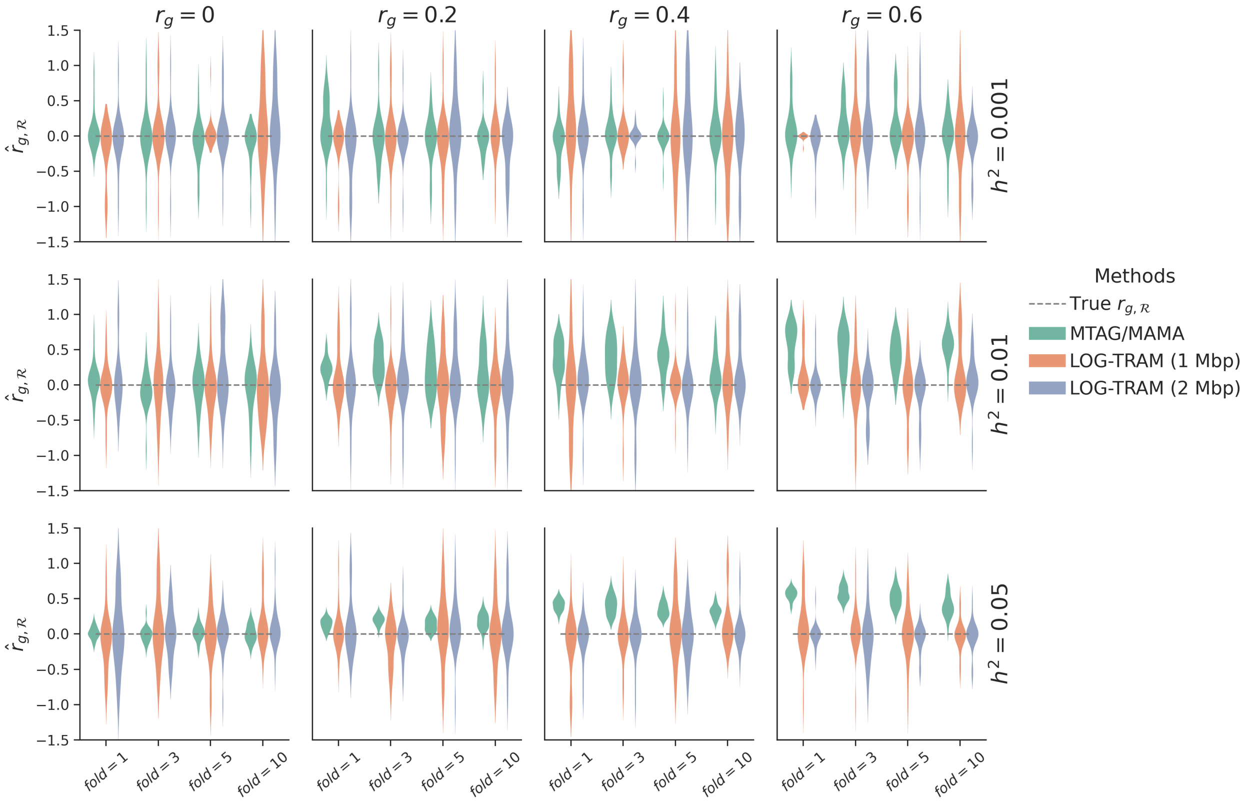


Figure S2: Comparisons among LOG-TRAM, MAMA and MTAG in simulation studies of 10% SNPs with shared non-zero effects. Local trans-ancestry genetic correlations ($\hat{r}_{g,\boldsymbol{\mathcal{R}}}$) were estimated under different settings of background trans-ancestry genetic correlations ($r_g$), fold enrichment ($fold$) of EUR local heritability, and total heritability ($h^2$) of the whole chromosome. The results were summarized over 20 replications. For GWAS meta-analysis methods with the homogeneous assumption (e.g., MTAG and MAMA), they used the global correlation $\hat{r}_g$ as the local correlation $\hat{r}_{g,\boldsymbol{\mathcal{R}}}$. Hence, their estimates were often biased.

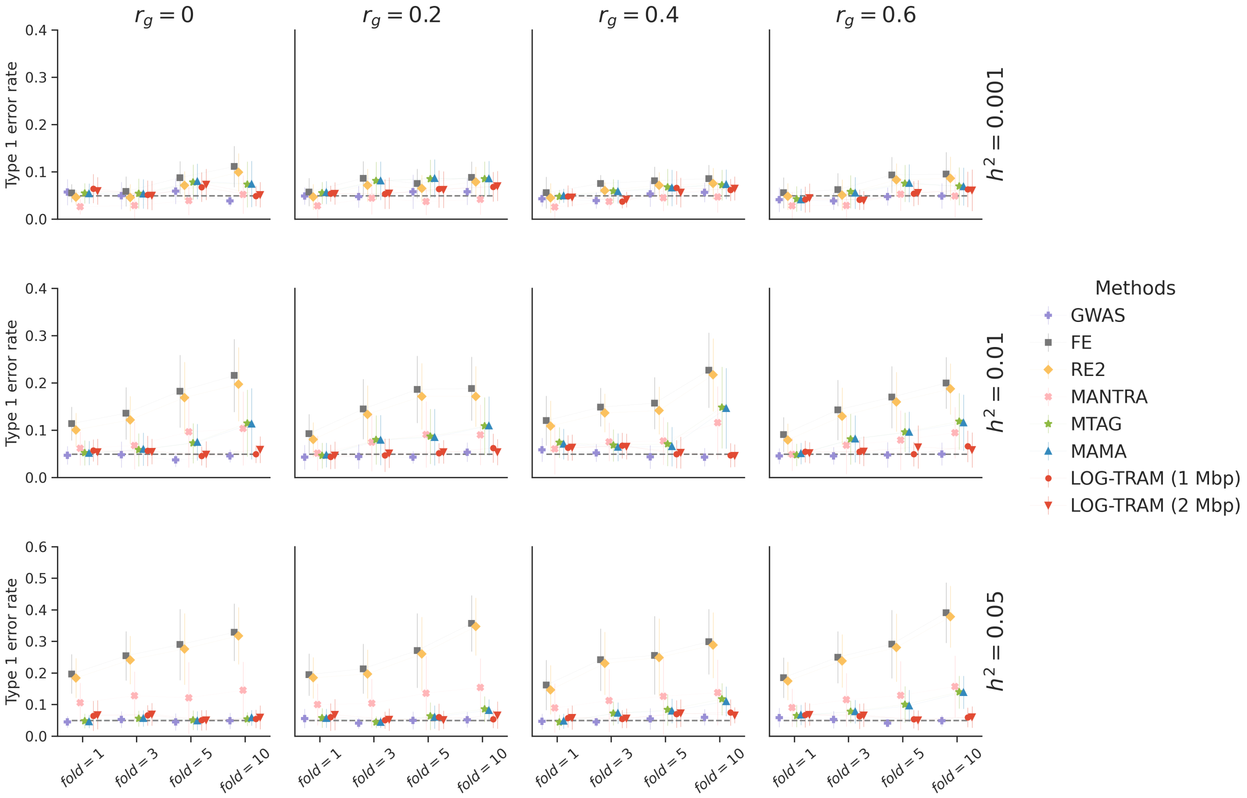


Figure S3: Comparisons among LOG-TRAM, MAMA, MTAG, MANTRA, RE2, FE, and GWAS in simulation studies of 1% SNPs with shared non-zero effects. Average type I error rates were evaluated under different settings of background trans-ancestry genetic correlations ($r_g$), fold enrichment ($fold$) of EUR local heritability, and total heritability ($h^2$) of the whole chromosome. Error bars represent the standard errors of type I error rates evaluated on 20 replications.

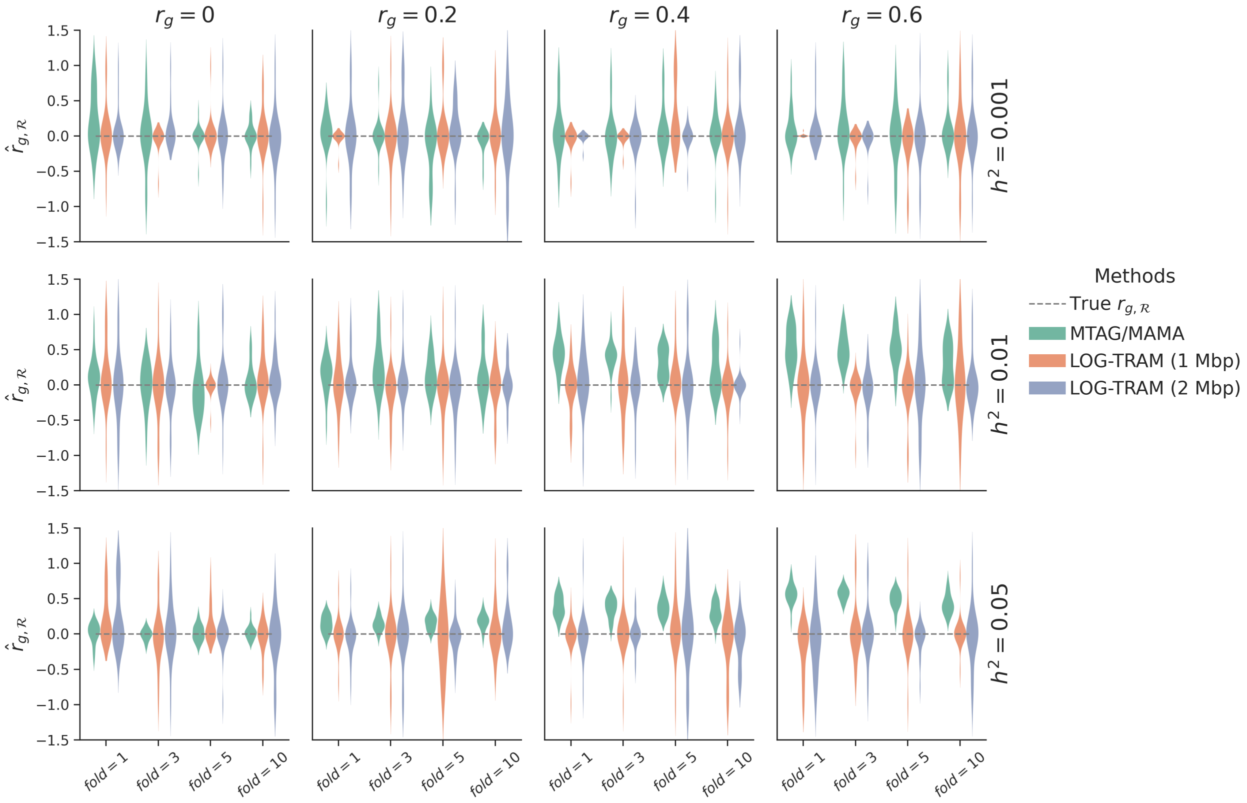


Figure S4: Comparisons among LOG-TRAM, MAMA, and MTAG in simulation studies of 1% SNPs with shared non-zero effects. Estimated local trans-ancestry genetic correlations ($\hat{r}_{g,\mathcal{R}}$) under different setting of background trans-ancestry genetic correlations ($r_g$), fold enrichment ($fold$) of EUR local heritability and total heritability across the chromosome ($h^2$). The results were summarized over 20 replications. For GWAS meta-analysis methods with the homogeneous assumption (e.g., MTAG and MAMA), they used the global correlation $\hat{r}_g$ as the local correlation $\hat{r}_{g,\mathcal{R}}$. Hence, their estimates were often biased.

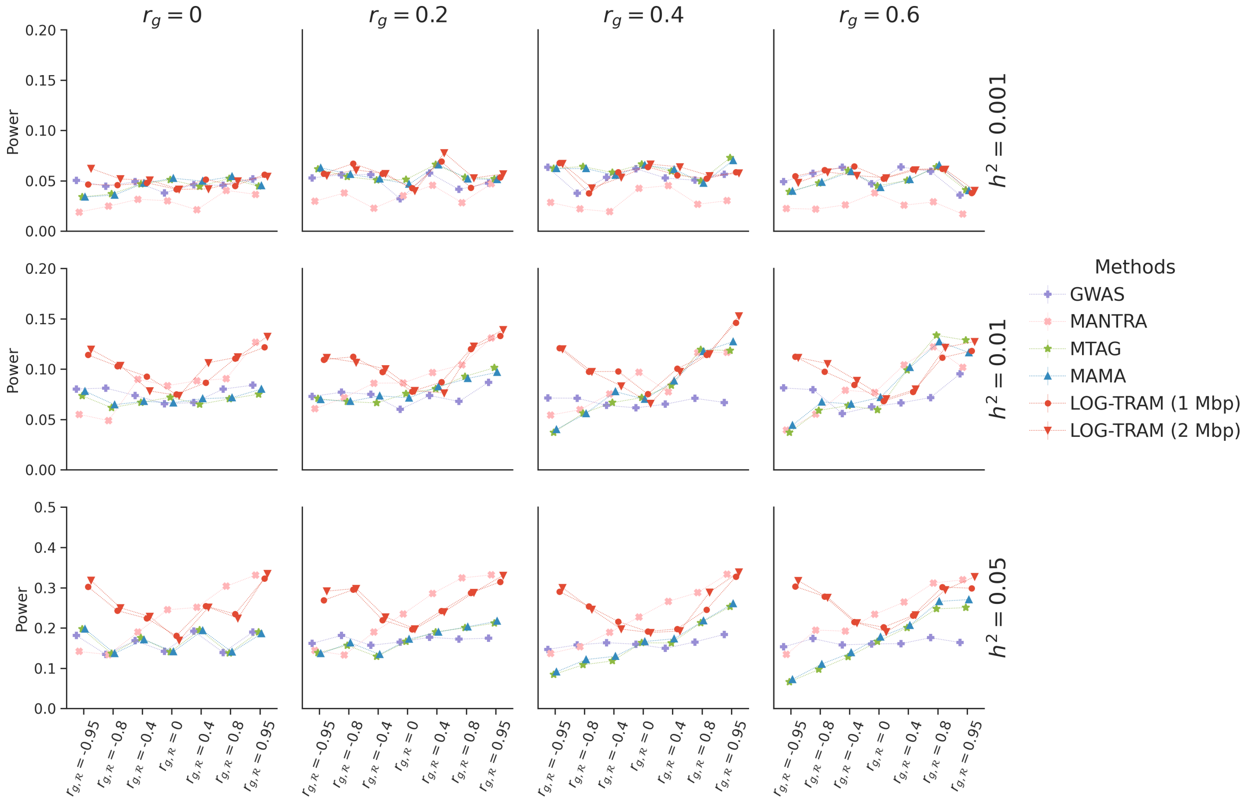


Figure S5: Comparisons among LOG-TRAM, MAMA, MTAG, MANTRA, RE2, FE, and GWAS in simulation studies of 10% SNPs with shared non-zero effects. Averaged statistical power were evaluated in multiple simulation scenarios with different combinations of background trans-ancestry genetic correlations ($r_g$), local trans-ancestry genetic correlations ($r_{g,\mathcal{R}}$) and total heritability ($h^2$) of the whole chromosome. Error bars represent the standard errors of type I error rate evaluated on 20 replications.

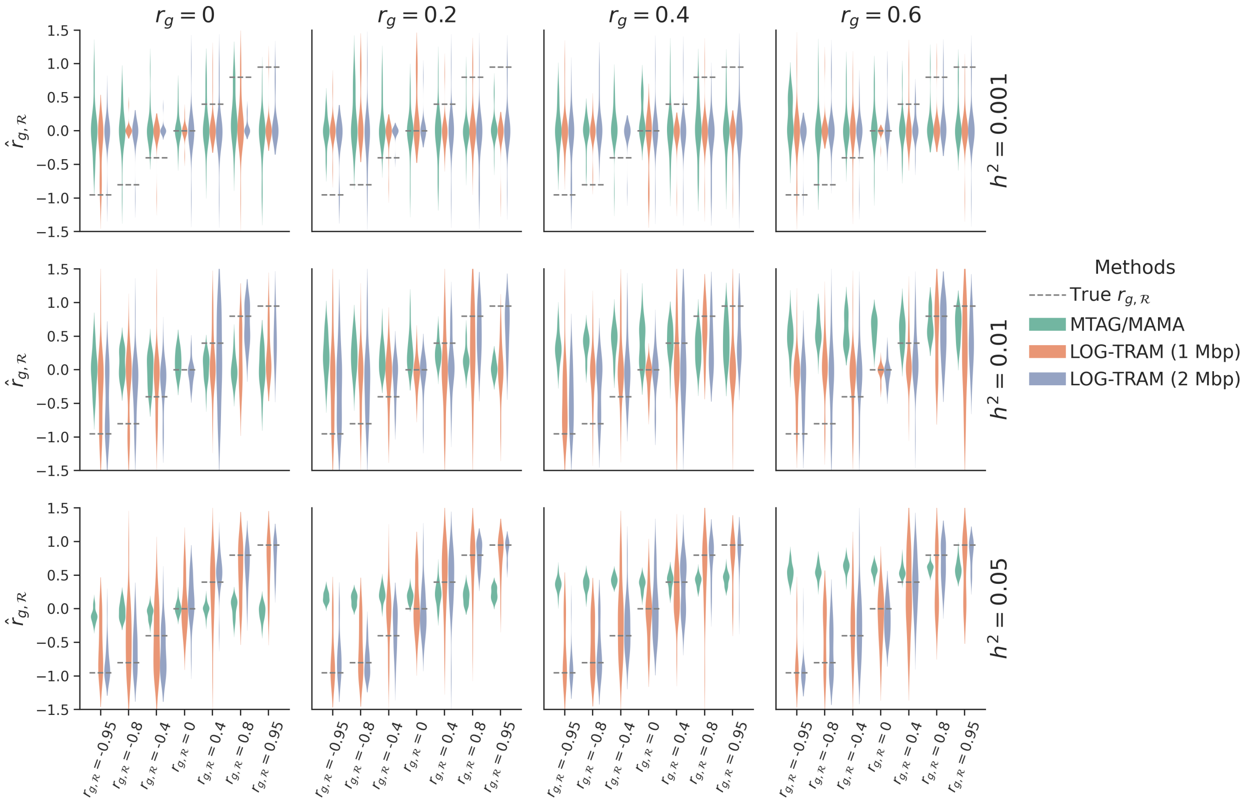


Figure S6: Comparisons among LOG-TRAM, MAMA, and MTAG in simulation studies of 10% SNPs with shared non-zero effects. Local trans-ancestry genetic correlations ($\hat{r}_{g,\boldsymbol{\mathcal{R}}}$) were estimated under different settings of background trans-ancestry genetic correlations ($r_g$), local trans-ancestry genetic correlations ($r_{g,\boldsymbol{\mathcal{R}}}$) and total heritability ($h^2$) of the whole chromosome. The results were summarized over 20 replications. For GWAS meta-analysis methods with the homogeneous assumption (e.g., MTAG and MAMA), they used the global correlation $\hat{r}_g$ as the local correlation $\hat{r}_{g,\boldsymbol{\mathcal{R}}}$. Hence, their estimates were often biased.

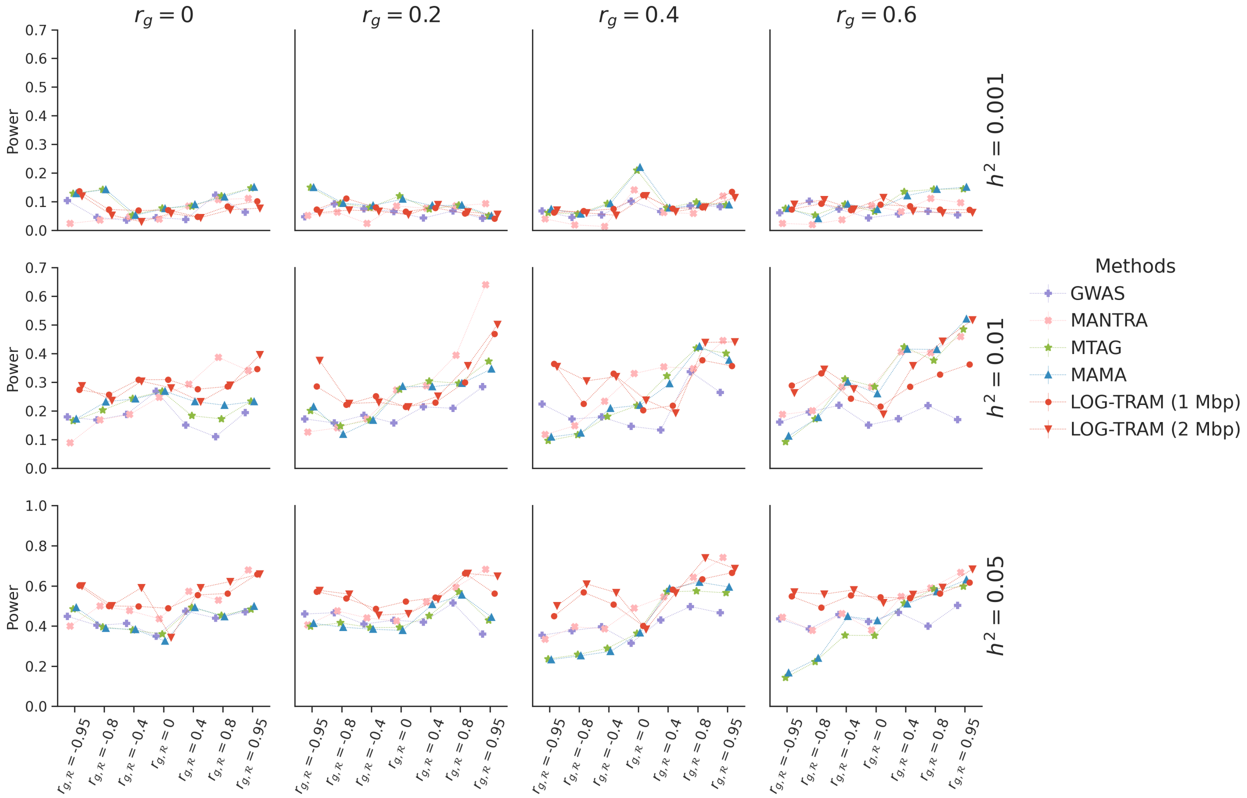


Figure S7: Comparisons among LOG-TRAM, MAMA, MTAG, MANTRA, RE2, FE, and GWAS in simulation studies of 1% SNPs with shared non-zero effects. Average statistical power were evaluated in multiple simulation scenarios with different combinations of background trans-ancestry genetic correlations ($r_g$), local trans-ancestry genetic correlations ($r_{g,\boldsymbol{\mathcal{R}}}$) and total heritability ($h^2$) of the whole chromosome. Error bars represent the standard errors of type I error rate evaluated on 20 replications.

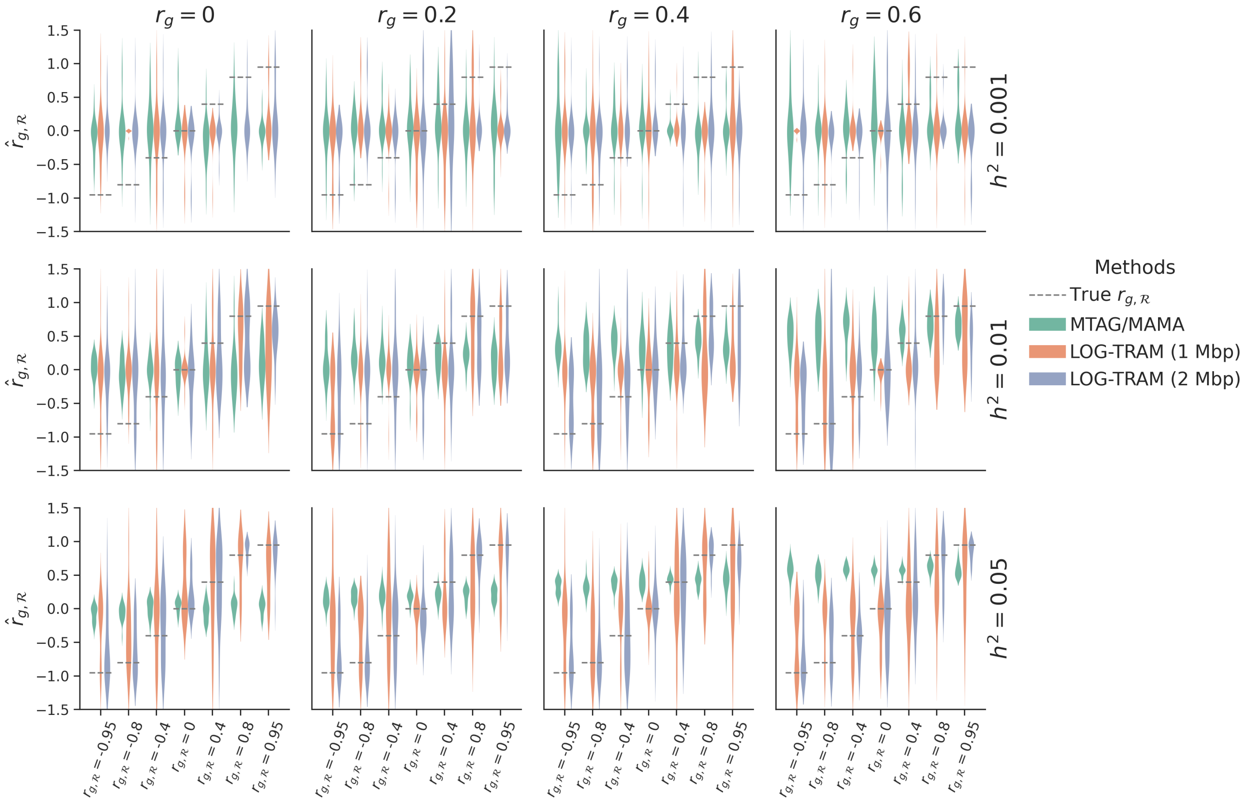


Figure S8: Comparisons among LOG-TRAM, MAMA, and MTAG in simulation studies of 1% SNPs with shared non-zero effects. Local trans-ancestry genetic correlations ($\hat{r}_{g,\mathcal{R}}$) were estimated under different settings of background trans-ancestry genetic correlations ($r_g$), local trans-ancestry genetic correlations ($r_{g,\mathcal{R}}$) and total heritability ($h^2$) of the whole chromosome. The results were summarized over 20 replications. For GWAS meta-analysis methods with the homogeneous assumption (e.g., MTAG and MAMA), they used the global correlation $\hat{r}_g$ as the local correlation $\hat{r}_{g,\mathcal{R}}$. Hence, their estimates were often biased.

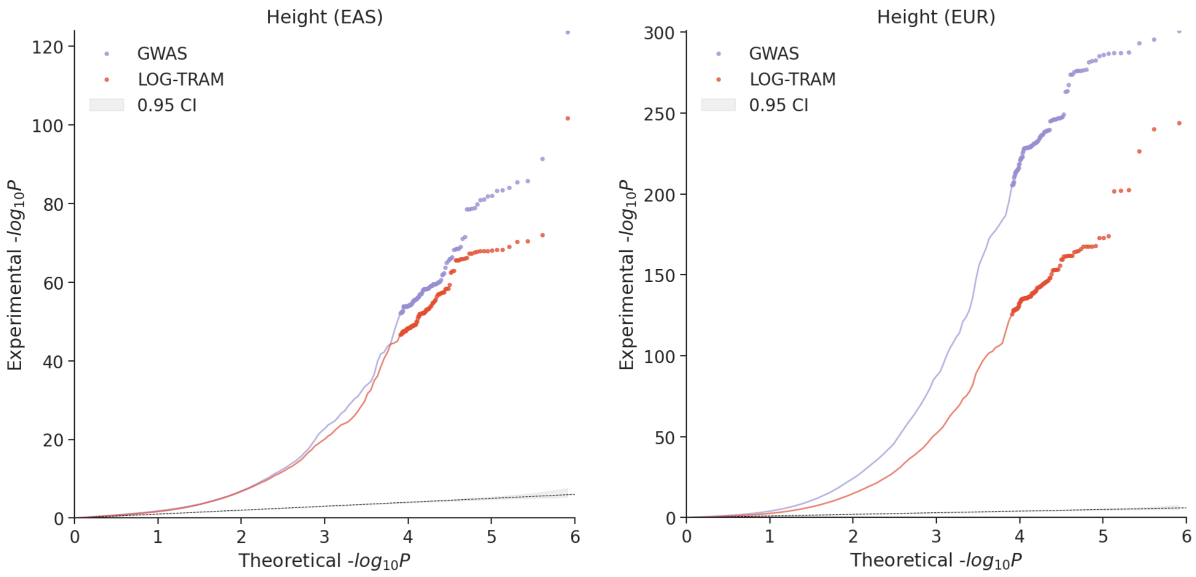


Figure S9: QQ plots of GWAS and LOG-TRAM results for height from EAS (left) and EUR (right). LOG-TRAM significantly reduces the hidden confounding bias in the standard GWAS results.

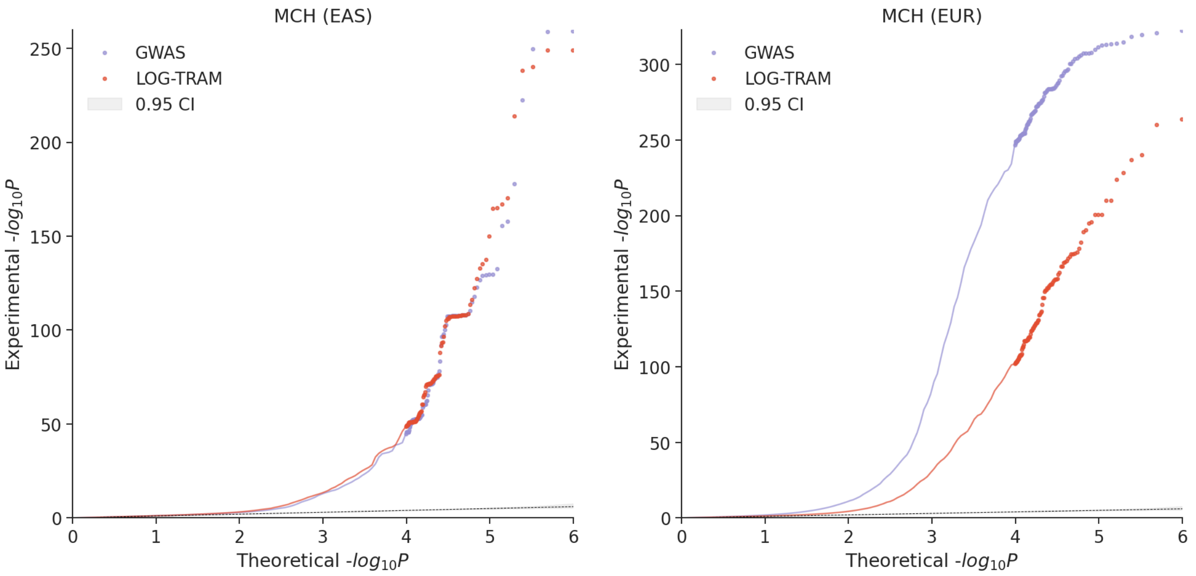


Figure S10: QQ plots of GWAS and LOG-TRAM results for MCH from EAS (left) and EUR (right). LOG-TRAM significantly reduces the hidden confounding bias in the standard EUR GWAS results.

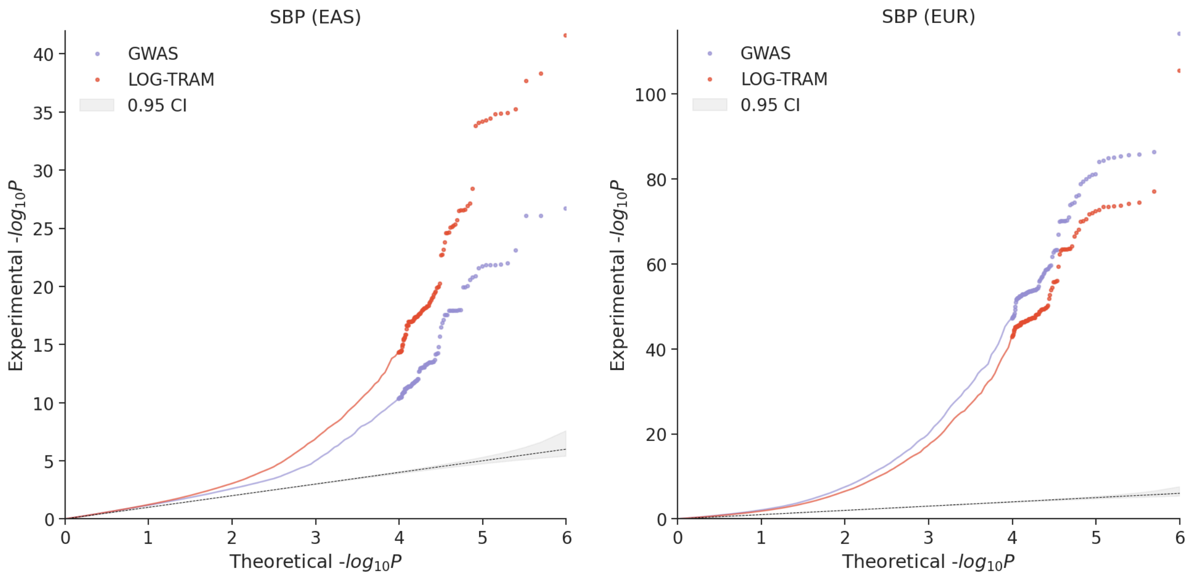


Figure S11: QQ plots of GWAS and LOG-TRAM results for SBP from EAS (left) and EUR (right). LOG-TRAM significantly reduces the hidden confounding bias in the standard EUR GWAS results. Furthermore, LOG-TRAM achieves substantial power gain in EAS.

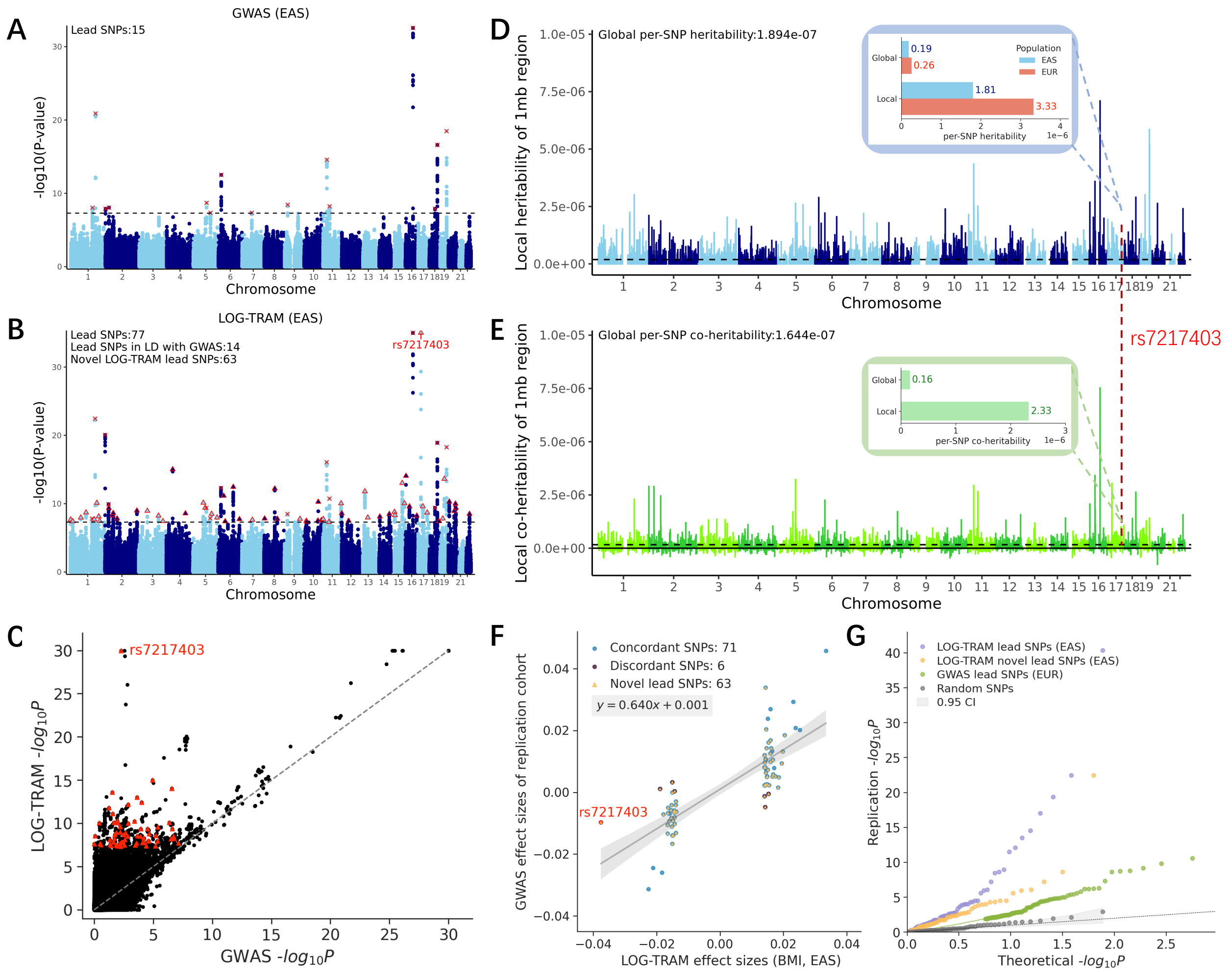


Figure S12: Trans-ancestry association results of the BMI. **A**: Manhattan plot of the BBJ females, where 15 independent lead SNPs were identified. **B**: Manhattan plot of BBJ female obtained by LOG-TRAM with 1 Mbp window size. LOG-TRAM identified 77 independent lead SNPs, of which 63 were not identified in the original BBJ female GWAS. Lead SNPs identified by both LOG-TRAM and GWAS are marked by a cross. Novel lead SNPs are marked by a triangle. **C**: Comparison of *p*-values output by LOG-TRAM and original BMI GWAS *p*-values of the BBJ females. **D**: Local per-SNP heritability estimated by LOG-TRAM in the BBJ females and UKBB. **E**: Local per-SNP co-heritability between the BBJ females and UKBB. **F**: Comparison of lead SNPs effect sizes estimated by LOG-TRAM and effect sizes of GWAS in the replication dataset (BBJ males). **G**: The QQ plot compares the *p*-values in the replication GWAS data: (1) LOG-TRAM lead SNPs, (2) LOG-TRAM novel lead SNPs, (3) Lead SNPs identified from UKBB GWAS, and (4) Randomly selected SNPs from the replication GWAS.

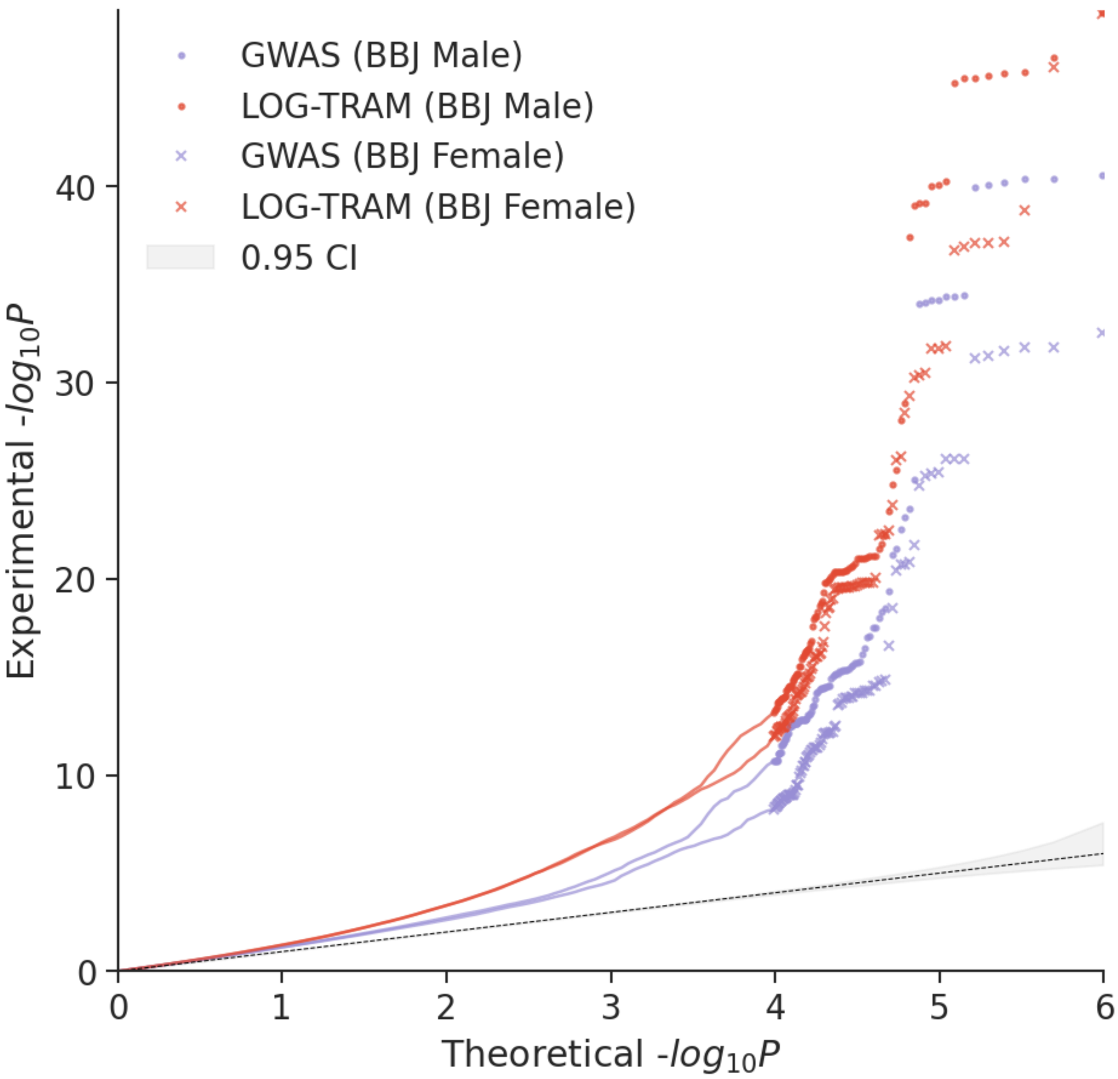


Figure S13: QQ plots of GWAS and LOG-TRAM results for BMI. LOG-TRAM achieves substantial power gain compared to the standard sex-stratified GWASs.

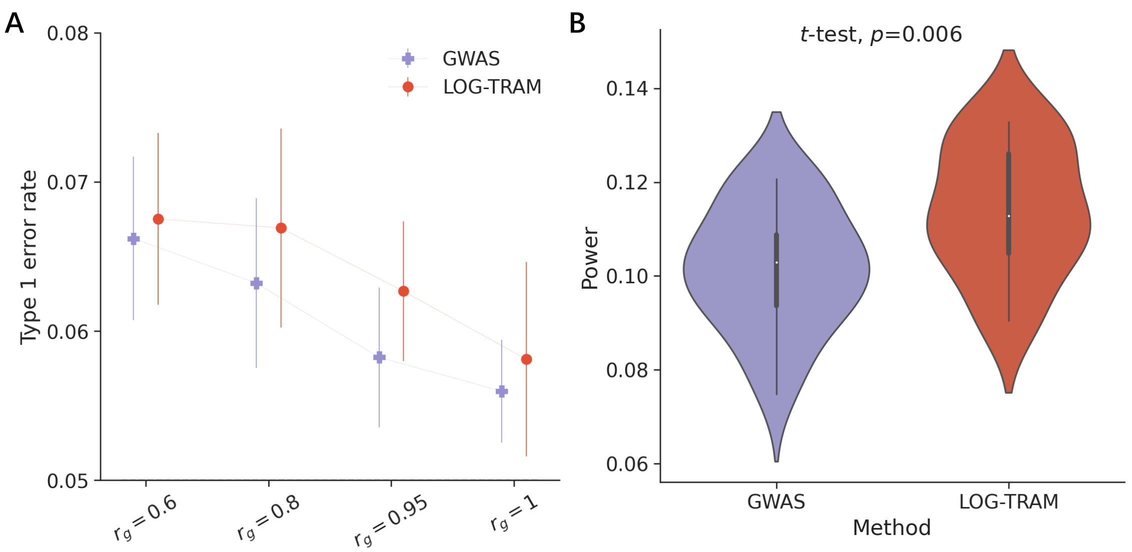


Figure S14: Comparison of the GWAS-based and LOG-TRAM-based difference tests in simulation studies. **A**: Average type I error rates evaluated under different settings of trans-ancestry genetic correlations ($r_g$). Error bars represent the standard errors of type I error rates evaluated on 20 replications. We considered 18K samples from the EAS cohort and 100K samples from UKBB as the target and auxiliary populations, respectively. We used their genotype matrices to mimic the different LD patterns and allele frequencies between populations. In our simulation study, 17,248 HapMap3 matched SNPs from chromosome 20 were used. To generate null SNPs (SNPs with the same effect size across the two populations), we simulated the shared effects ($\boldsymbol{\beta}$) from a bivariate normal distribution $\mathcal{N}\left(\mathbf{0}, \begin{bmatrix} h_1^2 & r_g h_1 h_2 \\ r_g h_1 h_2 & h_2^2 \end{bmatrix} /0.1M\right)$, where $r_g$ is the trans-ancestry genetic correlation, $h_1^2 = h_2^2 = 0.01$ means that the total heritability of the whole chromosome is 0.01, $M$ is number of SNPs, and 0.1 represents that 10% of SNPs with non-zero effects jointly contribute to the local heritability. We varied $r_g$ at $\{0.6, 0.8, 0.95, 1\}$. Given the effect sizes and genotype matrices, quantitative phenotypes in both populations were simulated using Eq. (3) in the main text. After that, we marginally regressed the simulated phenotypes on each SNP to obtain the $z$-scores of the two datasets. Finally, after performing meta-analysis, we reported the fraction of SNPs with a $p$-value less than 0.05 as the type 1 error rate. As expected, with the effect size getting more and more similar ($r_g$ increased from 0.6 to 1), the type 1 error rates of both GWAS-based and LOG-TRAM-based difference tests reduced to nearly 0.05. However, the heterogeneity of LD patterns across ancestries results in different marginal effect sizes ($\mathbf{b}$) even though the underlying true effect sizes ($\boldsymbol{\beta}$) are identical between the two populations. Therefore, both marginal effect sizes-based difference tests have type 1 error rates slightly larger than 0.05 when $r_g = 1$. **B**: Violin plots of statistical power evaluated over 20 replications. Similarly, we considered a polygenic scenario by setting 10% SNPs with shared non-zero effects and generated alternative SNPs (SNPs with different effect size across the two populations) using a bivariate normal distribution $\mathcal{N}\left(\mathbf{0}, \begin{bmatrix} h_1^2 & 0 \\ 0 & h_2^2 \end{bmatrix} /0.1M\right)$ to simulate $\boldsymbol{\beta}$. We simulated the quantitative phenotypes and obtained the $z$-scores of the two datasets. Finally, after performing meta-analysis, we reported the fraction of non-zero SNPs with a $p$-value less than 0.05 as the power. We observed that LOG-TRAM-based difference test have significantly higher power ($p$-value=0.006) than GWAS-based test.

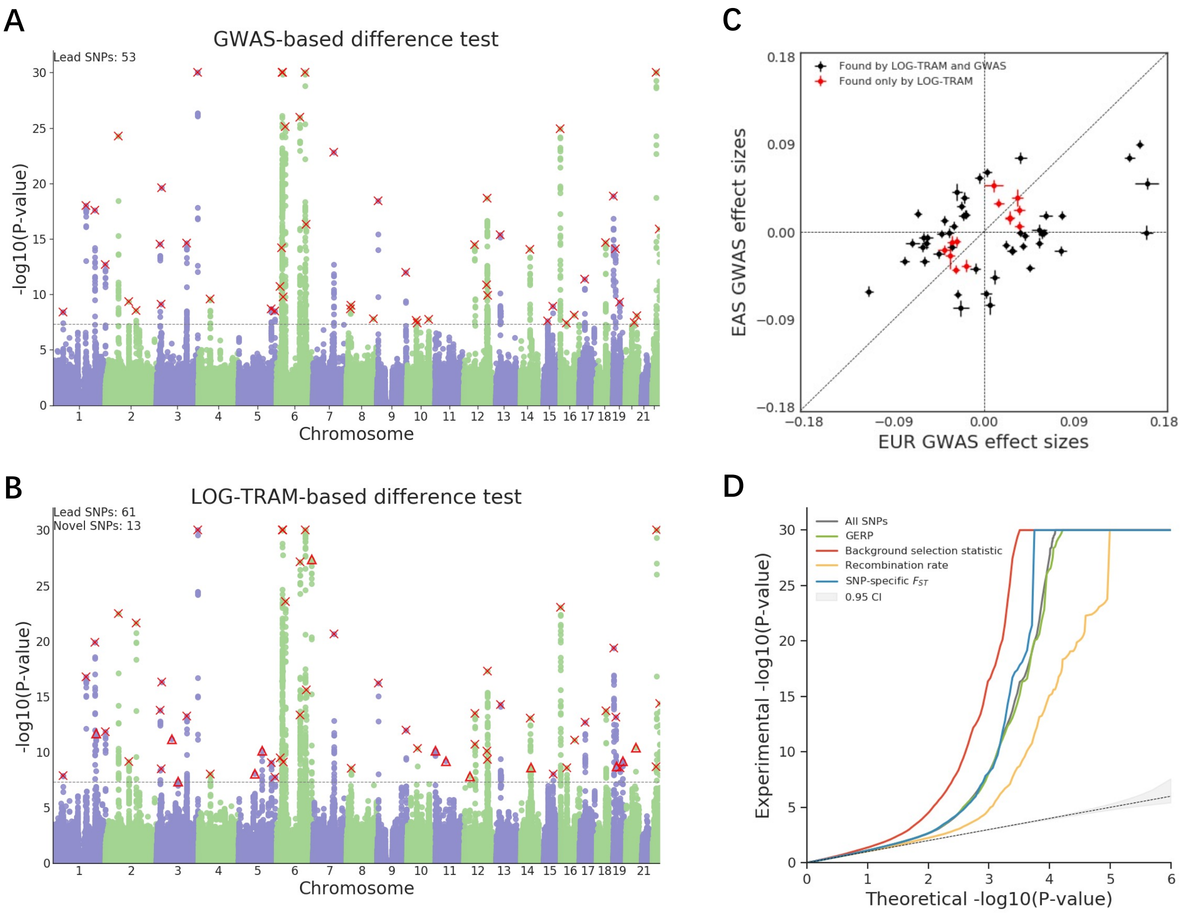


Figure S15: Ancestry-specific loci identification for MCV. **A**: Manhattan plot of the GWAS-based difference test for MCV. Lead SNPs are marked by a cross. **B**: Manhattan plot of the LOG-TRAM-based difference test for MCV. The dashed line marks the threshold for genome-wide significance ($P = 5 \times 10^{-8}$). Lead SNPs identified by both LOG-TRAM and GWAS-based difference test are marked by crosses. Novel lead SNPs are marked by triangles. **C**: Comparison of MCV GWAS effect sizes for lead SNPs between EAS and EUR. Red points denote novel loci identified by the LOG-TRAM-based test; black points denote lead loci identified in both the GWAS and LOG-TRAM-based difference tests. Vertical and horizontal error bars represent standard errors of GWAS effect sizes in EAS and EUR, respectively. **D** The QQ-plots of $p$-values for all SNPs and SNPs in the top quantile of four continuous-valued annotations.

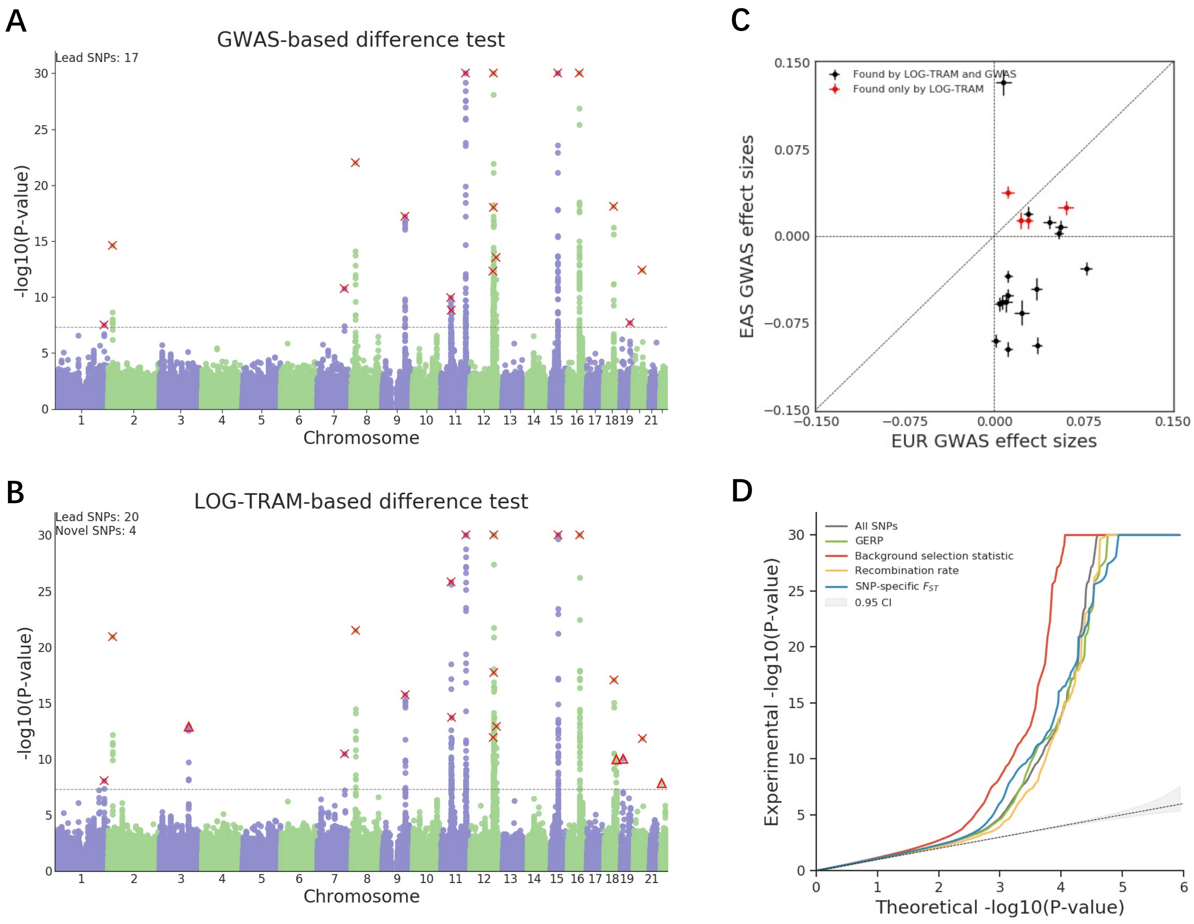


Figure S16: Ancestry-specific loci identification for HDL. **A**: Manhattan plot of the GWAS-based difference test for HDL. Lead SNPs are marked by crosses. **B**: Manhattan plot of the LOG-TRAM-based difference test for HDL. The dashed line marks the threshold for genome-wide significance ($P = 5 \times 10^{-8}$). Lead SNPs identified by both LOG-TRAM and GWAS-based difference test are marked by crosses. Novel lead SNPs are marked by triangles. **C**: Comparison of HDL GWAS effect sizes for lead SNPs between EAS and EUR. Red points denote novel loci identified by the LOG-TRAM-based test; black points denote lead loci identified in both the GWAS and LOG-TRAM-based difference tests. Vertical and horizontal error bars represent standard errors of GWAS effect sizes in EAS and EUR, respectively. **D** The QQ-plots of $p$-values for all SNPs and SNPs in the top quantile of four continuous-valued annotations.

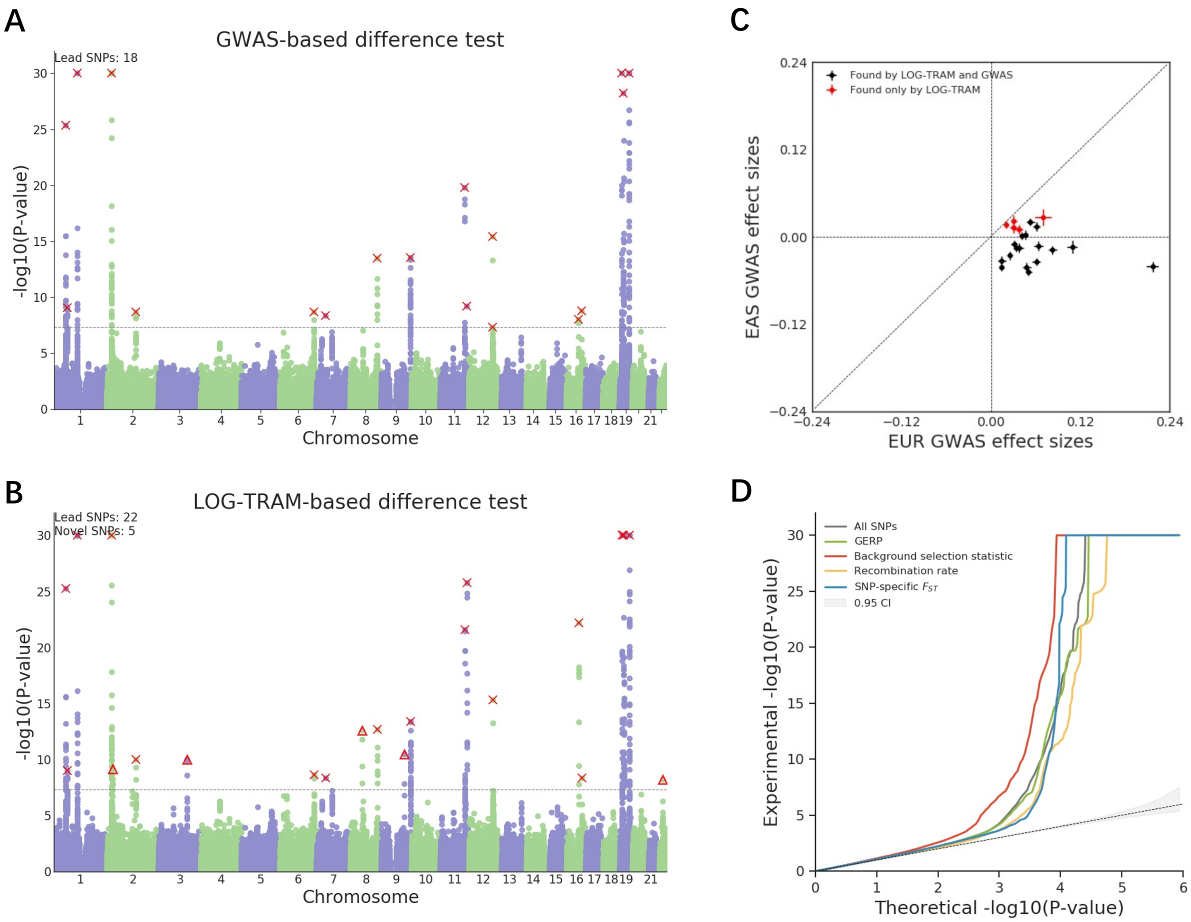


Figure S17: Ancestry-specific loci identification for LDL. **A**: Manhattan plot of the GWAS-based difference test for LDL. Lead SNPs are marked by a cross. **B**: Manhattan plot of the LOG-TRAM-based difference test for LDL. The dashed line marks the threshold for genome-wide significance ($P = 5 \times 10^{-8}$). Lead SNPs identified by both LOG-TRAM and GWAS-based difference test are marked by crosses. Novel lead SNPs are marked by triangles. **C**: Comparison of LDL GWAS effect sizes for lead SNPs between EAS and EUR. Red points denote novel loci identified by the LOG-TRAM-based test; black points denote lead loci identified in both the GWAS and LOG-TRAM-based difference tests. Vertical and horizontal error bars represent standard errors of GWAS effect sizes in EAS and EUR, respectively. **D** The QQ-plots of $p$-values for all SNPs and SNPs in the top quantile of four continuous-valued annotations.

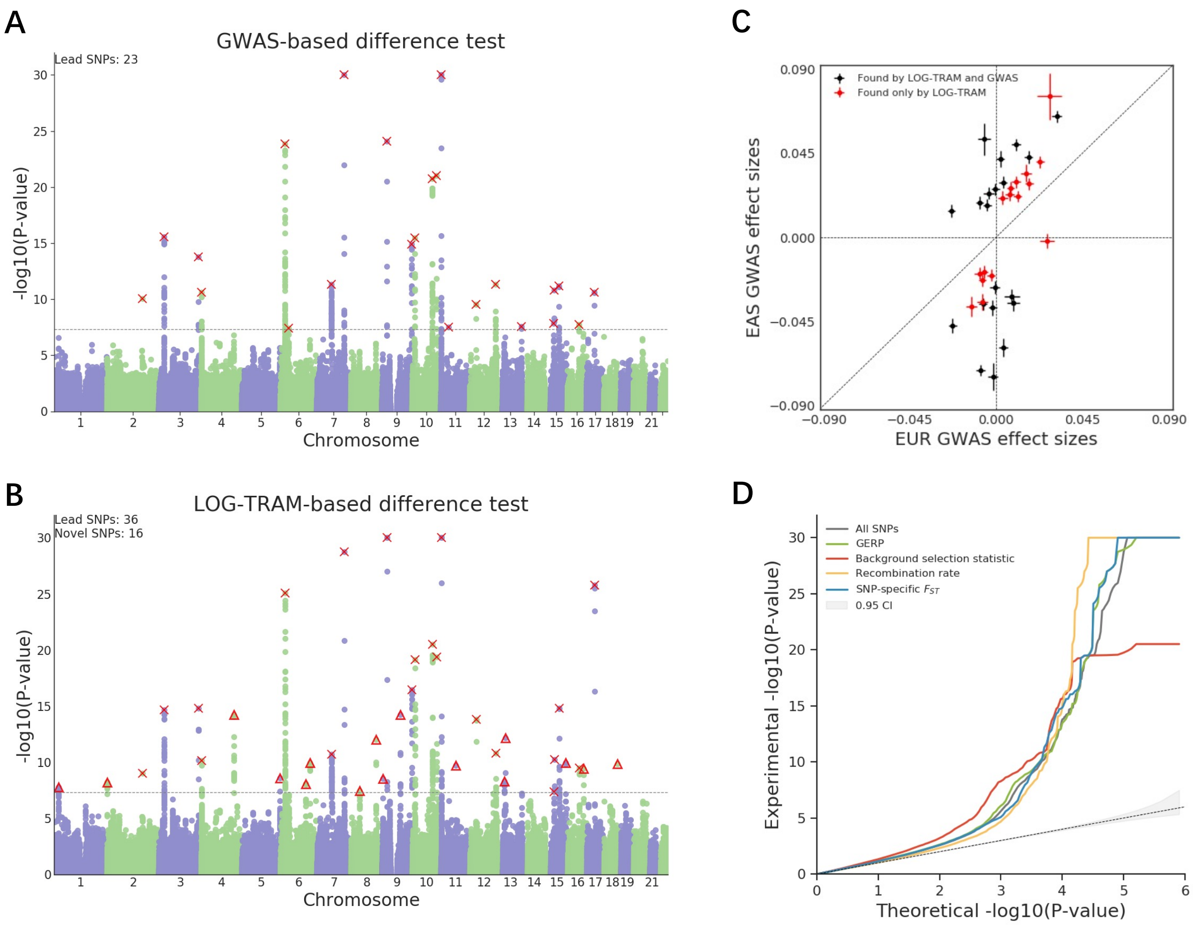


Figure S18: Ancestry-specific loci identification for T2D. **A**: Manhattan plot of the GWAS-based difference test for T2D. Lead SNPs are marked by crosses. **B**: Manhattan plot of the LOG-TRAM-based difference test for T2D. The dashed line marks the threshold for genome-wide significance ($P = 5 \times 10^{-8}$). Lead SNPs identified by both the LOG-TRAM and GWAS-based difference tests are marked by crosses. Novel lead SNPs are marked by triangles. **C**: Comparison of T2D GWAS effect sizes for lead SNPs between EAS and EUR. Red points denote novel loci identified by the LOG-TRAM-based test; black points denote lead loci identified in both the GWAS and LOG-TRAM-based difference tests. Vertical and horizontal error bars represent standard errors of GWAS effect sizes in EAS and EUR, respectively. **D** The QQ-plots of $p$-values for all SNPs and SNPs in the top quantile of four continuous-valued annotations.

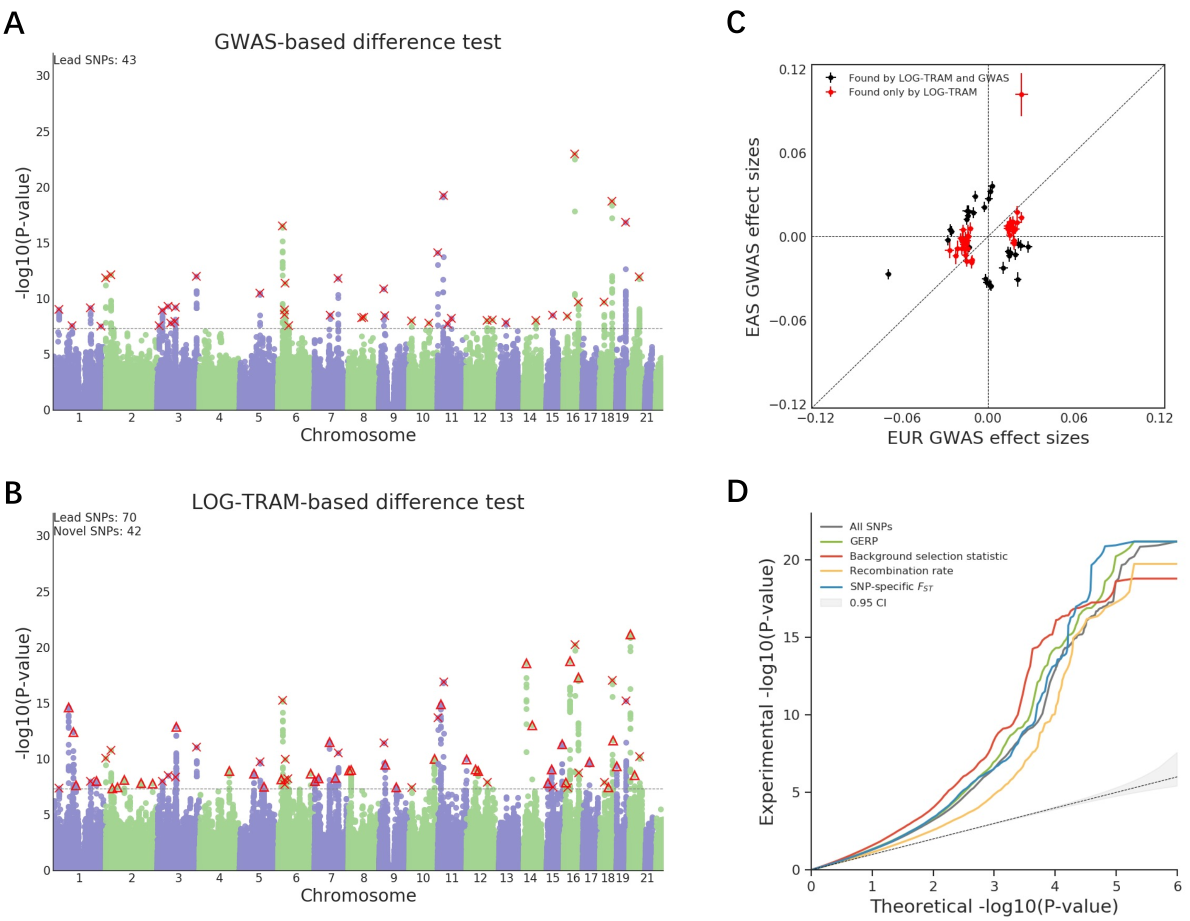


Figure S19: Ancestry-specific loci identification for BMI. **A**: Manhattan plot of the GWAS-based difference test for BMI. Lead SNPs are marked by crosses. **B**: Manhattan plot of the LOG-TRAM-based difference test for BMI. The dashed line marks the threshold for genome-wide significance ($P = 5 \times 10^{-8}$). Lead SNP identified by both the LOG-TRAM and GWAS-based difference tests are marked by crosses. Novel lead SNPs are marked by triangles. **C**: Comparison of BMI GWAS effect sizes for lead SNPs between EAS and EUR. Red points denote novel loci identified by the LOG-TRAM-based test; black points denote lead loci identified in both the GWAS and LOG-TRAM-based difference tests. Vertical and horizontal error bars represent standard errors of GWAS effect sizes in EAS and EUR, respectively. **D** The QQ-plots of $p$-values for all SNPs and SNPs in the top quantile of four continuous-valued annotations.

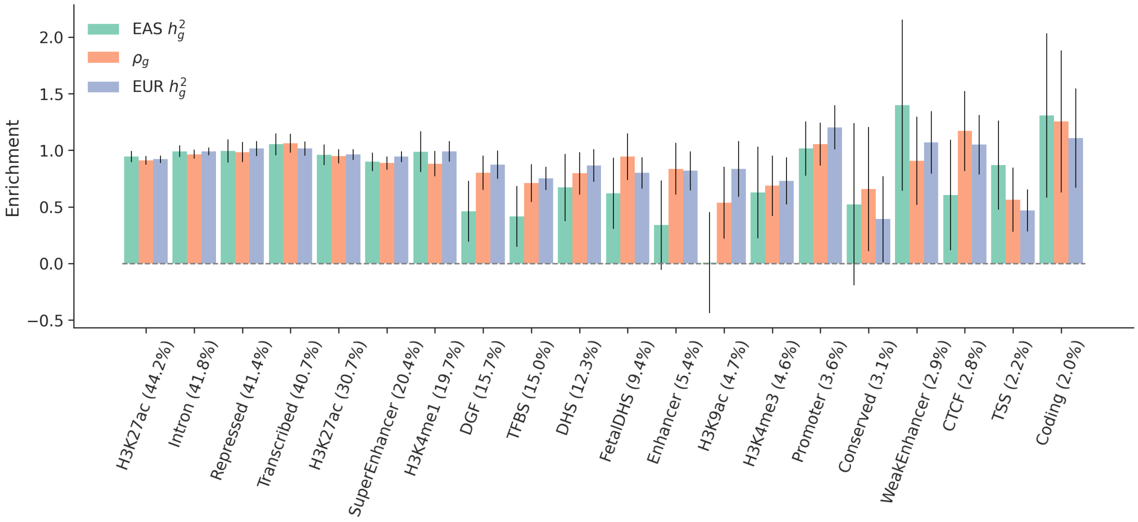


Figure S20: The S-LDXR estimates of the enrichment/depletion of trans-ancestry genetic correlation ($\rho_g$) and population-specific heritability across 20 binary functional annotations in Baseline-LD-X model for BMI. The functional annotations are sorted by proportion of SNPs (displayed in parentheses)

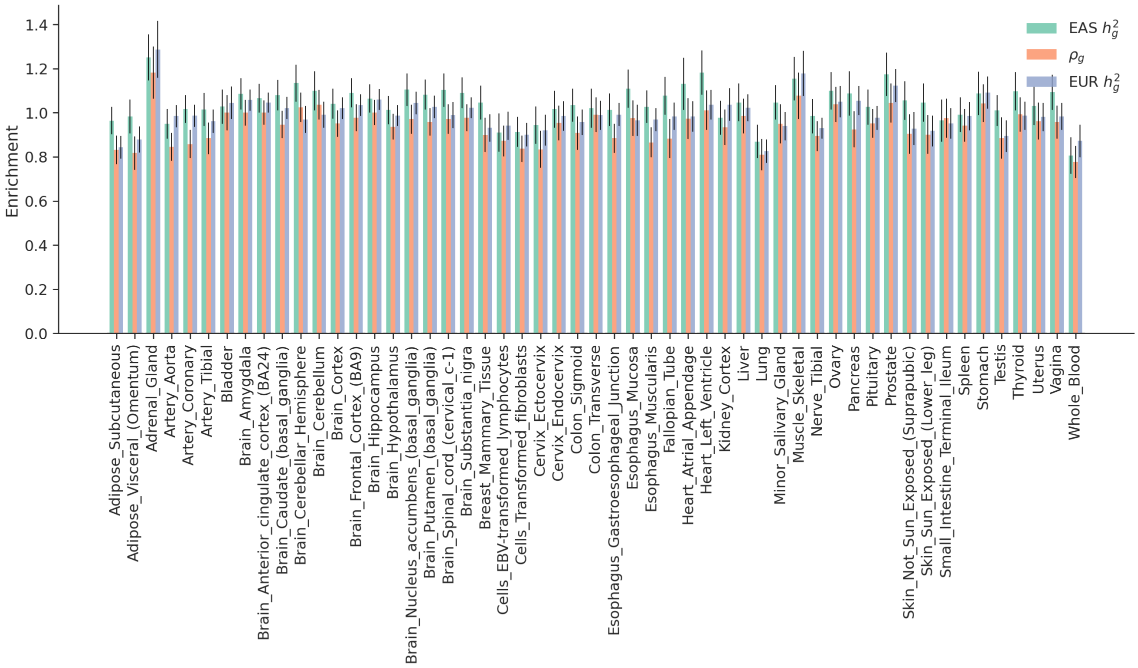


Figure S21: The S-LDXR estimates of the enrichment/depletion of trans-ancestry genetic correlation ($\rho_g$) and population-specific heritability across 53 tissue/cell type specifically expressed genes (SEG) annotations for BMI.

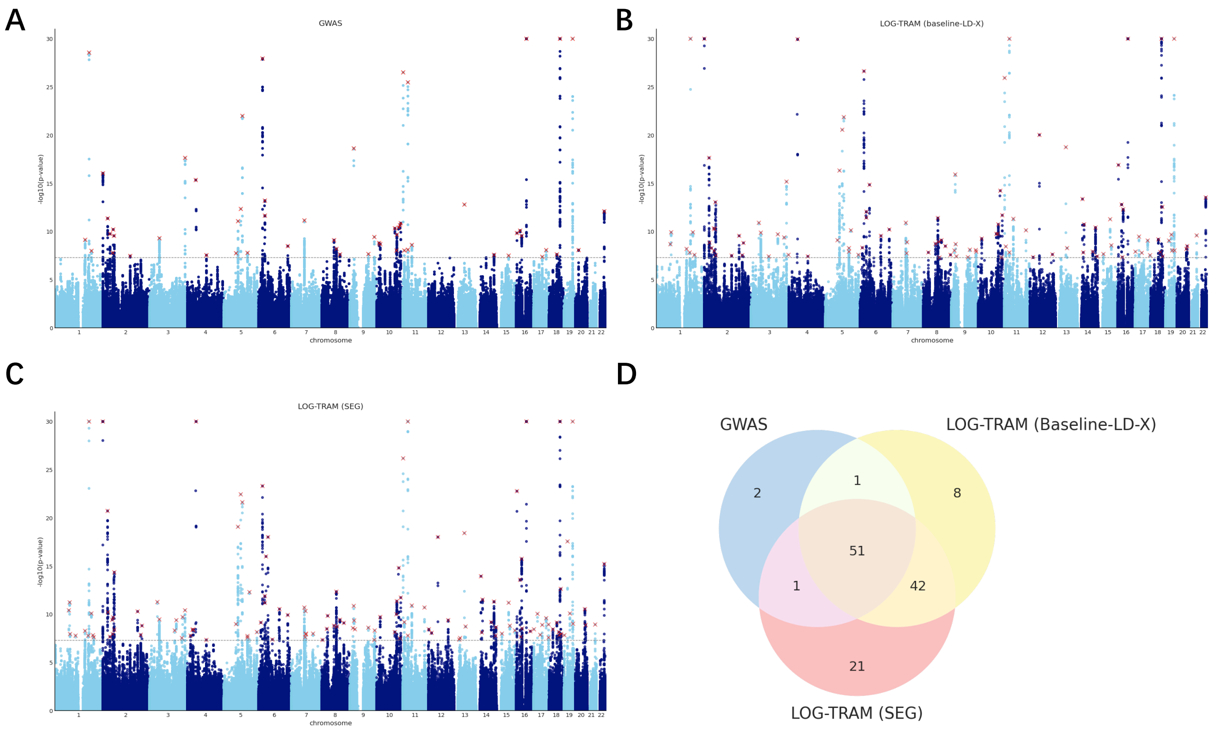


Figure S22: Meta-analysis results of the BMI using annotations as local regions. **A**: Manhattan plot of the BBJ GWAS data, where 55 independent loci were identified. **B**: Manhattan plot of BBJ meta-analysis result obtained by the integrative analysis of LOG-TRAM with 20 binary functional annotations in Baseline-LD-X model as local regions. LOG-TRAM identified 102 independent lead SNPs, of which 50 were not identified in the original BBJ GWAS. Lead SNPs identified by both the LOG-TRAM and GWAS are marked by crosses. **C**: Manhattan plot of BBJ meta-analysis result obtained by the integrative analysis of LOG-TRAM with 53 SEG annotations as local regions. LOG-TRAM identified 115 independent lead SNPs, of which 63 were not identified in the original BBJ GWAS. **D**: Venn diagram comparing the loci found by GWAS, LOG-TRAM with Baseline-LD-X model and LOG-TRAM with SEG model.

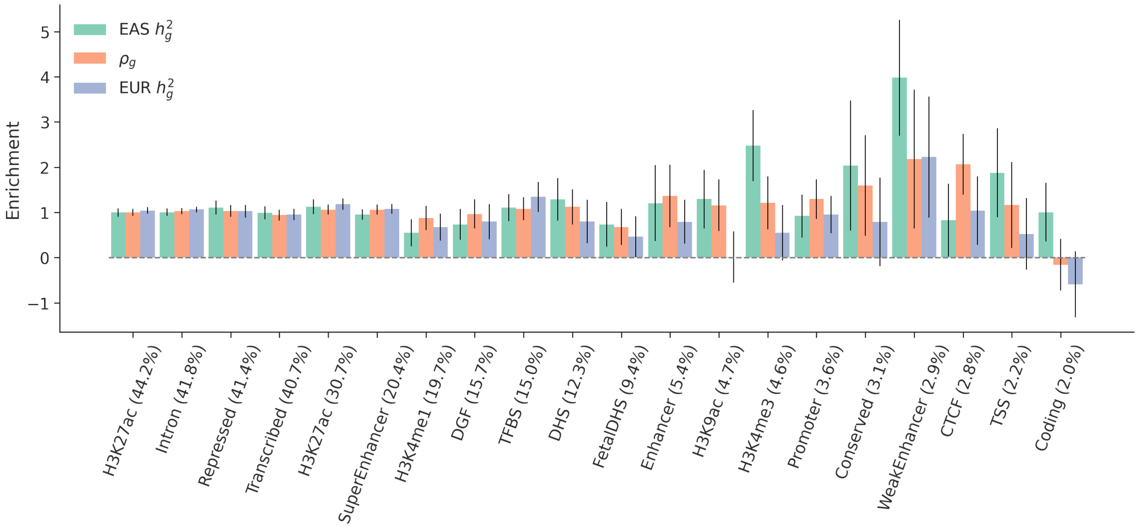


Figure S23: The S-LDXR obtained estimates of the enrichment/depletion of trans-ancestry genetic correlation ($\rho_g$) and population-specific heritability across 20 binary functional annotations in Baseline-LD-X model for T2D. The functional annotations are sorted by proportion of SNPs (displayed in parentheses)

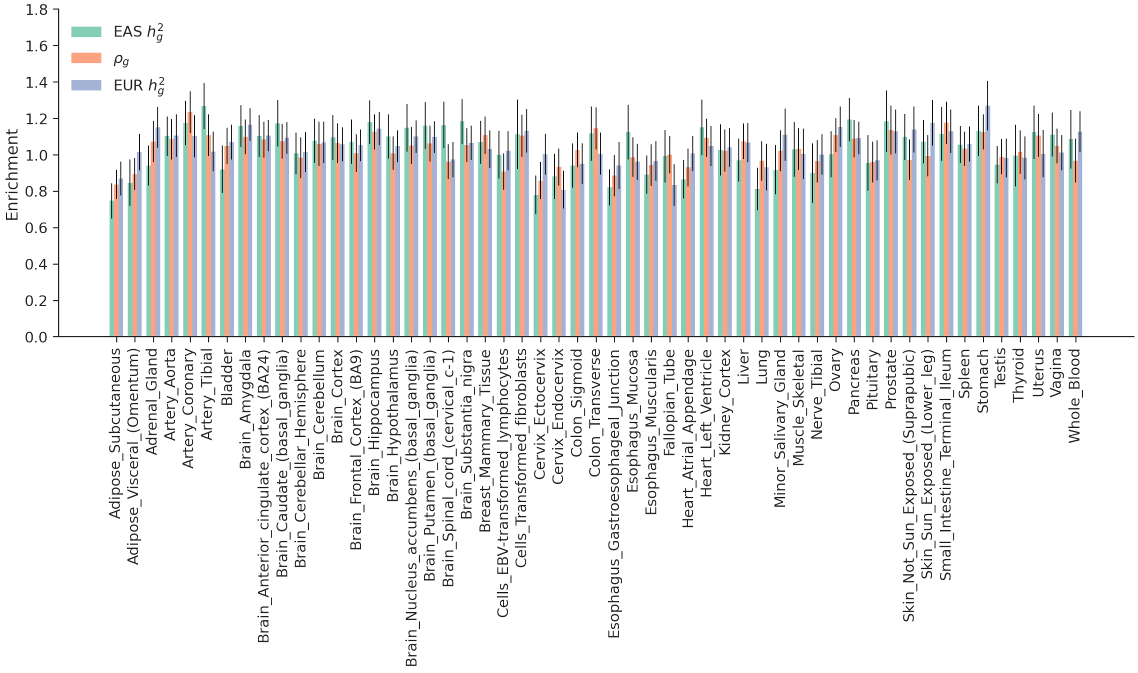


Figure S24: The S-LDXR estimates of the enrichment/depletion of trans-ancestry genetic correlation ($\rho_g$) and population-specific heritability across 53 SEG annotations for T2D.

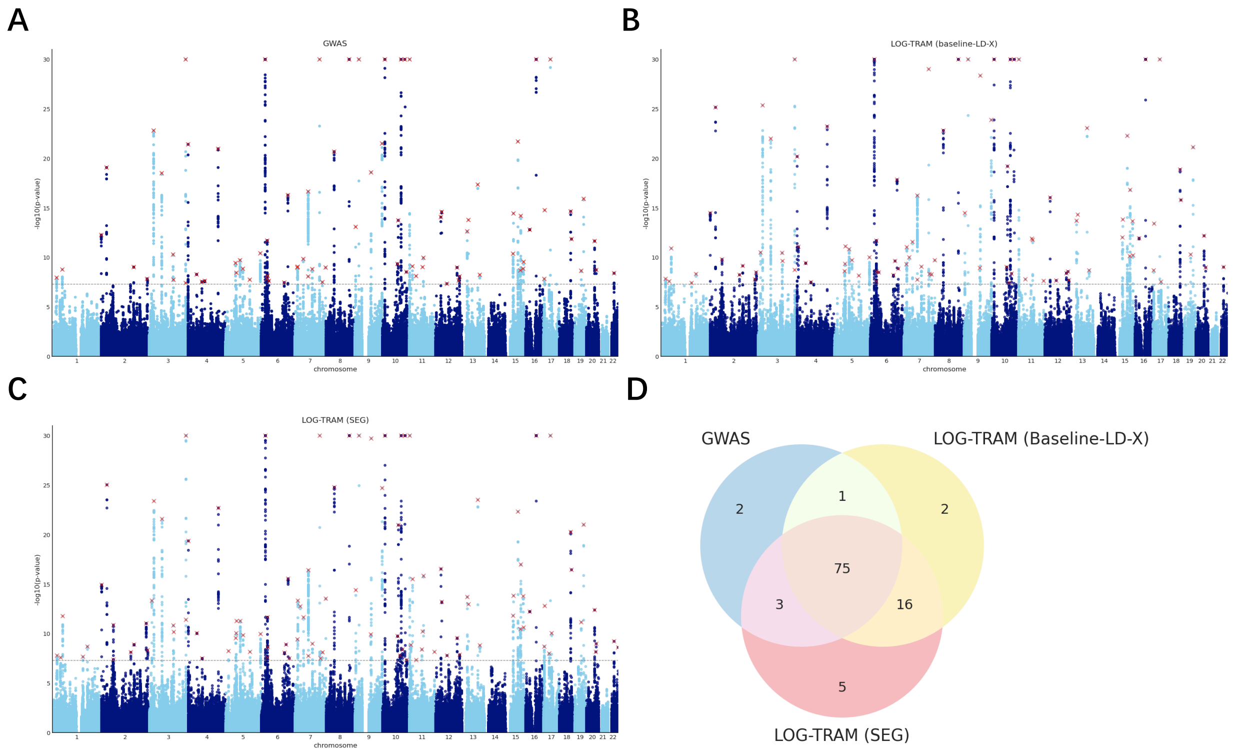


Figure S25: Meta-analysis results of the T2D using annotations as local regions. **A**: Manhattan plot of the BBJ GWAS data, where 81 independent loci were identified. **B**: Manhattan plot of BBJ meta-analysis result obtained by the integrative analysis of LOG-TRAM with 20 binary functional annotations in Baseline-LD-X model as local regions. LOG-TRAM identified 94 independent lead SNPs, of which 18 were not identified in the original BBJ GWAS. Lead SNPs identified by both LOG-TRAM and GWAS are marked by a cross. **C**: Manhattan plot of BBJ meta-analysis result obtained by the integrative analysis of LOG-TRAM with 53 SEG annotations as local regions. LOG-TRAM identified 99 independent lead SNPs, of which 21 were not identified in the original BBJ GWAS. **D**: Venn diagram comparing the loci found by GWAS, LOG-TRAM with Baseline-LD-X model and LOG-TRAM with SEG model.

### 2 Supplementary Tables

| Trait name | Full name | sample size (case+control) | paper/data link |
|---|---|---|---|
| AD-EAS | Atopic dermatitis | 2385+209651 | https://www.nature.com/articles/s41588-020-0640-3 |
| AD-EUR | Atopic dermatitis | 9321+351820 | http://www.nealelab.is/uk-biobank/ |
| AF-EAS | Atrial Fibrillation | 8180+28612 | https://www.nature.com/articles/ng.3842 |
| AF-EUR | Atrial Fibrillation | 10986+233904 | https://www.gov.uk/government/publications/atrial-fibrillation-prevalence-estimates-for-local-populations |
| Asthma-EAS | Asthma | 8216+201592 | https://www.nature.com/articles/s41588-020-0640-3 |
| Asthma-EUR | Asthma | 41633+318894 | http://www.nealelab.is/uk-biobank/ |
| BMI-EAS | Body Mass Index | 158284 | https://www.nature.com/articles/ng.3951 |
| BMI-EUR | Body Mass Index | 457824 | https://www.nature.com/articles/s41588-018-0144-6 |
| BUN-EAS | Blood urea nitrogen | 139818 | https://www.nature.com/articles/s41588-018-0047-6 |
| BUN-EUR | Blood urea nitrogen | 480698 | https://www.ncbi.nlm.nih.gov/pmc/articles/PMC6377354/ |
| COPD-EAS | Chronic obstructive pulmonary disease | 3315+201592 | https://www.nature.com/articles/s41588-020-0640-3 |
| COPD-EUR | Chronic obstructive pulmonary disease-ICD90(J44) | 1531+359663 | http://www.nealelab.is/uk-biobank/ |
| CoCa-EAS | Colorectal cancer | 7062+195745 | https://www.nature.com/articles/s41588-020-0640-3 |
| CoCa-EUR | malignant neoplasm of colon-IDC10(C18) | 2226+358968 | http://www.nealelab.is/uk-biobank/ |
| Eosino-EAS | Eosinophil count | 62076 | http://jenger.riken.jp/en/result |
| Eosino-EUR | Eosinophil count | 349856 | http://www.nealelab.is/uk-biobank/ |
| FG-EAS | Fasting glucose | 31669 | https://magicinvestigators.org/ |
| FG-EUR | Fasting glucose | 159940 | https://magicinvestigators.org/ |
| FI-EAS | Fasting insulin | 26691 | https://magicinvestigators.org/ |
| FI-EUR | Fasting insulin | 159940 | https://magicinvestigators.org/ |
| Glaucoma-EAS | Glaucoma | 5761+206692 | https://www.nature.com/articles/s41588-020-0640-3 |
| Glaucoma-EUR | Glaucoma | 6674+355433 | http://www.nealelab.is/uk-biobank/ |
| HDL-EAS | High-density-lipoprotein cholesterol | 70657 | https://www.nature.com/articles/s41588-018-0047-6 |
| HDL-EUR | High-density-lipoprotein cholesterol | 188577 | https://www.nature.com/articles/ng.2797 |
| HbA1c-EAS | Hemoglobin A1c | 42790 | http://jenger.riken.jp/en/result |
| HbA1c-EUR | Hemoglobin A1c | 159940 | https://magicinvestigators.org/ |
| Height-EAS | height | 159095 | https://www.nature.com/articles/s41467-019-12276-5 |
| Height-EUR | height | 458303 | https://www.nature.com/articles/s41588-018-0144-6 |
| LDL-EAS | Low-density-lipoprotein cholesterol | 72866 | https://www.nature.com/articles/s41588-018-0047-6 |
| LDL-EUR | Low-density-lipoprotein cholesterol | 188577 | https://www.nature.com/articles/ng.2797 |
| Lym-EAS | Lymphocyte count | 62076 | http://jenger.riken.jp/en/result |
| Lym-EUR | Lymphocyte count | 349856 | http://www.nealelab.is/uk-biobank/ |
| MCH-EAS | Mean corpuscular hemoglobin concentration | 108054 | https://www.nature.com/articles/s41588-018-0047-6 |
| MCH-EUR | Mean corpuscular hemoglobin | 132224 | https://www.cell.com/cell/references/S0092-8674(16)31463-5 |
| MCHC-EAS | Mean corpuscular hemoglobin concentration | 108728 | https://www.nature.com/articles/s41588-018-0047-6 |
| MCHC-EUR | Mean corpuscular hemoglobin concentration | 132586 | https://www.cell.com/cell/references/S0092-8674(16)31463-5 |
| MCV-EAS | Mean corpuscular volume | 108256 | https://www.nature.com/articles/s41588-018-0047-6 |
| MCV-EUR | Mean corpuscular volume | 132353 | https://www.cell.com/cell/references/S0092-8674(16)31463-5 |
| Mono-EAS | Monocyte count | 62076 | http://jenger.riken.jp/en/result |
| Mono-EAS | Monocyte count | 62076 | http://jenger.riken.jp/en/result |
| Mono-EUR | Monocyte count | 349856 | http://www.nealelab.is/uk-biobank/ |
| Mono-EUR | Monocyte count | 349856 | http://www.nealelab.is/uk-biobank/ |
| Plt-EAS | Platelet count | 108208 | http://jenger.riken.jp/en/result |
| Plt-EUR | Platelet count | 350474 | http://www.nealelab.is/uk-biobank/ |
| PrCa-EAS | Prostate cancer | 5408+103939 | https://www.nature.com/articles/s41588-020-0640-3 |
| PrCa-EUR | Prostate cancer | 6321+160699 | http://www.nealelab.is/uk-biobank/ |
| RBC-EAS | Red blood cell count | 108794 | http://jenger.riken.jp/en/result |
| RBC-EUR | Red blood cell count | 350475 | http://www.nealelab.is/uk-biobank/ |
| SBP-EAS | Systolic blood pressure | 136597 | http://jenger.riken.jp/en/result |
| SBP-EUR | Systolic blood pressure | 422771 | http://www.nealelab.is/uk-biobank/ |
| T2D-EAS | Type 2 Diabetes | 36614+155150 | https://www.nature.com/articles/s41588-020-0640-3 |
| T2D-EUR | Type 2 Diabetes | 459324 | https://www.nature.com/articles/s41588-018-0144-6 |
| TC-EAS | Total cholesterol | 128305 | https://www.nature.com/articles/s41588-018-0047-6 |
| TC-EUR | Total cholesterol | 188577 | https://www.nature.com/articles/ng.2797 |
| TG-EAS | Triglyceride | 105597 | https://www.nature.com/articles/s41588-018-0047-6 |
| TG-EUR | Triglyceride | 188577 | https://www.nature.com/articles/ng.2797 |
| UF-EAS | Uterine fibroids | 5954+95010 | https://www.nature.com/articles/s41588-020-0640-3 |
| UF-EUR | Uterine fibroids | 5514+188639 | http://www.nealelab.is/uk-biobank/ |
| eGFR-EAS | Estimated glomerular filtration rate | 143658 | https://www.nature.com/articles/s41588-018-0047-6 |
| eGFR-EUR | Estimated glomerular filtration rate | 480698 | https://www.ncbi.nlm.nih.gov/pmc/articles/PMC6377354/ |

Table S1: Sources of 29 traits from EUR and EAS.

| Trait | GWAS (EAS) | LOG-TRAM (EAS) | GWAS (EUR) | LOG-TRAM (EUR) |
|---|---|---|---|---|
| AD | 212036.0 | 336578.99 | 361141.0 | 545100.43 |
| AF | 36792.0 | 44807.55 | 244065.0 | 304184.41 |
| Asthma | 209808.0 | 316055.09 | 360527.0 | 385759.11 |
| BMI | 158284.0 | 217512.52 | 457824.0 | 417499.64 |
| BUN | 139818.0 | 169815.0 | 240985.0 | 278999.82 |
| COPD | 204907.0 | 298551.59 | 361194.0 | 556956.65 |
| CoCa | 202807.0 | 268546.72 | 361194.0 | 596956.58 |
| Eosino | 62076.0 | 118482.84 | 349856.0 | 355655.76 |
| FG | 31666.0 | 82424.06 | 166355.0 | 173884.53 |
| FI | 26690.0 | 76470.54 | 124119.0 | 140880.51 |
| Glaucoma | 212453.0 | 267859.4 | 361194.0 | 606870.88 |
| HDL | 70657.0 | 77132.53 | 94280.0 | 81387.99 |
| HbA1c | 42790.0 | 69718.05 | 128610.0 | 146733.36 |
| LDL | 72866.0 | 103591.71 | 89858.0 | 81217.14 |
| Lym | 62076.0 | 119853.52 | 349856.0 | 328102.8 |
| MCH | 108054.0 | 132064.76 | 132224.0 | 142997.66 |
| MCHC | 108728.0 | 144459.9 | 132586.0 | 155073.74 |
| MCV | 108256.0 | 130994.28 | 132353.0 | 137945.08 |
| Mono | 62076.0 | 103975.47 | 349856.0 | 286454.08 |
| Plt | 108208.0 | 157902.19 | 350474.0 | 299383.67 |
| PrCa | 109347.0 | 134173.38 | 167020.0 | 224174.66 |
| RBC | 108794.0 | 159145.66 | 350475.0 | 310820.46 |
| SBP | 136597.0 | 231835.59 | 422771.0 | 374903.24 |
| T2D | 210865.0 | 229534.82 | 459324.0 | 522750.9 |
| TC | 128305.0 | 173209.07 | 94564.0 | 96226.11 |
| TG | 105597.0 | 123625.95 | 90982.0 | 96870.7 |
| UF | 100964.0 | 171576.43 | 194153.0 | 342573.31 |
| eGFR | 143658.0 | 190291.09 | 560658.0 | 641597.88 |
| height | 159095.0 | 160762.66 | 458303.0 | 299754.07 |

Table S2: The effective sample size of LOG-TRAM for 29 traits from EUR and EAS.

| Trait | GWAS | LOG-TRAM | MTAG | MAMA | RE2 |
|---|---|---|---|---|---|
| height | 1.2235 (0.0233) | 1.0499 (0.0231) | 0.9478 (0.0235) | 1.1495 (0.0306) | 1.9237 (0.0493) |
| BMI | 1.0754 (0.013) | 1.0447 (0.0152) | 0.8547 (0.0139) | 1.0793 (0.0167) | 1.3503 (0.0235) |
| T2D | 1.0395 (0.013) | 1.0097 (0.0135) | 0.9729 (0.0135) | 0.9968 (0.0138) | 1.0047 (0.0136) |
| MCH | 1.0554 (0.0116) | 1.013 (0.0128) | 0.7697 (0.0149) | 0.9619 (0.0195) | 1.2423 (0.0268) |
| SBP | 1.0515 (0.0106) | 1.0112 (0.0119) | 0.8247 (0.0113) | 1.0276 (0.0141) | 1.2764 (0.0203) |
| eGFR | 1.0885 (0.0106) | 1.0185 (0.0122) | 0.9094 (0.0122) | 0.9902 (0.0129) | 1.0556 (0.0155) |
| BUN | 1.0577 (0.0093) | 1.0068 (0.0101) | 0.9665 (0.0105) | 0.9082 (0.0099) | 0.9966 (0.0109) |
| MCHC | 1.0473 (0.0097) | 1.002 (0.0097) | 0.9495 (0.0092) | 1.0062 (0.0095) | 0.9969 (0.0112) |
| MCV | 1.0631 (0.0123) | 1.0026 (0.0126) | 0.9155 (0.0128) | 0.9901 (0.0138) | 1.0775 (0.0175) |
| AF | 1.2106 (0.01) | 1.0041 (0.0094) | 0.9687 (0.0089) | 0.8255 (0.0077) | 0.9594 (0.0086) |
| Asthma | 1.007 (0.0084) | 1.0196 (0.0097) | 0.9229 (0.0096) | 1.0441 (0.0109) | 1.0134 (0.0127) |
| UF | 0.9994 (0.0076) | 1.0089 (0.0084) | 0.9954 (0.0081) | 1.0096 (0.0082) | 0.9259 (0.008) |
| Glaucoma | 1.0049 (0.008) | 1.0173 (0.0083) | 0.9973 (0.0077) | 0.9989 (0.0076) | 0.9259 (0.0067) |
| CoCa | 1.0004 (0.0073) | 1.0076 (0.0083) | 0.9954 (0.0082) | 0.9957 (0.0084) | 0.9302 (0.0077) |
| COPD | 0.992 (0.0083) | 1.019 (0.0089) | 0.9952 (0.008) | 1.008 (0.008) | 0.9169 (0.0073) |
| PrCa | 0.9606 (0.0102) | 1.0047 (0.0104) | 0.9639 (0.0102) | 0.979 (0.0102) | 0.9334 (0.0103) |
| AD | 0.9775 (0.008) | 1.0133 (0.0092) | 0.9944 (0.0085) | 1.0157 (0.0087) | 0.9334 (0.0082) |

Table S3: The estimated LDSC intercepts for traits from EAS.

| Trait | GWAS | LOG-TRAM | MTAG | MAMA | RE2 |
|---|---|---|---|---|---|
| height | 1.7784 (0.0505) | 1.0907 (0.0401) | 1.2156 (0.0434) | 1.2509 (0.0348) | 1.9237 (0.0493) |
| BMI | 1.2121 (0.0197) | 1.02 (0.0185) | 1.127 (0.0201) | 1.1067 (0.0161) | 1.3503 (0.0235) |
| T2D | 1.0463 (0.0101) | 1.0209 (0.0122) | 0.956 (0.013) | 1.0413 (0.0165) | 1.0047 (0.0136) |
| MCH | 1.28 (0.0286) | 1.0348 (0.0267) | 0.9615 (0.025) | 1.0098 (0.0205) | 1.2423 (0.0268) |
| SBP | 1.2113 (0.0179) | 1.0339 (0.017) | 1.044 (0.0168) | 1.0421 (0.0131) | 1.2764 (0.0203) |
| eGFR | 1.0279 (0.012) | 1.0155 (0.0144) | 1.022 (0.014) | 1.0229 (0.0127) | 1.0556 (0.0155) |
| BUN | 1.0222 (0.0089) | 1.0064 (0.0114) | 0.9765 (0.0113) | 0.938 (0.0109) | 0.9966 (0.0109) |
| MCHC | 1.0326 (0.0092) | 1.0212 (0.0116) | 0.9975 (0.0109) | 1.0251 (0.0105) | 0.9969 (0.0112) |
| MCV | 1.0976 (0.0164) | 1.0221 (0.0174) | 0.9583 (0.0163) | 1.0386 (0.0162) | 1.0775 (0.0175) |
| AF | 1.0063 (0.0082) | 1.0098 (0.0102) | 0.9831 (0.0101) | 0.9766 (0.0101) | 0.9594 (0.0086) |
| Asthma | 1.0563 (0.011) | 1.0162 (0.0126) | 1.0241 (0.0126) | 1.057 (0.0115) | 1.0134 (0.0127) |
| UF | 1.0054 (0.0065) | 1.0115 (0.0088) | 0.9902 (0.0089) | 1.0091 (0.0088) | 0.9259 (0.008) |
| Glaucoma | 0.9997 (0.006) | 1.0187 (0.0077) | 0.9906 (0.0073) | 1.0078 (0.0081) | 0.9259 (0.0067) |
| CoCa | 1.0058 (0.0062) | 1.0117 (0.008) | 0.9948 (0.0074) | 1.0093 (0.0087) | 0.9302 (0.0077) |
| COPD | 0.9968 (0.0063) | 1.0187 (0.008) | 0.9956 (0.0076) | 1.0208 (0.0079) | 0.9169 (0.0073) |
| PrCa | 1.0213 (0.0074) | 1.0166 (0.0094) | 0.9699 (0.0098) | 0.9942 (0.0116) | 0.9334 (0.0103) |
| AD | 1.0169 (0.007) | 1.0181 (0.0088) | 0.9938 (0.0083) | 1.0336 (0.0094) | 0.9334 (0.0082) |

Table S4: The estimated LDSC intercepts for traits from EUR.
